# Rust Fungus Red Data List and Census Catalogue for Wales

**DOI:** 10.1101/2024.03.13.582811

**Authors:** Ray G. Woods, R. Nigel Stringer, Debbie A. Evans, Arthur O. Chater

**Affiliations:** Aberystwyth University; none

## Abstract

The rust fungi are a group of specialised plant pathogens. Conserving them seems to fly in the face of reason. Yet as our population grows and food supplies become more precarious, controlling pathogens of crop plants becomes more imperative. Breeding resistance genes into such plants has proved to be the most cost effective solution. Such resistance genes evolve only in plants challenged by pathogens. We hope this report will assist in prioritising the conservation of natural ecosystems and traditional agro-ecosystems that are likely to be the richest sources of resistance genes. Despite its small size (11% of mainland Britain) Wales has supported 225 rust fungi taxa (including 199 species) representing 78% of the total British mainland rust species. For the first time using widely accepted international criteria and data collected from a number of mycologists and institutions, a Welsh regional threat status is offered for all native Welsh rust taxa. The results are compared with other published Red Lists for Wales. Information is also supplied in the form of a census catalogue, detailing the rust taxa recorded from each of the 13 Welsh vice-counties.

Of the 225 rust taxa so far recorded from Wales 7 are probably extinct (3% of the total), and 39 (18%) are threatened with extinction. Of this latter total 13 taxa (6%) are considered to be Critically Endangered, 15 (7%) to be Endangered and 13 (6%) to be Vulnerable. A further 20 taxa (9%) are Near Threatened, whilst 15 taxa (7%) lacked sufficient data to permit evaluation.

Just over a third of Welsh rust fungi taxa require some form of action, either to better understand their status, or reverse known ongoing declines. This is a similar figure to those presented elsewhere for bryophytes and lichens but higher than the 19% of vascular plants requiring action.

## Foreword

Wales has been particularly fortunate in having a dedicated band of mycologists recording rust fungi. This report would not have been possible without the work of Nigel Stringer who has made innumerable records, checked specimens, enthused us all and has collated the records of those working in Wales. A small but dedicated group of largely amateur mycologists has covered large parts of the country. Debbie Evans has amassed many records from North Wales, especially in the vice-counties of Caernarvonshire and Anglesey and is now able to provide an excellent picture of rust fungi distribution in what was a relatively poorly surveyed area. Some of her records appear with those of others such as Richard Shattock and Charles Aron in the latter’s *Fungi of Northwest Wales* (2005), this being an invaluable compilation of records. Andrew Graham, Wendy McCarthy, John Bratton and Paul Reade have contributed some interesting records of late. Bruce Ing has collected regularly in NE Wales.

Tom Preece, Ray Woods and Graham Motley have worked in central and east Wales whilst Arthur Chater has made numerous collections from Cardiganshire and parts of the neighbouring vice-counties. Carmarthenshire has been covered in detail by Nigel Stringer (creating over 18,000 personal records), Richard Davies, Ian Morgan and others to produce a database of over 41,000 records from this one vice-county. Paul Smith has collected in both Monmouthshire and Glamorgan. Pembrokeshire has been surveyed by visiting mycologists and now has an active fungus recording group. Since the publication of Wilson and Henderson in 1966 this band of recorders has found as new to Britain and Ireland almost 250 previously unreported combinations of rust and host taxa and four species of rust new to Britain.

This work also draws on specimens held at Kew Gardens and in the National Museum and Galleries of Wales, Cardiff and on the records in the British Mycological Society’s Fungus Records Database of Britain and Ireland (FRDBI) and the Association of British Fungus Groups (ABFG) CATE2 database.

## 1 Introduction

The Kingdom of the Fungi is a diverse one. Hawksworth (2001) calculates that there are at least 1.5 million species worldwide. For the most part little action has been taken to directly conserve them. Yet we derive immense benefits directly and indirectly from fungi and occasionally come to grief due to their activities. Fungi could well be Wales’ most beneficial and under-used natural resource. Any attempt to develop Wales sustainably that ignores the fungi is doomed to failure. It is anticipated, for example, that inadequate biosecurity will shortly create a massive clean-up bill from the certain death of ash trees due to the failure to exclude or control ash die-back disease. Fungi provide us with some of the most widely employed pharmaceuticals. The search for new antibiotics might usefully be directed first towards the fungi from which so many of our most beneficial antibiotics have been obtained in the past.

Animals and Fungi diverged from a common ancestor and the Fungi are a sister kingdom to that of the Animals. We share many physiological functions with them making the treatment of the few fungal pathogenic diseases we suffer from often problematic. On the positive side some antibiotics that fungi have evolved to protect themselves work well in humans. With the development of techniques to identify fungi living within plants (endophytic fungi) has come the realisation that all plants interact with fungi and many more benefit than suffer from them. Endophytic fungi have developed complex interactions with bacteria. These relationships have yet to be exploited but offer huge potential for increasing crop yields and improvements in resistance to pests, diseases and changes in climate and may even allow crops to be taken from soils currently considered unsuitable for cultivation.

The rust fungi form a numerous and well-defined group called the Pucciniales (or Uredinales) within the Basidiomycota - the group that contains the mushrooms and toadstools. Of the 22,000 species described worldwide from the Basidiomycota (35% of all known fungi) a third (7,400 species) are rust fungi. Since all rust fungi are plant parasites, with a few occasionally causing devastating consequences to our food production, we ignore them at our peril. Some probably play a significant role in influencing the composition of vegetation. Zadoks (2005) for example demonstrated the important role that rust and other pathogenic fungi play in modifying the species-composition of salt marsh vegetation in the Netherlands. Plants introduced to new continents without their rust fungi such as bramble *Rubus fruticosus* agg. in parts of Australia and mouse-ear hawkweed *Pilosella officinalis* in New Zealand have rapidly grown to pest proportions not seen in their native, rust-rich home territories. In Britain field trials have commenced to evaluate the efficacy of the non-native rust *Puccinia komarovii* var. *glanduliferae* as a possible control agent for the introduced Himalayan balsam *Impatiens glandulifera*.

Several rust species threaten the success of our food and fibre production. Our recent aggressive expansion of agriculture and forestry, often employing monocultures of genetically identical organisms, has left us woefully exposed to the depredations of some fungi. Such production methods fly in the face of the basic principles of epidemiology. Black stem rust *Puccinia graminis*, for example, regularly devastated crops of wheat (Ingram and Robertson 1999). Yet for almost the last 50 years this rust has ceased to be a problem due to resistance genes introduced into wheat from rye. In 1999 the rust overcame this resistance in Uganda and, having spread out of Africa and known as Ug99, now threatens wheat crops around the world.

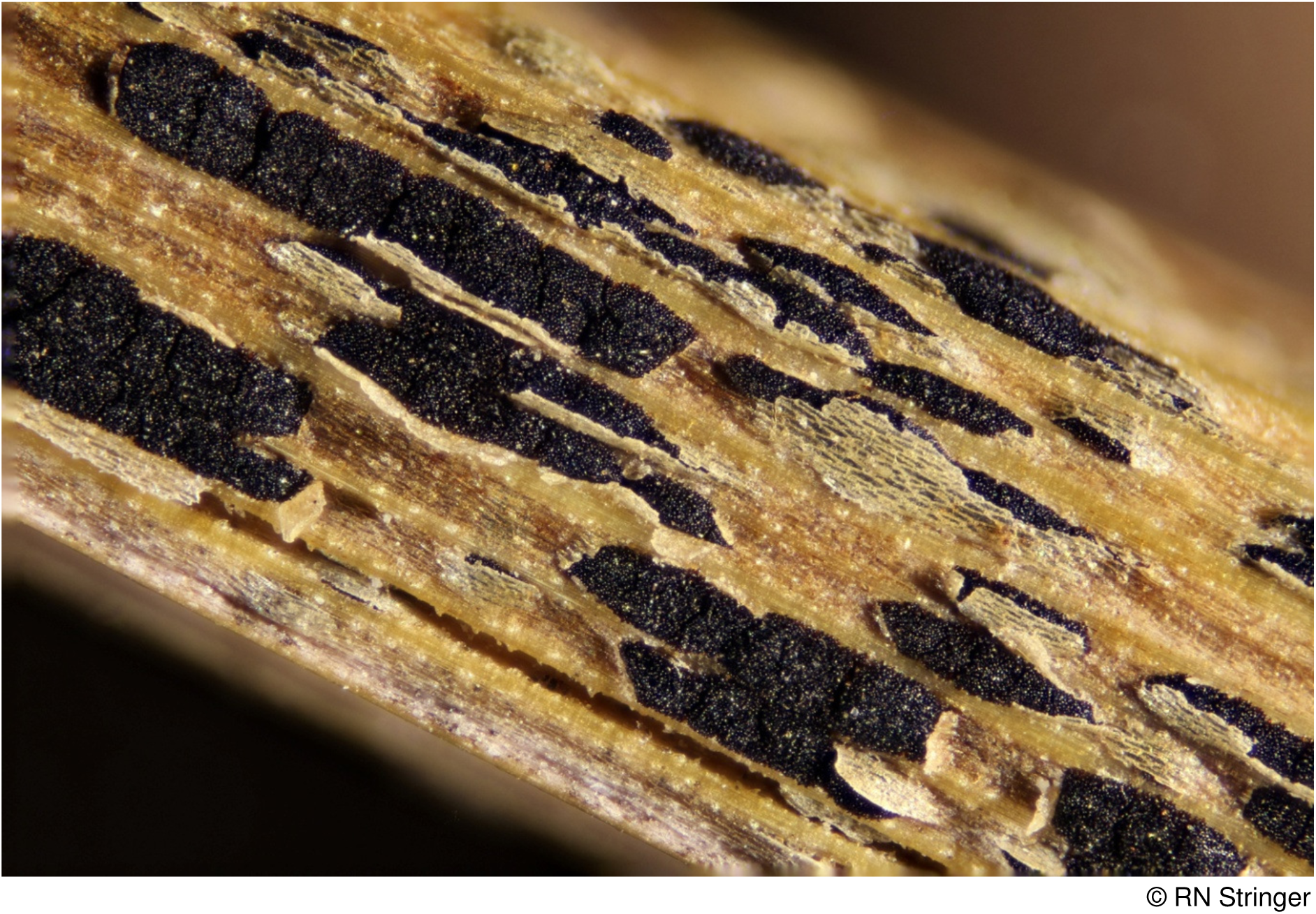
Black stem-rust telia on cock’s-foot grass

Having erroneously considered the battle won, wheat breeding programmes had been wound down. The search is now on for new resistance genes in wild and cultivated plants. During the course of the collection of data for this RDL we have seen an increase in abundance of black stem-rust on ryegrass *Lolium perenne* leys and barberry *Berberis vulgaris* in Wales, probably attributable to particularly warm summers. This grass is at the centre of livestock-production in Wales. Unless additional resistance can be bred, in a warming climate we could see this rust causing economic losses. Resistance genes to many rust fungi may already be present in the wild plant populations but they are unlikely to persist in plants if those with the resistance are not regularly selected for by the challenge of infection. Lose the rust fungi and we may lose the genes for resistance.

The large scale shipping of plants and plant-derived commodities around the world further increases the chances of importing undesirable fungi. Following concerns over the importation of ash disease the Government has set up an independent task force on tree health and plant biosecurity. One of their recommendations, that a risk register be established, has been implemented (see http://secure.fera.defra.gov.uk/phiw/riskRegister). This risk register includes several rust fungi.

The development of woody species for biomass production has stimulated much new work on the impact of rust species on willow clones and the search for resistance to these fungi. It can cost up to £200 million pounds and take more than ten years to bring a new crop protection product to market (http://www.rothamsted.ac.uk/intopractice/FungicideResistanceReport.pdf). It is easy to see why resistance genes bred into a crop plant can be a very cost-effective alternative option. Biocontrol of rust fungi in crop plants by employing hyperparasitic fungi, bacteria or mycoviruses offers considerable promise (Moricca *et al*. 2006). By conserving rare wild rust fungi we may inadvertently be conserving hyperparasitic organisms that are potentially effective biocontrol agents of those rust fungi that damage crop plants. There is a distinct probability that the rarer the rust taxon the more likely it is to support debilitating hyperparasites. Their presence may be keeping the wild rust in check.

Ingram (1999) in his presidential address to the British Society for Plant Pathology in 1998 noted that plant pathogens are a significant component of natural ecosystems and are essential to the evolutionary selection of new disease resistance factors in the wild relatives of crops. He also notes that the conservation of the plant diversity of natural ecosystems and traditional agro-ecosystems is the greatest challenge of our age.

As we place ever more pressure on land, the area of species-rich semi-natural habitats declines in Wales. For example it is believed that 98% of dry lowland flower-rich grassland has been lost in the last 50 years (Stevens *et al*. 2010) mostly to agricultural intensification. With a decline in available host plants there will have been an associated decline in the rust fungi. In many parts of Wales the few remaining examples are now subject to nutrient enrichment from emissions of ammonia from ever-more intensive livestock and poultry rearing systems.

Even such apparently unspoiled habitats as sub-montane cliffs have not been immune, receiving acidifying atmospheric pollutants and are now being subject to a rapidly changing climate. Research at the Royal Botanic Gardens, Edinburgh is investigating possible changes to rust species infecting arctic-alpine willow species. The greater fragmentation of habitats may also reduce the ability of those rusts that require two completely contrasting host species in order to compete their life cycle to survive. Some rust fungi may select particular genotypes of their host species so highlighting genetic diversity in the host plants that was previously unsuspected Helfer (1993) opened a debate on the issue of conserving rust fungi in a paper entitled “Rust Fungi - A Conservationist’s Dilemma” by noting that 52 out of 102 vascular plant species considered rare or endangered in the British Isles supported rust species and asked whether the rust fungi should enjoy a similar level of protection to their hosts. A lack of information precluded a definitive answer. Helfer *et al*. (2011) were participants in a European Uredinales Initiative (EURED) that called for a renewed pan-European effort to better understand the taxonomy, biogeography and ecology of rusts for many of the reasons given above. It is hoped this Red List will go a little way towards answering some of these questions.

Whilst Sites of Special Scientific Interest and National Nature Reserves provide refugia for potential host plants, fungi have rarely featured in their selection nor have they been taken into account in framing management strategies. This RDL is intended to help with the selection and management of sites with a view to ensuring the conservation of a fuller spectrum of the living world. It is also hoped that the Welsh Government, in its aspiration to develop Wales sustainably, will now be able to take into account this group of fungi.

## 2 Implementation of the Red Data List

The aim of this report is to assess the level of threat facing rust fungi in Wales so that priorities can be identified for conservation action. Any taxon that is threatened (Critically Endangered, Endangered, Vulnerable or Near Threatened) in Britain (Evans *et al*. 2006) should also be regarded as a priority for conservation in Wales, regardless of its threat status in Wales.

If a taxon is less threatened in Wales than it is in Britain (if it has a lower category of threat than in Britain or is even classified as Least Concern in Wales), the Welsh population must still be regarded as a critically important component of the British population and deserves full protection in Wales with appropriate conservation measures. Should the British population outside Wales continue to decline, the Welsh population will become increasingly important, again regardless of its status within Wales. Should the Welsh population begin to decline, the species will be regarded as even more threatened in Britain as a whole.

## 3 Taxonomic Coverage

All rust fungi species, subspecies and varieties occurring on native, archaeophyte and neophyte Welsh flowering plant and fern species have been considered. Some rusts are able to infect a wide range of hosts. The mallow rust *Puccinia malvacearum*, for example, infects all native British species in the mallow (Malvaceae) family, whilst the rust *Coleosporium tussilaginis* is recorded from over a dozen plant species in Wales in four different families. Whether such rust species have distinct races or pathotypes only able to infect some of these plant species is not fully known. *P. malvacearum*, a rust described in 1852 from Chile, first appeared in Europe in 1869 and in Britain in 1873 (Wilson and Henderson 1966). It quickly became a common rust of hollyhock *Alcea rosea* and common mallow *Malva sylvestris*. It was not until the 1990’s that it was first recorded on musk mallow *M. moschata*, with records quickly appearing from all over Britain. Had a new genotype of the rust appeared or was a new genotype of the host spreading across Britain? If such races definitely do exist then efforts should be made to identify and conserve them. An indication of the likelihood of such races or pathotypes being present in Wales is given in the text below. It is highly desirable that this infraspecific variation of both host and rust fungus is rapidly recognised if at all possible by DNA sequencing.

Rust fungi confined to cultivated non-native plants that have not become naturalised in Wales have been omitted from consideration as to conservation status but are listed in the Census Catalogue. Where rusts have been reported from more than one cultivar, the additional cultivars are reported in Appendix 1. It should be noted that the recording of cultivars has been patchy across Wales and this list is far from complete.

Species names for the rust fungi mostly follow those used in the Kew *Checklist of the British and Irish Basidiomycota* (Legon and Henrici 2005) including the latest online updates at www.BasidioChecklist.info. Occasionally where a very recent revision has been adopted *Index Fungorum* has been followed. The taxonomy of the host plants follows Stace (2010).

## 4 Geographic Coverage

This Red Data List covers the country of Wales, made up of the 13 Watsonian Vice-counties of Monmouthshire, Glamorgan, Breconshire, Radnorshire, Carmarthenshire, Pembrokeshire, Cardiganshire, Montgomeryshire, Merionethshire, Caernarvonshire, Denbighshire, Flintshire and Anglesey (Watson 1883). Recording effort varies considerably between vice-counties with Breconshire, Radnorshire, Carmarthenshire, Cardiganshire, Caernarvonshire and Anglesey being possibly the best covered whilst Monmouthshire, Glamorgan, Pembrokeshire, Denbighshire and Flintshire are the least well covered.

## 5 Date Classification of Records

A “recent” record is one made in 1964 or later, i.e. in the last 50 years.

## 6 Application of IUCN Criteria

The standard IUCN Red Data List Categories (IUCN 2001) are used with minor modifications to take account of the regional nature of this analysis:

Taxa extinct within Wales but extant in other parts of the world are classified as Regionally Extinct (RE). A taxon is RE when there is no reasonable doubt that the last individual in the region has died or a species has not been seen since 1964. In this report, taxa extinct in Britain as a whole are classified as EX, while those extinct in Wales but still present elsewhere in Great Britain are classified as RE. The list of extinctions for Wales therefore includes both EX and RE taxa.

Considerable guidance is given by IUCN (2003) regarding the application of standard IUCN criteria and categories (IUCN, 2001) to a region (defined as any sub-global geographically defined area, such as a continent, country, state, or province). Provided that the regional population being assessed is isolated by, for example, unsuitable habitat or a lack of susceptible host material from conspecific populations of the rust outside the region, the IUCN Red Data List Criteria (IUCN 2001) can be used without modification within any geographically defined area.

However, when the criteria are applied to part of a population defined by a geopolitical border, as in the case of Wales sharing a border with England, the threshold values listed under each criterion may be inappropriate because the unit being assessed is not the same as the whole population or subpopulation. As a result, the estimate of extinction risk may be inaccurate.

In order to take this into account, we need to ask whether the Welsh population experiences any significant immigration of viable propagules from England. If not (or it is unknown), there is no change in the IUCN category assessed from a Welsh perspective. If, however, it is known or appears likely that viable propagules are entering Wales from England, the Welsh IUCN category is downgraded by one level provided that the GB population is stable or increasing. If the British population is decreasing the Welsh IUCN category remains the same. The level of propagule immigration is, however, almost impossible to assess. The spores of some rust species such as the urediniospores of *Puccinia coronata* are known to travel on average greater distances than the spores of *P. sessilis* (Stringer *et al*. 2011).

However, an attempt has been made to determine how likely immigration is by a consideration of the proximity of threatened Welsh taxa to English populations.

The threat category of Welsh taxa where the entire population is close to English populations may therefore be downgraded by one category if the British population is Least Concern. If, however, the British population is threatened the Welsh IUCN category remains unaltered.

Historic records of rust fungi in Wales are sparse and reside in a few old collections in national institutions, whilst a few published accounts e.g. Vize (1882) provide tantalizing hints of a one-time much richer mycota. This lack of any detailed information concerning the long term changes in population sizes permits only a very limited number of IUCN criteria to be employed. For most species only presence/absence data is available and population sizes can at best only be estimated.

Fluctuations from year to year in the number of “mature individuals” also form part of the criteria. Whilst such fluctuations may threaten the existence of, for example, birds or mammals, many fungi, including rusts, appear to vary naturally in abundance from year to year as measured by their visible fruit bodies. Until more is understood of these variations they have not formed a significant element of the evaluation process.

Rusts also pose some unique problems in assessing threats to populations. Issues include the selection of a threat status for populations of native rust species that appear clearly threatened on wild hosts, but that have, of late, appeared on cultivated plants in the horticultural trade. Such problems are highlighted in the species notes below.

Any direct measure of decline based on records over different time periods is problematic due to a substantial increase in recording effort in recent years. Instead, indirect methods have had to be adopted using known loss of habitat and changes in abundance of host species. All rusts are obligate parasites and some require more than one host to complete their life cycle. Their success is therefore intimately linked to the success and conservation status of their host plant or plants. For a rust species confined to a red data listed host the threat to the rust has been taken to be no lower than that of its host. Vascular plant red listings follow Cheffings *et al*. (2005) and Dines (2008).

Following in part Evans *et al*. (2006) the criteria listed below have been employed.

### Extinct

A taxon is listed as Extinct (EX) when there have been no records from Britain for 50 years (i.e. since 1964) or Regionally Extinct (RE) if there have been no records from Wales since 1964.

### Critically Endangered

**D1**: Found in 1 hectad and there is evidence of decline or a clear threat to host or habitat is identified.

**D2**: Where the total population in Wales numbers fewer than 50 individuals*.

### Endangered

**B2**: Found in 2-5 hectads and there is evidence of decline or a clear threat to host or habitat is identified.

**D**: Where the population in Wales numbers fewer than 250 individuals*.

### Vulnerable

**B**: Found in 6-10 hectads in Wales and there is evidence of decline or a clear threat to its host or habitat is identified.

**D**: Where the population in Wales is small and estimated to number fewer than 1000 individuals*.

**D1**: Found in 1-5 hectads in Wales with no evidence of decline of host or habitat.

**D2**: Found in 5 or fewer locations within Wales (where a location is a “Wells” site i.e. a moveable 1km square) and as a result is rendered liable to extinction through human activities or stochastic events.

#### Near Threatened

Found in 6-10 hectads within Wales with no evidence of decline of host or habitat or 11-20 hectads with a threat to host or habitat identified.

#### Data Deficient

Where there is insufficient information to place a probably threatened species in one of the categories above.

* IUCN uses the number of “mature individuals” as a measure of population size. This may be straightforward for elephants, but for rusts it is often unclear how many individuals might be present. A “mature individual” has here been taken to be a single infected plant. For some plants such as extensive stands of rhizomatous species *e.g.* saw sedge (*Cladium mariscus*) it is not clear where one individual host plant ends and another begins. A single discrete stand has been taken to be an individual. Worryingly this may still be a poor measure of the likelihood of the survival of some rust populations. In Australia Burdon *et al*. (1995) showed that the severity of rust infection of meadowsweet *Filipendula ulmaria* was positively correlated with the host population size and that a minimum number of host plants (termed a “deme”) had to be present before infection occurred.

## 7 Explanation of the Wales Red Data List and Census Catalogue

### Lists of Rust and Host Species (Columns 1 and 2)

Where infraspecific taxa (including varieties and pathotypes) have been listed, that information has been recorded but we advise its use with caution. Where a host plant name is followed by (A) this indicates it is considered to be an archaeophyte (naturalised before AD 1500); by (N) a neophyte (naturalised post AD 1500) and by (H) a non-naturalised cultivated plant (including forestry and agricultural crops as well as ornamental plantings). (Cas) indicates a species of casual occurrence and is usually applied to short lived weed species of temporary habitats. These definitions largely follow Preston *et al*. (2002). Where more than one category is applied it indicates that the status of this species varies from place to place in Wales.

### Wales Red List Threat Status and Criteria (Column 3)

These follow the explanation offered in section 6 above. EX indicates a taxon extinct in Britain; RE extinct in Wales but not Britain; CR indicates Critically Endangered; EN Endangered; VU Vulnerable; NT Near Threatened; DD Data Deficient and NE Not Evaluated. This last criterion is largely reserved for rusts confined to cultivated plants. In addition, LC identifies taxa of Least Concern

#### Census Catalogue (Column 4)

Under the heading of “Vice-County” are listed the Welsh vice-counties from which each rust fungus has been recorded. Vice-county numbers in square brackets indicate that no record has been traced post 1964. The vice-county numbers equate to the vice-counties as follows (Watson 1883):-

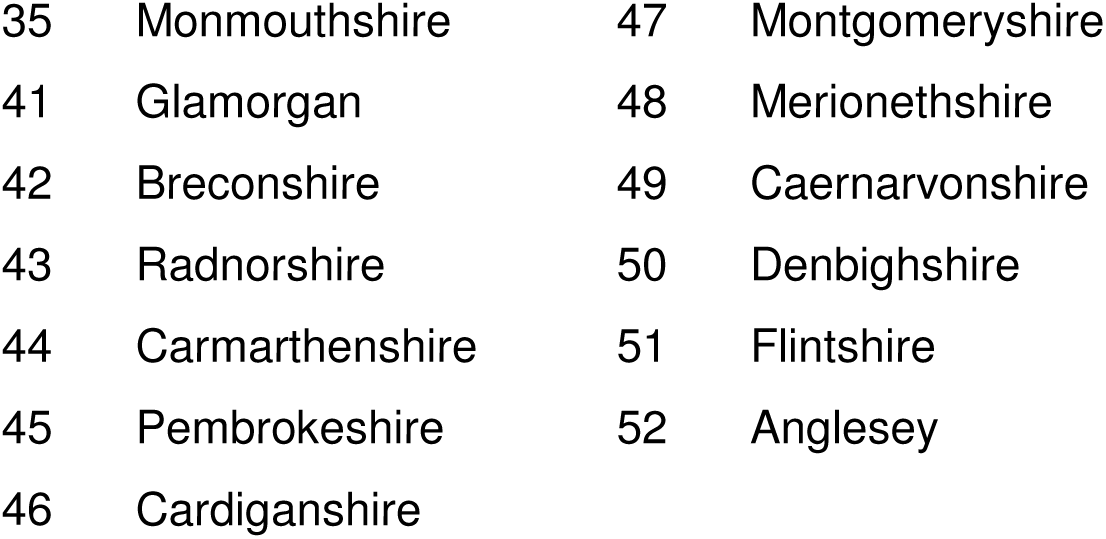

## 8 Analysis of Red Data List and Census catalogue

### 8.1 Red Data List

Presenting the results of this evaluation requires some initial explanation. A separation has been made in places between species and taxa. Species numbers refer as one would expect just to the species. Taxa numbers include in addition subspecies, varieties and pathotypes. Note also that some rust species appear in more than one threat category when differing threat statuses have been attached to different subspecies, varieties or pathotypes. Legon and Henrici (2005) list 248 species of rust fungi from mainland Britain. Since this list was published five more rust species have been added to the British list giving a total of 253 species. Records of 199 rust species within 225 separate taxa have been traced from Wales (79% of the British total) occurring in 1036 rust/host taxa interactions. This compares with the 68% of the British mainland lichen mycota reported from Wales (Woods 2010) and almost 75% of the British bryophyte flora (Bosanquet & Dines 2011). Yet Wales makes up only 11% of the total British mainland surface area and this further emphasises the exceptional biodiversity of the principality.

Of the 199 rust species recorded from Wales 13 only occur on neophyte vascular plants and the conservation statuses of these species have not been evaluated. Of the 186 rust species recorded on native or archaeophyte hosts, six species (3%) and seven taxa are considered to be extinct in Wales. Of the 180 remaining rust species on native hosts, the continued existence of 50 species (28%) is considered to be threatened or likely to become threatened in some way, with a further 15 taxa (8%) lacking in sufficient data to currently ascribe them to a threat category. This compares with 26% of lichens and 19% of bryophytes and vascular plants considered under threat and 2% of lichens and 3% of bryophytes and vascular plants considered extinct in Wales (Woods (2010), Bosanquet and Dines (2011) and Dines (2008) respectively).

The numbers in each category are recorded in the table below.

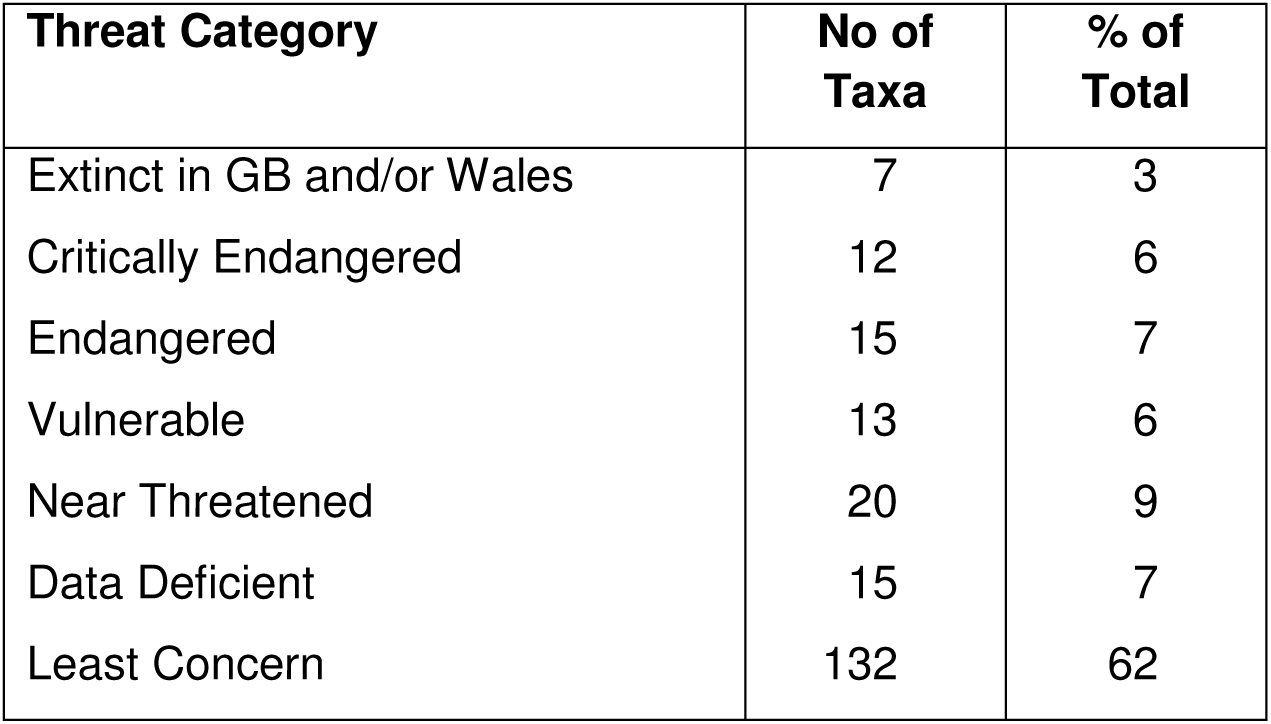

| **Threat Category** | **No of Taxa** | **% of Total** |
| --- | --- | --- |
| Extinct in GB and/or Wales | 7 | 3 |
| Critically Endangered | 12 | 6 |
| Endangered | 15 | 7 |
| Vulnerable | 13 | 6 |
| Near Threatened | 20 | 9 |
| Data Deficient | 15 | 7 |
| Least Concern | 132 | 62 |

Discounting the 3% that are probably extinct and those in the near threatened category, 19% of the extant Welsh rust taxa are considered to be under immediate threat. This compares with 16% of the lichens (Woods 2010), 18% of the bryophytes (Bosanquet 2011) and 17% of the vascular plants (Dines 2008). Given the dependence of rust fungi on vascular plants, a figure somewhat higher than the 17% threat percentage of the vascular plants might have been expected.

An examination of the habitats occupied by threatened rust taxa shows the largest number to occur in fens and marshy grasslands, closely followed by those of woodlands and woodland edges. Heathlands and sand dune systems support the only other significant number of taxa.

#### 8.1 a Extinct Taxa

Seven taxa are considered to have become regionally extinct in Wales including one rust of uncertain taxonomic status that may have become extinct in Britain and two species that may have also become extinct in Britain. Most have not been seen for over 60 years in Wales and their loss may pre-date the great agricultural improvements of the second half of the 20^th^ century. All losses have occurred in the northern half of Wales and involve both common and rare hosts and a range of habitats. For more details on individual species see the notes section below.

| Rust Taxon | Host | Wales Red List | Date Last Seen | VC Last Seen | Habitat |
| --- | --- | --- | --- | --- | --- |
| *Melampsora lini*<br>var. *radiolae* | *Radiola linoides* | EX | 1936 | 52 | Damp bare ground near coast |
| *Puccinia apii* | *Apium graveolens* | RE | 1913 | 51 | Damp open habitats near coast |
| *Puccinia heraclei* | *Heracleum sphondylium* | RE | 1909 | 48 | Wayside? |
| *Puccinia major* | *Crepis paludosa* | RE | 1887 | 48 | Wet woodland |
| *Puccinia uliginosa* | *Carex nigra* | RE | 1948 | 52 | Fen? |
| *Puccinia veronicae-longifoliae* | *Veronica spicata* | RE or EX? | 1950 | 50 | Basic dry grassland |
| *Uromyces inaequalis* | *Silene nutans* | RE or EX? | 1921 | 51 | Calcareous grassland |

#### 8.1 b Critically Endangered Taxa

Twelve rust taxa are considered to be Critically Endangered in Wales. Only one rust species occurs on a host to which Dines (2008) ascribes a threat category (*Scorzonera humilis* VU). Three taxa occur on hosts with very patchy British distributions and the Welsh populations are distant from those in England. Seven taxa are inexplicably rare given the general distribution and abundance of their hosts. Only one or possibly two species of Critically Endangered rusts may regularly alternate between two vascular plant hosts.

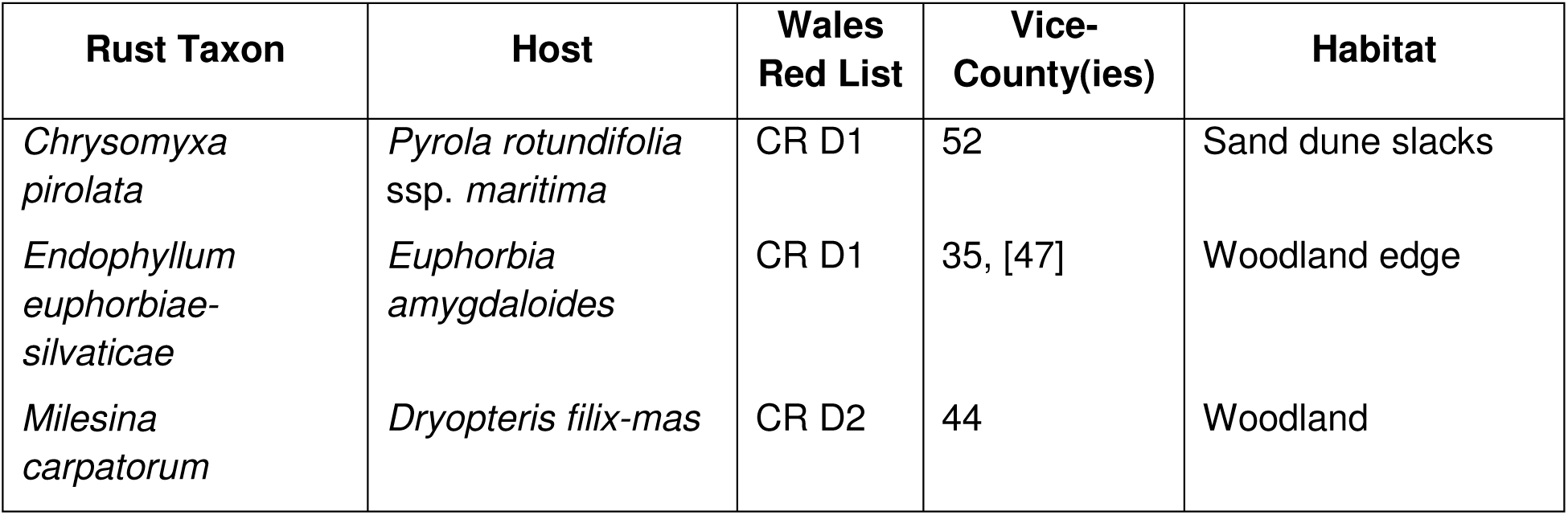
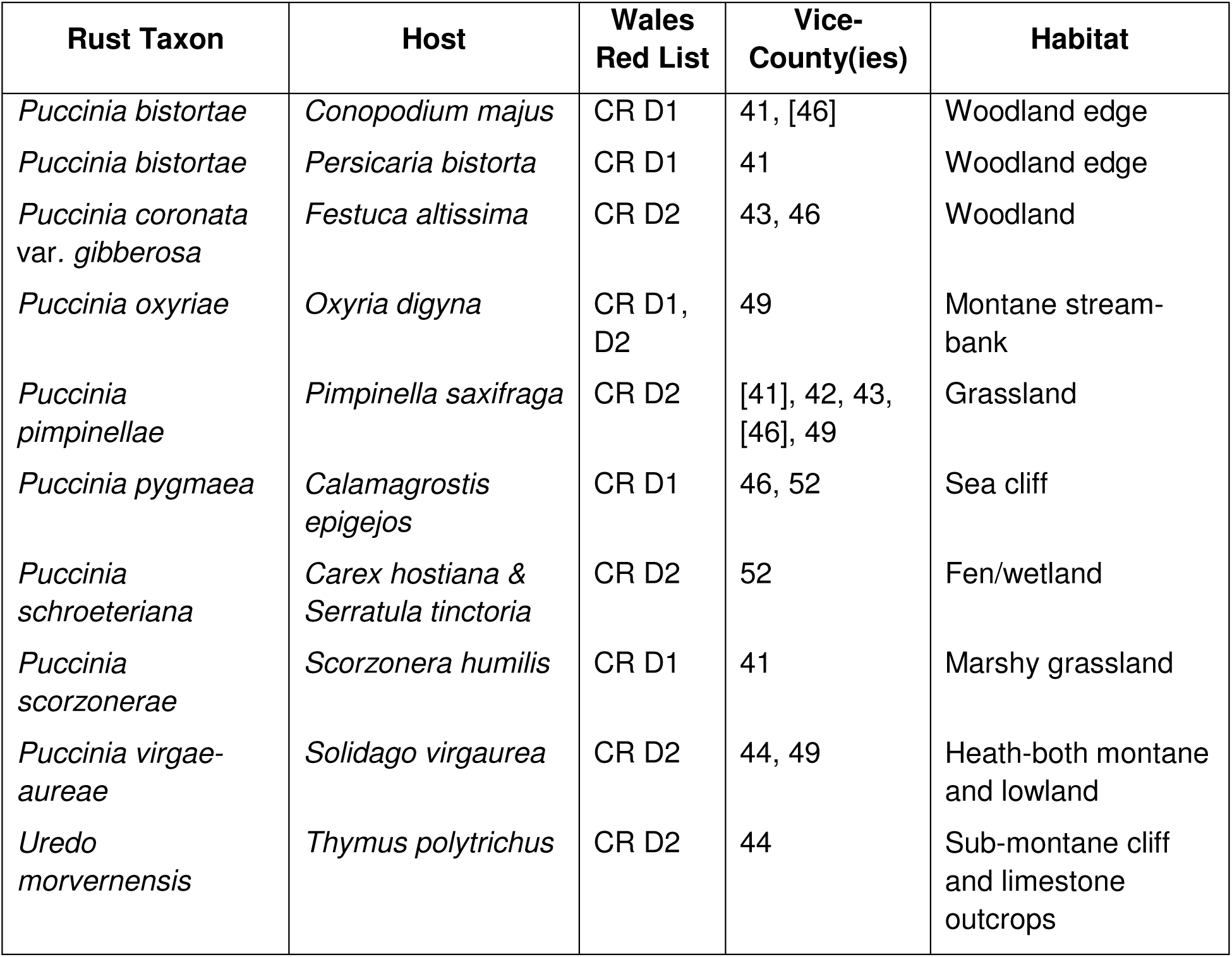

| Rust Taxon | Host | Wales Red List | Vice-County(ies) | Habitat |
| --- | --- | --- | --- | --- |
| *Chrysomyxa pirolata* | *Pyrola rotundifolia*<br>ssp. *maritima* | CR D1 | 52 | Sand dune slacks |
| *Endophyllum euphorbiae-silvaticae* | *Euphorbia amygdaloides* | CR D1 | 35, [47] | Woodland edge |
| *Milesina carpatorum* | *Dryopteris filix-mas* | CR D2 | 44 | Woodland |
| *Puccinia bistortae* | *Conopodium majus* | CR D1 | 41, [46] | Woodland edge |
| *Puccinia bistortae* | *Persicaria bistorta* | CR D1 | 41 | Woodland edge |
| *Puccinia coronata*<br>var. *gibberosa* | *Festuca altissima* | CR D2 | 43, 46 | Woodland |
| *Puccinia oxyriae* | *Oxyria digyna* | CR D1, D2 | 49 | Montane stream-bank |
| *Puccinia pimpinellae* | *Pimpinella saxifraga* | CR D2 | [41], 42, 43, [46], 49 | Grassland |
| *Puccinia pygmaea* | *Calamagrostis epigejos* | CR D1 | 46, 52 | Sea cliff |
| *Puccinia schroeteriana* | *Carex hostiana* & *Serratula tinctoria* | CR D2 | 52 | Fen/wetland |
| *Puccinia scorzonerae* | *Scorzonera humilis* | CR D1 | 41 | Marshy grassland |
| *Puccinia virgae-aureae* | *Solidago virgaurea* | CR D2 | 44, 49 | Heath-both montane and lowland |
| *Uredo morvernensis* | *Thymus polytrichus* | CR D2 | 44 | Sub-montane cliff and limestone outcrops |

#### 8.1 c Endangered Taxa

Fifteen Endangered rust taxa have been identified. Only one, *Melampsora arctica* occurs on a host given a threat status in Wales by Dines (2008), whilst four species are confined to hosts which have a rather limited range or disjunct distribution in Wales. Ten occur on relatively widespread and even common hosts. No single host habitat predominates, though four of the rust taxa occur in fens.

| Rust Taxon | Host | Wales Red List | Vice-County(ies) | Habitat |
| --- | --- | --- | --- | --- |
| *Melampsora arctica* | *Salix herbacea* | EN D | 49 | Montane heath |
| *Melampsora amygdalinae* | *Salix triandra* | EN B2 | 44, 46 | Fen |
| *Puccinia angelicae* | *Angelica sylvestris* | EN D | [35], 41, 44, 46, [49], 52 | Fen and coastal ditchside |
| *Puccinia cladii* | *Cladium mariscus* | EN D | 41 | Fen |
| *Puccinia dioicae* | *Carex dioica* | EN D | 42, 49 | Wetland |
| *Puccinia dioicae* | *Cirsium palustre* | EN D | 48, 49 | Wetland |
| *Puccinia eriophorae* | *Solidago virgaurea* & *Trichophorum cespitosum* agg. | EN D | 49 | Montane & sub-heath |
| *Puccinia fergussonii* | *Viola palustris* | EN D | 41, 42, 44, 46, 48, 49, 52 | Wetland |
| *Puccinia galii-cruciatae* | *Cruciata laevipes* | EN B2, D | 41, 42, 44 | Neutral Grassland |
| *Puccinia hydrocotyles* | *Hydrocotyle vulgaris* | EN D | 44, 46, 52 | Poor fen and dune slack |
| *Puccinia maculosa* | *Mycelis muralis* | EN D | 42-44, [48], [49], 51 | Rock outcrops/walls |
| *Puccinia saxifragae* | *Saxifraga granulata* | EN D | 43 | Woodland |
| *Puccinia saxifragae* | *Saxifraga stellaris* | EN D | [46], 48, 49 | Sub-montane flush |
| *Puccinia thymi* | *Thymus polytrichus* | EN D | 42, 45 | Calcareous grassland |
| *Puccinia urticata* var. *biporula* | *Carex pallescens* | EN D | 41, 42, 46 | Garden/wet pasture |
| *Uromyces scrophulariae* | *Scrophularia auriculata* | EN D | 44, [47] | Wayside |
| *Uromyces sommerfeltii* | *Solidago virgaurea* | EN D | 46, 49 | Montane heath |

#### 8.1 d Vulnerable Taxa

Thirteen rust taxa have been placed in the category of Vulnerable. None of their hosts are considered to be under threat. Four rust taxa probably alternate between two or more host taxa regularly in Wales and six occur associated with woodlands or woodland edges. Seven species occur on hosts associated with open ground such as arable fields, waysides and sand dunes.

| Rust Taxon | Host | Wales Red List | Vice-County(ies) | Habitat |
| --- | --- | --- | --- | --- |
| *Gymnosporangium clavariiforme* | *Juniperus communis* & *Crataegus monogyna* | VU D | 49 | Sub-montane heath |
| *Melampsora allii-fragilis* | *Allium ursinum* | VU D2 | 42, 46, 49, 52 | Wet woodland |
| *Melampsora allii-fragilis* | *Salix × fragilis* | VU D2 | 43, 44 | River bank |
| *Melampsora allii-fragilis* | *Salix pentandra* | VU D2 | 44 | Hedges and stream banks |
| *Milesina vogesiaca* | *Polystichum setiferum* | VU D1, D2 | 44 | Calcareous woodland |
| *Puccinia dioicae* var. *schoeleriana* | *Carex arenaria* | VU D | 44, 52 | Sand dunes |
| *Puccinia dioicae* var. *schoeleriana* | *Senecio jacobaea* | VU D | 44, 48, 52 | Waste ground |
| *Puccinia dioicae* var. *schoeleriana* | *Senecio* sp. | VU D | 35 | Sand dunes |
| *Puccinia elymi* | *Ammophila arenaria* | VU D | 41, 45, 46, 49 | Sand dunes |
| *Puccinia elymi* | *Leymus arenarius* | VU D | 44, 46, 49, 51, 52 | Sand dunes |
| *Puccinia hysterium* | *Tragopogon pratensis* | VU D1 | 41, 45 | Wayside |
| *Puccinia luzulae* | *Luzula* sp. | VU D | 47 | ? |
| *Puccinia luzulae* | *Luzula pilosa* | VU D | 44, [46], 48, [49] | Woodland |
| *Puccinia nemoralis* | *Melampyrum pratense* & *Molinia caerulea* | VU D2 | 48, 49 | Sessile oak woodland |
| *Puccinia nitida* | *Aethusa cynapium* ssp. *agrestis* | VU D2 | 35, 46, [49] | Arable |
| *Puccinia nitida* | *Aethusa cynapium* ssp. *cynapium* | VU D2 | 43, 47 | Arable |
| *Puccinia polygoni-amphibii* var. *convolvuli* | *Fallopia convolvulus* | VU D | [41], 42, 44, 49 | Arable |
| *Puccinia urticata* var. *urticae-ripariae* | *Carex riparia* | VU D1, D2 | 44, 46 | Fen |
| *Triphragmium filipendulae* | *Filipendula vulgaris* | VU B, D1 | 49 | Calcareous grassland |
| *Uromyces viciae-fabae* var. *orobi* | *Lathyrus linifolius* var. *montanus* | VU D | 43, 47, 48 | Woodland edge |

#### 8.1 e Near Threatened Taxa

None of the 20 Near Threatened rust taxa occur on hosts that are considered to be threatened in Wales. The largest proportion (8 taxa) occurs in grasslands and on waysides, followed by those occurring in lakes, by rivers and in wetlands (5 taxa) and then woodland (4 taxa). Three other taxa occur on moorland and/or cliffs.

*Chrysomyxa empetri* on *Empetrum nigrum*.

*Cronartium quercuum* on *Quercus cerris, Q. ilex, Q. petraea* and *Q. robur. Hyalopsora polypodii* on *Cystopteris fragilis*.

*Melampsora salicis-albae* on *Salix alba*.

*Melampsorella caryophyllacearum* on *Cerastium fontanum* sspp. *holosteoides* and *vulgare,*

*C. glomeratum, Stellaria alsine, S. graminea, S. holostea* and *S. media. Ochropsora ariae* on *Anemone nemorosa*.

*Puccinia calthae* on *Caltha palustris*.

*Puccinia caricina* var*. magnusii* on *Carex riparia*.

*Puccinia caricina* var*. ribis-nigri-paniculatae* on *Carex paniculata*.

*Puccinia conii* on *Conium maculatum*.

*Puccinia difformis* on *Galium aparine*.

*Puccinia crepidicola* on *Crepis biennis, C. capillaris* and *C. vesicaria*.

*Puccinia glomerata* on *Senecio aquaticus* and *S. jacobaea*.

*Puccinia opizii* on *Carex muricata* ssp*. pairae, C. paniculata* and *C. spicata*.

*Puccinia primulae* on *P. veris* and *P. vulgaris*.

*Puccinia urticata* var*. urticae-inflatae* on *Carex rostrata*.

*Pucciniastrum agrimoniae* on *Agrimonia* sp. and *A. eupatoria*.

*Trachyspora intrusa* on *Alchemilla* spp*., A. glabra, A. vulgaris* and *A. xanthochlora*.

*Uromyces ervi* on *Vicia hirsuta* and *V. tetrasperma*.

*Uromyces sparsus* on *Spergularia marina* and *S. media*.

#### 8.1 f Data Deficient Taxa

Fifteen taxa lack sufficient information to be assigned to a threat category. Work on these species is a high priority since many of them may be critically endangered.

*Coleosporium tussilaginis* on *Euphrasia* spp*., Parentucellia viscosa* and *Senecio sylvaticus*.

*Gymnosporangium confusum* on *Crataegus monogyna* and *Mespilus germanica*.

*Gymnosporangium cornutum* on *Sorbus aucuparia* and *Juniperus communis*.

*Melampsora allii-populina* on *Populus nigra, P. trichocarpa* and *Arum maculatum*.

*Melampsora epitea* on *Dactylorhiza* spp*., Epipactis* spp*., Platanthera chlorantha* and *Salix repens*.

*Melampsora laricis-pentandrae* on *Salix pentandra. Melampsora ribesii-viminalis* on *Salix viminalis*.

*Phragmidium potentillae* on *Potentilla anglica* and *P.* × *suberecta*.

*Puccinia campanulae* on *Campanula rotundifolia* and *Jasione montana*.

*Puccinia cancellata* on *Juncus acutus* and *J. maritimus*.

*Puccinia caricina* var*. pringsheimiana* on *Ribes uva-crispa* and *Carex acuta*.

*Puccinia chaerophylli* on *Anthriscus sylvestris* and *Chaerophyllum temulentum*.

*Puccinia graminis* on a wide range of grasses.

*Puccinia scirpi* on either *Nymphoides peltata* or *Schoenoplectus lacustris* or both.

*Puccinia urticata* var*. urticae-acutae* on *Carex nigra*.

### 8.2 Census Catalogue

Recording effort has varied greatly between the vice-counties with Monmouthshire, Pembrokeshire, Montgomeryshire, Merionethshire, Denbighshire and Flintshire receiving the least attention and Carmarthenshire, Cardiganshire, Caernarvonshire and Anglesey the most. Despite considerable effort within Radnorshire, its small size and lack of coast or significant areas of calcareous rock have produced only a rather limited range of both rust fungi and vascular plants. The table below displays the total number of rust species and the total number of rust taxa (including subspecies, varieties etc.) recorded for each Welsh vice-county and compares this number with the total number of vascular plants (Ellis 1983).

These numbers can be compared with other vice-counties outside Wales. Preece (1995) notes 104 rust species (127 taxa) from Shropshire. This accords with the rather similar numbers recorded from the Welsh vice-counties along the middle March. Clark (1980) reports 94 rust species from Warwickshire. Chris Yeates maintains a database of rust records from the five vice-counties that make up the old county of Yorkshire. He has records (pers. comm.) of 183 rust taxa (166 species) from this area. Given that Yorkshire at 6081 square miles is 80% the size of Wales, the two areas have very similar ratios of species to area, with Yorkshire having one rust species to 37 square miles and Wales one species to 38 square miles. Dorset has records of 143 species (Bryan Edwards pers. comm.) whilst the FRDBI indicates that three vice-counties have more than 150 rust taxa i.e. Surrey with 166, East Norfolk with 161 and North East Yorkshire with 157.

The high totals of 164 rust species (186 taxa) from Carmarthenshire and 161 species (178 taxa) from Caernarvonshire reflect both the intensive recording effort but also vice-counties rich in habitats, including sub-montane cliffs, limestone grassland, extensive areas of wetland and a diverse coastline supporting many potential hosts.

The census catalogue records demonstrate the wide distribution of mycological highlights in Wales. The Isle of Anglesey and its sand dune systems support, or have in the past supported, *Chrysomyxa pirolata* on round-leaved wintergreen, *Puccinia hydrocotyles* on marsh pennywort and *Melampsora lini* var. *radiolae* on allseed, the only ever British record on this host. Its fens support nationally important populations of *Puccinia schroeteriana* on tawny sedge and saw-wort and of *P. commutata* on common valerian. The mountains of Snowdonia within Caernarvonshire have records of montane species at the southern edge of their British range in *Melampsora arctica* on dwarf willow, *Puccinia oxyriae* on mountain sorrel and the only British record of the rust *Pucciniastrum epilobii* on chickweed willowherb *Epilobium alsinifolium*. Other notable taxa include *Uromyces sommerfeltii, Puccinia virgae-aureae* and *P. eriophorae*, all on golden rod, with the latter species also occurring on deergrass. Merionethshire supported the last known population in Wales of *P. major* on marsh hawk’s-beard.

Cardiganshire has notable populations in its wetlands of *Puccinia hydrocotyles* on marsh pennywort, *P. commutata* on common valerian and *P. fergussonii* on marsh violet, and a very disjunct locality on the coast for *Puccinia pygmaea* on wood small-reed. Radnorshire supports the only known population of *P. saxifragae* on meadow saxifrage in Wales and Breconshire has *P. hieracii* on a number of its endemic hawkweeds.

Of the South Wales vice-counties Carmarthenshire stands out for the diversity of rust taxa, particularly those on rush, poplar, willow and sedge species as well as rusts of salt marsh plants. Historically, south-east Carmarthenshire was heavily industrialised but over the last two decades a decline in heavy industry has led initially to large areas of brownfield sites and reclamation plantings. These plantings included many varieties of poplar and willow including hybrids which are now infected with rusts. Reclaimed opencast sites predominantly planted up with *Alnus* are notable for large populations of the rust *Melampsoridium hiratsukanum -* a recent arrival to the UK from south-east Asia.

Glamorgan has one of only two known extant populations in Britain of *Puccinia cladii* on saw sedge.

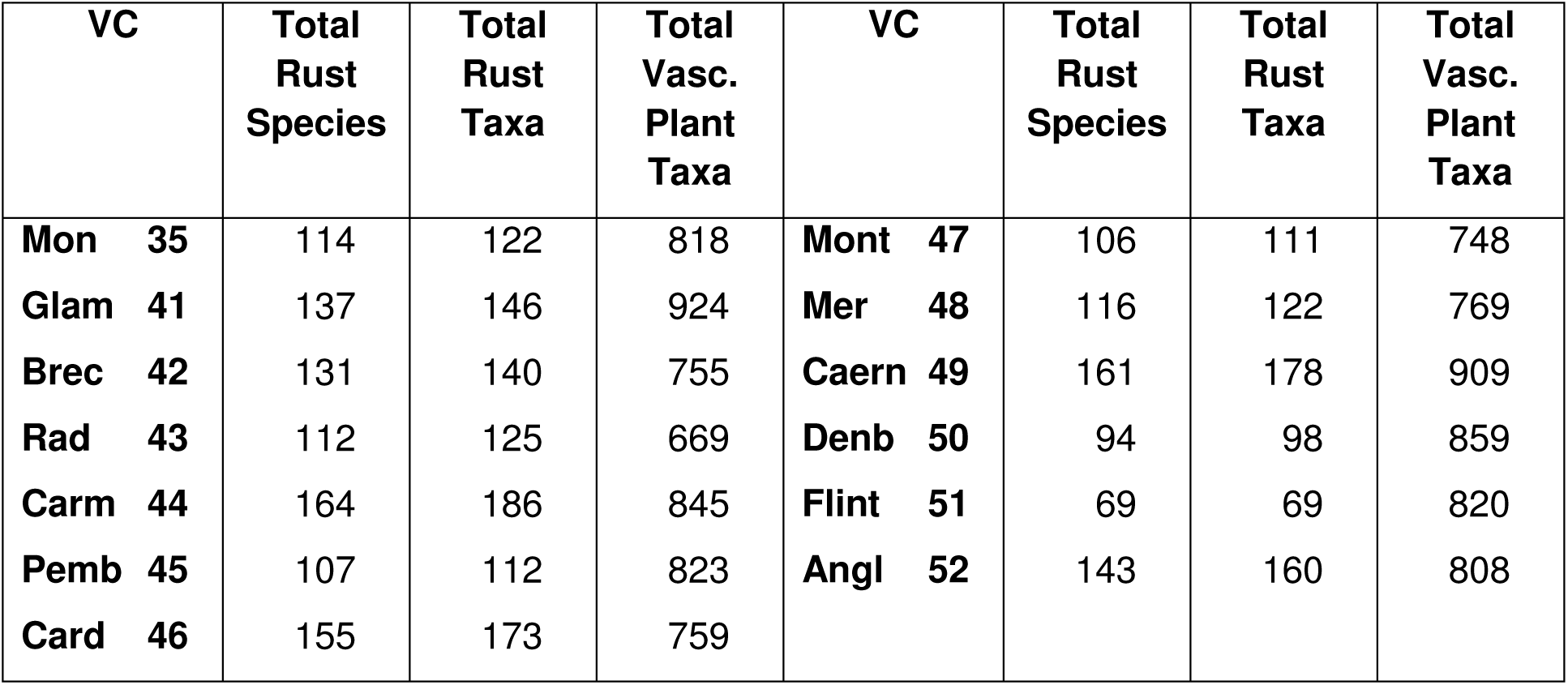

## 9 Notes on Individual Taxa

Notes are provided on all species with a conservation status to explain the reasoning behind such a decision. A few additional taxa have been provided with notes where, for example, records on the FRDBI and CATE2 might indicate a threat status to be warranted but recent recording in Wales indicates otherwise. Notes are also provided where problems of identification or uncertain taxonomy preclude a decision, or a rust is confined to cultivated hosts in Wales.

### Chrysomyxa empetri (Pers.) J. Schröt

This rust is of scattered occurrence in Wales on crowberry *Empetrum nigrum*, a widespread dwarf shrub. The orange uredinia occur on the upper leaf surface, though since the leaves are rolled into a tube most uredinia occur on what appears to be the lower surface. It can nevertheless be quite conspicuous, but the possibility exists that it may have been overlooked. Twenty populations have been recently located in Wales. The greatest concentration occurs in the mountains of Snowdonia with 12 monad records from Caernarvonshire, where it is found at up to 1000m, often on exposed ridges, and from five sites in Cardiganshire. There are single recent records from Monmouthshire, Glamorganshire, Breconshire and Denbighshire and an old record from Merionethshire (near Bala 1865). Whilst all records from Mid Wales are likely to be from *E. nigrum* ssp. *nigrum*, further work is required to establish the proportions of records on this ssp. as opposed to ssp. *hermaphroditum* from North Wales since Paul Reade reports this rust on at least one population of ssp. *hermaphroditum* on the summit of Tryfan at 915m. There are nearly 40 records from Scotland on crowberry in the last 50 years (FRDBI and Paul Smith pers. comm.), mostly from the islands off the north and west coasts. There are no English or Irish records. Evans *et al*. (2006) have placed this rust in the **Near Threatened** category in the Provisional British RDL of Fungi, a category it is proposed to adopt in Wales.

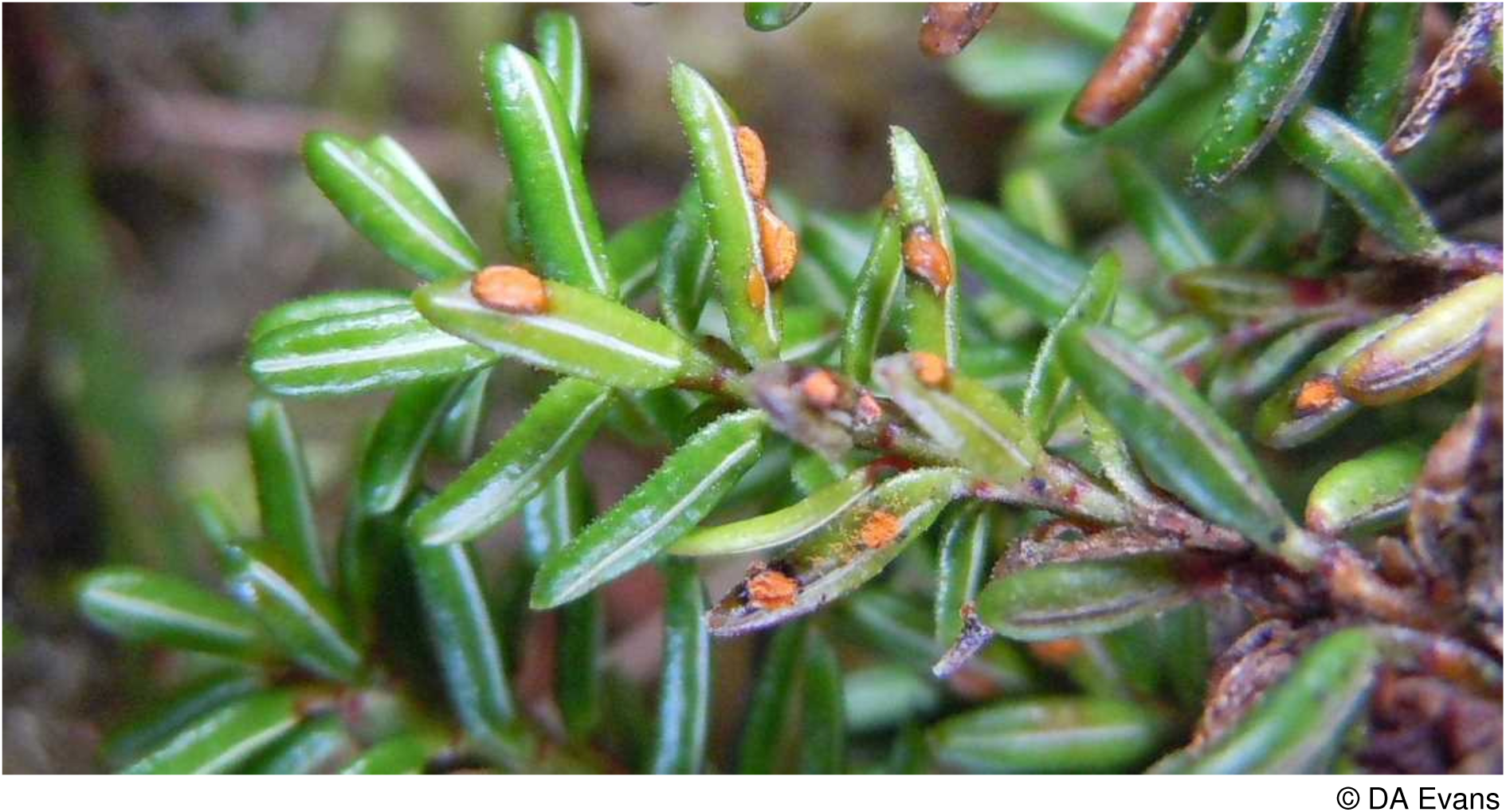

### Chrysomyxa pirolata (Schwein.) G. Winter

Known in Wales only from round-leaved wintergreen *Pyrola rotundifolia* var. *maritima* on sand dunes at Newborough Warren, Anglesey, the size of the population is unknown and recent attempts to re-find it have failed. Elsewhere it occurs on dunes near Southport, South Lancashire and in Westmorland. There is also a recent record from an unspecified *Pyrola* sp. from S. Wilts on the FRDBI and old records from *P. rotundifolia* from Shropshire and Gloucestershire. All Scottish records are from the 1800’s. It does not appear to alternate with *Picea* spp. in Britain as it does elsewhere.

In view of it being known from a single locality in Wales it is accorded a **Critically Endangered D1** conservation evaluation. Evans *et al*. (2006) have placed this rust in the same category in the Provisional British RDL of Fungi. It has a British Biodiversity Action Plan and is listed on Section 42 of the Natural Environment and Rural Communities Act (2006) as being a species of principal importance in Wales.

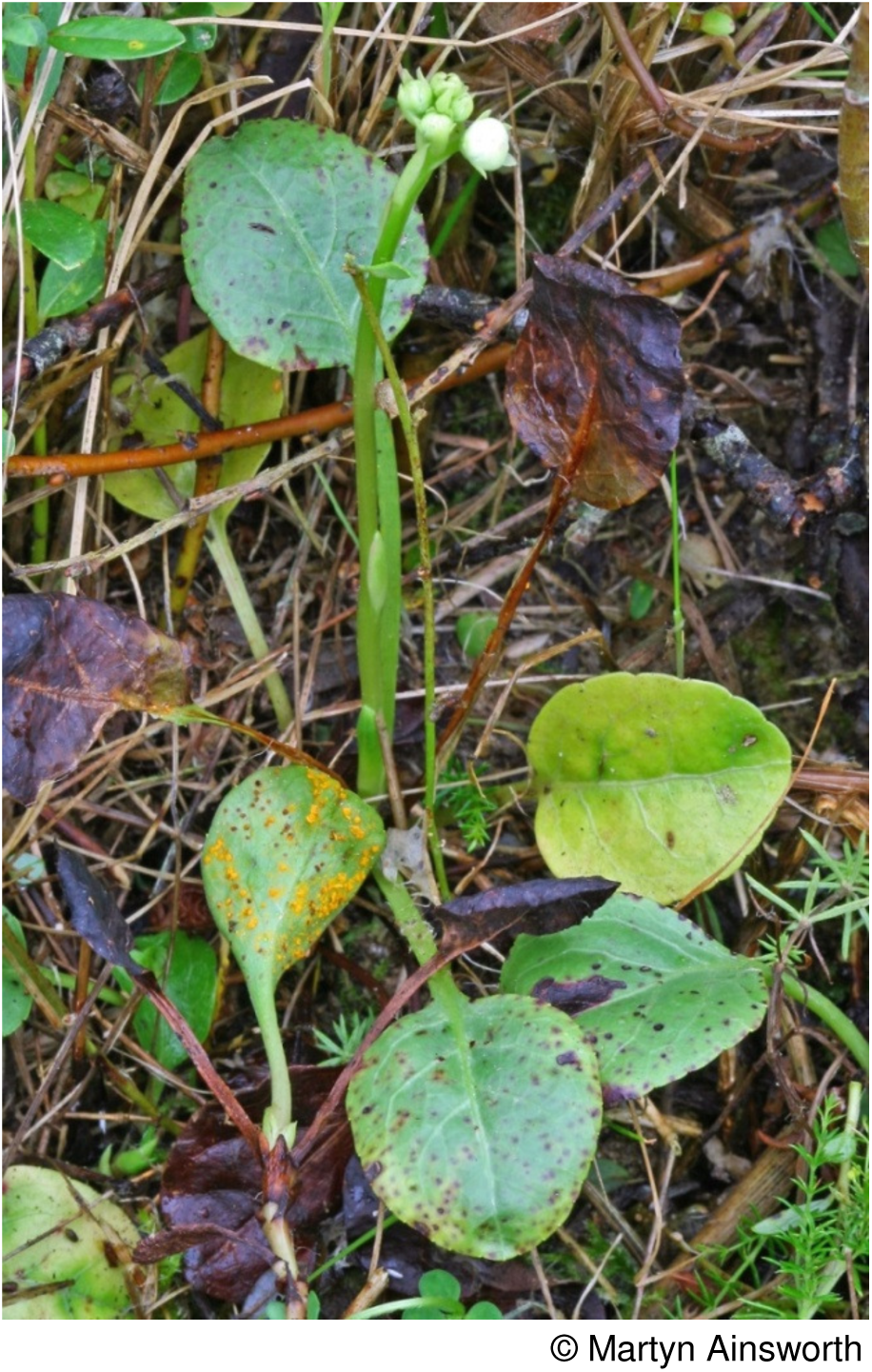

### Coleosporium tussilaginis (Pers.) Lév

This rust develops spermogonia and aecia on *Pinus sylvestris* and *P. nigra* neither of which are considered to have native populations in Wales. There are few records but searches have been few. Uredinia and telia develop on a wide range of hosts including harebell *Campanula rotundifolia* and garden *Campanula* spp., eyebright *Euphrasia officinalis* agg., common cow-wheat *Melampyrum pratense,* red bartsia *Odontites vernus,* yellow bartsia *Parentucellia viscosa,* butterbur *Petasites hybridus,* yellow-rattle *Rhinanthus minor,* ragwort and groundsel species *Senecio* spp., sow-thistle species *Sonchus* spp. and colt’s-foot *Tussilago farfara*. It is frequent in Wales on some of these hosts but rarer on others. Once considered a multiplicity of species based on the host plant, it is clear that such distinctions could not be maintained as cross-infection from one host to another is possible in at least some cases. Further work is required to establish the existence of distinct races worthy of conservation status. For the time being every effort should be made to conserve diverse populations of eyebright species, particularly the rarer ones, bearing this rust and any infected populations of *yellow bartsia*. Records on wood groundsel *Senecio sylvaticus* are sparse and notable populations should be protected. Braithwaite *et al*. (2006) note declines in abundance of harebell, hay-rattle and cow-wheat.

A conservation evaluation of **Data Deficient** is proposed for populations on *Euphrasia arctica* ssp. *arctica* and ssp. *borealis, E. confusa, E. nemorosa, E. officinalis* ssp. *pratensis* (= *E. rostkoviana*) and ssp. *anglica* (= *E. anglica*), *E. tetraquetra, Parentucellia viscosa* and *Senecio sylvaticus*.

### Cronartium quercuum (Berk.) Miyabe ex Shirai

The golden uredinia on acorn cups and the underside of oak leaves are quite distinctive, though at times hard to locate despite the frequency of the host. In Wales there are recent records on sessile oak *Quercus petraea* from two sites in each of Breconshire, Pembrokeshire and Cardiganshire, a single site in Merionethshire and 15 sites in Caernarvonshire. It has been found on pedunculate oak *Q. robur* in one site in Carmarthenshire and on the non-natives, evergreen oak *Q. ilex* in Glamorgan and Turkey oak *Q. cerris* in two sites in Bangor, Caernarvonshire. Records are more frequent on young trees and those trimmed in hedgerows. Very rarely is a whole tree infected and some trees infected in one year seem uninfected in subsequent years. There are no recent records on native oak *Quercus* species from mainland England or Scotland on the FRDBI. In view of the small and scattered population (even including the populations on non-native oak species) occupying 10 hectads in Wales it is considered to be **Near Threatened**.

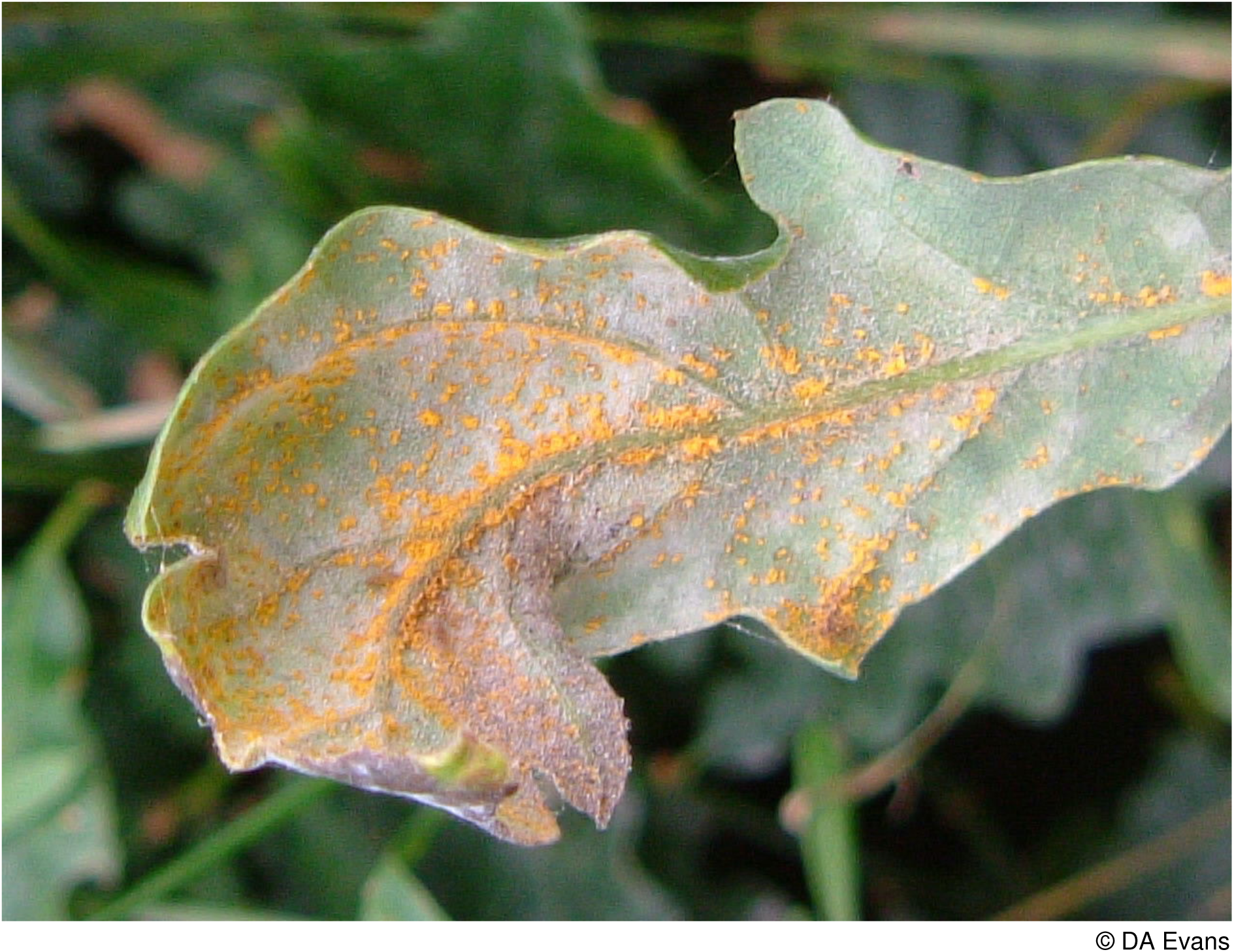

### Endophyllum euphorbiae-silvaticae (DC.) G. Winter

The distinctive yellow-etiolated orange cup-covered infected shoots of wood spurge *Euphorbia amygdaloides* cannot easily be missed. The Welsh populations of wood spurge are very much on the western edge of its British range so it is perhaps not surprising that it is a rare rust in Wales. Vize reported it from near Welshpool and Forden in Montgomeryshire in the 1800’s. Whilst its host persists in a number of mostly estate woodlands in the area the rust has not been reported again. The sole recent Welsh record is from Monmouthshire, the size of which population is not known. In England with the exception of a single Derbyshire record, all recent records are from the SW of England with records from Herefordshire, Gloucestershire, Dorset and Devon (FRDBI). The sole Welsh record, taken with a history of decline, warrants a critically endangered conservation evaluation but since the inoculum may come from the adjacent English populations IUCN guidance recommends downgrading by a category unless the adjacent population is showing a decline. From the limited information available on the FRDBI and on CATE2 of the ABFG there appear to be no recent records from Kent, Oxfordshire or the North of England and in consequence a decline is indicated.

Therefore a **Critically Endangered D1** evaluation in Wales is warranted though other criteria may apply once the population size can be determined.

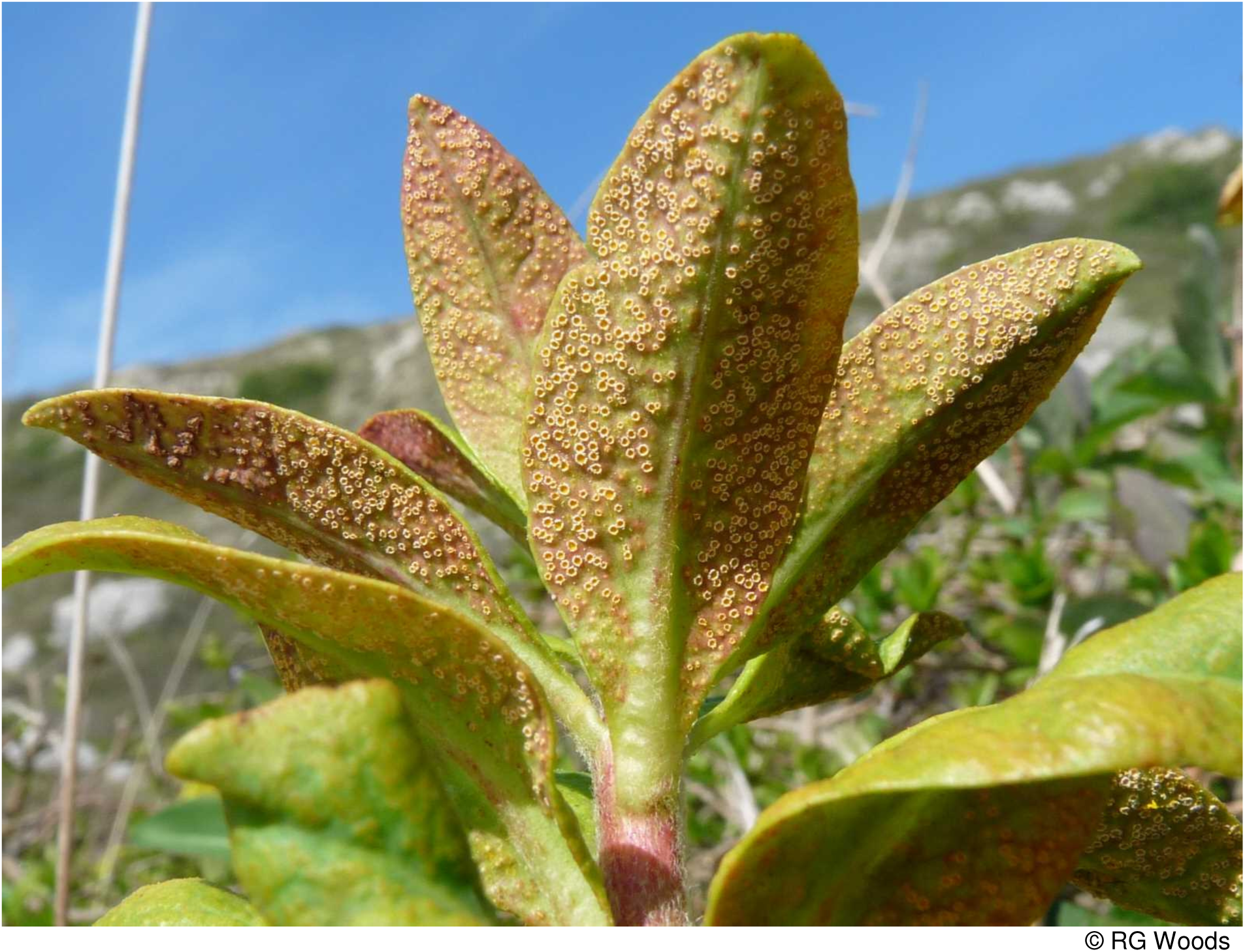

### Gymnosporangium species

There are four species of *Gymnosporangium* recorded from Britain. All alternate between juniper *Juniperus* species and woody members of the Rosaceae. Three species have been recorded recently from Wales but only *G. clavariiforme* has been found in a natural or non-man-made habitat.

### Gymnosporangium clavariiforme (Wulfen) DC

Alternating between common juniper *Juniperus communis* and hawthorn *Crataegus monogyna*, this rust is of widespread occurrence in Scotland and England. In Caernarvonshire of note is its recent discovery on native juniper (as var. *nana*) in Cwm

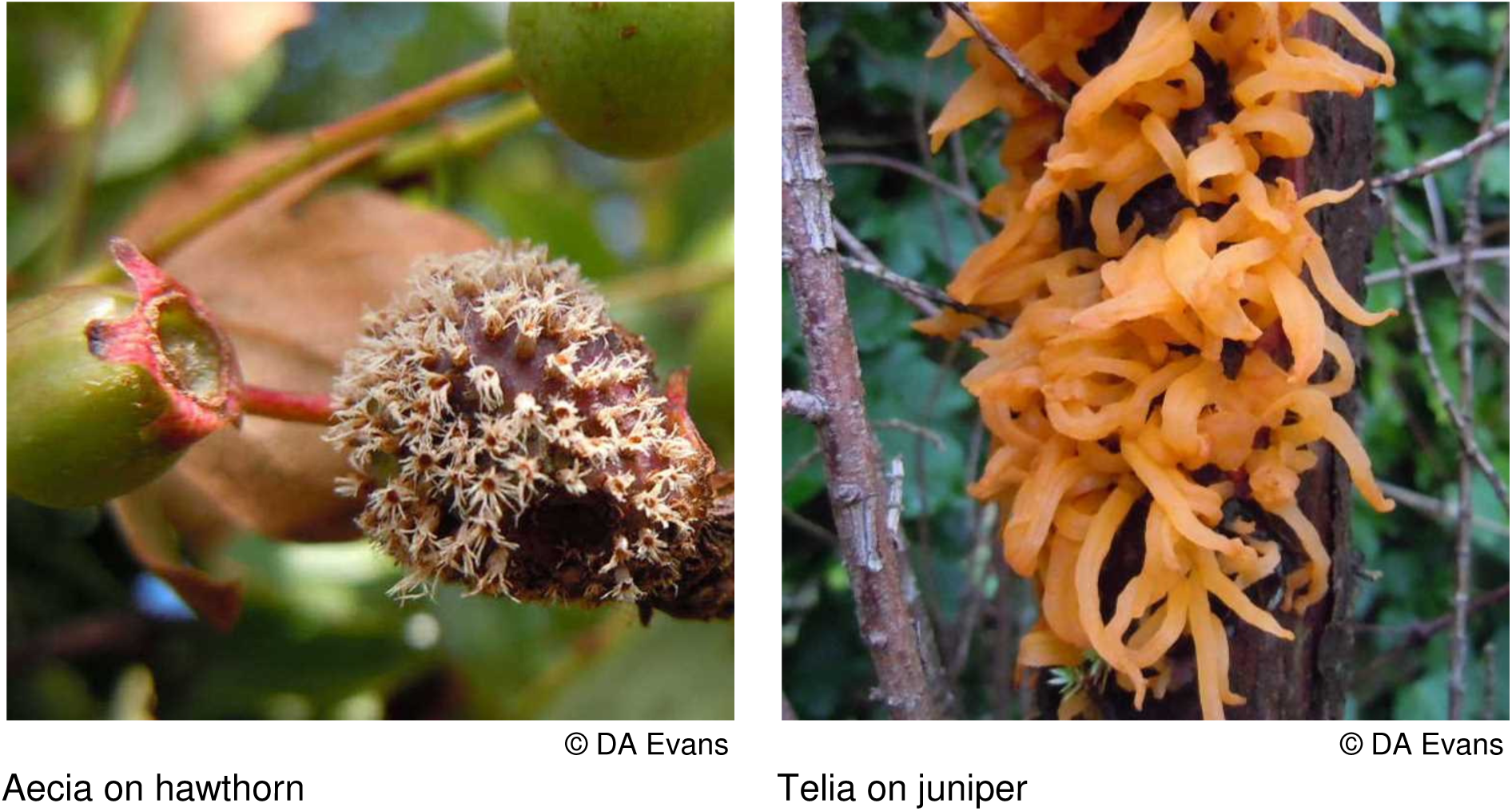

Carregog at over 600m below Snowdon by Iwan Edgar (confirmed Debbie Evans). It also occurs on both hawthorn and native juniper on a limestone escarpment at Nant y Gamar and on hawthorn at Bryn Pydew just east of the previous site, whilst Aron (2005) notes records on juniper (species not given) in Merion Lane, Bangor (presumably a garden) and on hawthorn in Nantporth and there are additional more recent records from a scatter of hawthorn bushes across the vice-county. There is one record on hawthorn and juniper in a Cardiganshire garden, whilst at Menai Bridge in Anglesey a common juniper in a garden is infected together with several hawthorns in the general area. There are in addition, on hawthorn, two records from Monmouthshire and two from Radnorshire, the latter two in trackside hedges. In at least one of the Radnorshire sites common juniper is present nearby in a garden, though no infections were detected on it. There are also two records on juniper from Montgomeryshire but since common juniper is not known in the wild from Montgomeryshire (Trueman *et al*. 1995) these records must be of garden origin. Evaluating its status on the basis of its occurrences on native species remote from gardens i.e. on juniper and hawthorn in Snowdonia it is considered to be **Vulnerable D** since it probably occurs on fewer than a 1000 host bushes.

### Gymnosporangium confusum *Plowr*

Alternating between the cultivated juniper *Juniperus sabina* and hawthorn *Crataegus* spp. and medlar (*Mespilus germanica*), there are numerous records of this rust from hawthorn and less frequently *J. sabina* in central England. In Wales there are three records from hawthorn in gardens and hedges in and around Bangor, Caernarvonshire and five records from medlar in gardens in this VC. There are single records from hawthorn in Monmouthshire and Radnorshire and one record from medlar in Anglesey.

It is doubtful whether this species would survive in the wild without garden populations of *J. sabina* and it may well be an introduced species. Pending resolution of this issue and until more gardens can be surveyed it is considered to be **Data Deficient**.

### Gymnosporangium cornutum Arthur ex F. Kern

The FRDBI notes a record in the literature of this rust (originally incorrectly considered to be *G. juniperi*) from Swallow Falls, Caernarvonshire in 1924. No host is given and the source of the record is said to be from the JNCC literature records database. Alternating between juniper *Juniperus communis* and rowan *Sorbus aucuparia,* both hosts are present in the area, though the nearest recent records of this mostly northern species on the FRDBI to Wales are North Yorkshire and Buckinghamshire and on CATE2 on an unspecified host from Cheshire. In 1988 this rust was collected from North Wales on rowan but unfortunately no further locality details were given. The conservation status of this fungus is considered to be **Data Deficient**.

### Gymnosporangium sabinae (Dicks.) G. Winter

In Wales known only from a wide range of pear *Pyrus communis* cultivars in both Caernarvonshire and Anglesey. If searched for it will probably be found elsewhere. Since it has not been found in the wild its conservation status has not been evaluated.

### Hyalopsora polypodii (Pers.) Magnus

The orange uredinia are a distinctive feature of this rust on the delicate fronds of brittle bladder-fern *Cystopteris fragilis*. Noted recently from seven vice-counties in Wales, only in Breconshire and Carmarthenshire is it at all frequent on this uncommon fern of basic rock crevices and rarely lime mortar.

Preston *et al*. (2002) note a decline in records from Wales of this fern, probably reflecting the loss of habitat on old walls and buildings.

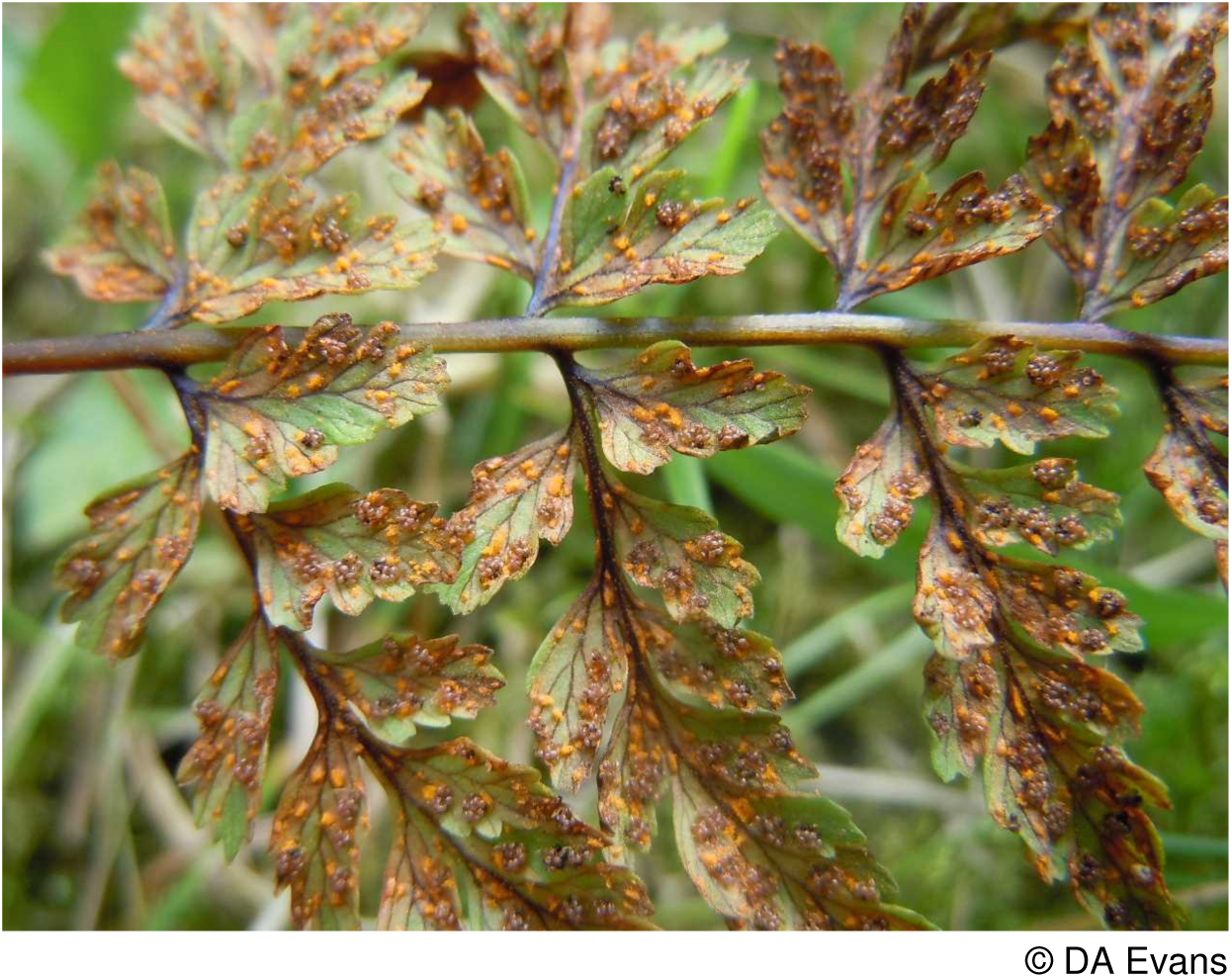

As there appear to be fewer than 20 hectad records from Wales with some evidence of decline of its host, this rust is placed in the **Near Threatened** category.

### Melampsora

This rust genus, when infecting willows and poplars and their hybrids, with or without alternating generations on various conifers, dicotyledonous and monocotyledonous plants, presents a challenge in both the accurate determination of the host, particularly where hybrids are concerned and in the complex and somewhat confused state of the taxonomy of the rusts. A single broad species concept can be adopted with a number of races distinguished based on the different alternate hosts, or as here a more restricted species concept employed. Anyone seriously considering the significance of willow and poplar rusts should consult Pei and McCracken (2005). The loss of flood plain woodland in Wales has undoubtedly reduced the populations of black-poplar *Populus nigra* and willow species such as almond willow *Salix triandra* and purple willow *S. purpurea* and their associated rusts.

The environmental assessment of the impact of re-introducing beavers must take account of their likely impact on these rust species. Other threats are noted below.

### Melampsora allii-fragilis *Kleb*

With aecia on ramsons *Allium ursinum* and uredinia and telia on crack-willow *Salix* × *fragilis* and bay willow *S. pentandra* and their hybrid, there are single recent records on ramsons on a riverbank at Pont Tanycastell, Rhydyfelin, Cardiganshire and from Llangoed in Breconshire; four records from Caernarvonshire and one from Anglesey. There are single records on crack-willow from Radnorshire and Carmarthenshire and from the latter VC on bay willow. Wilson and Henderson (1966) consider the aecia on ramsons to be indistinguishable from those caused by the next species. In central Southern England and now in Breconshire crack-willows are declining in vigour due to new non-rust fungal and bacterial infections and the continued existence of this rust on this host is clearly threatened. Given this threat and the small number of known sites this rust is considered to be **Vulnerable D2**.

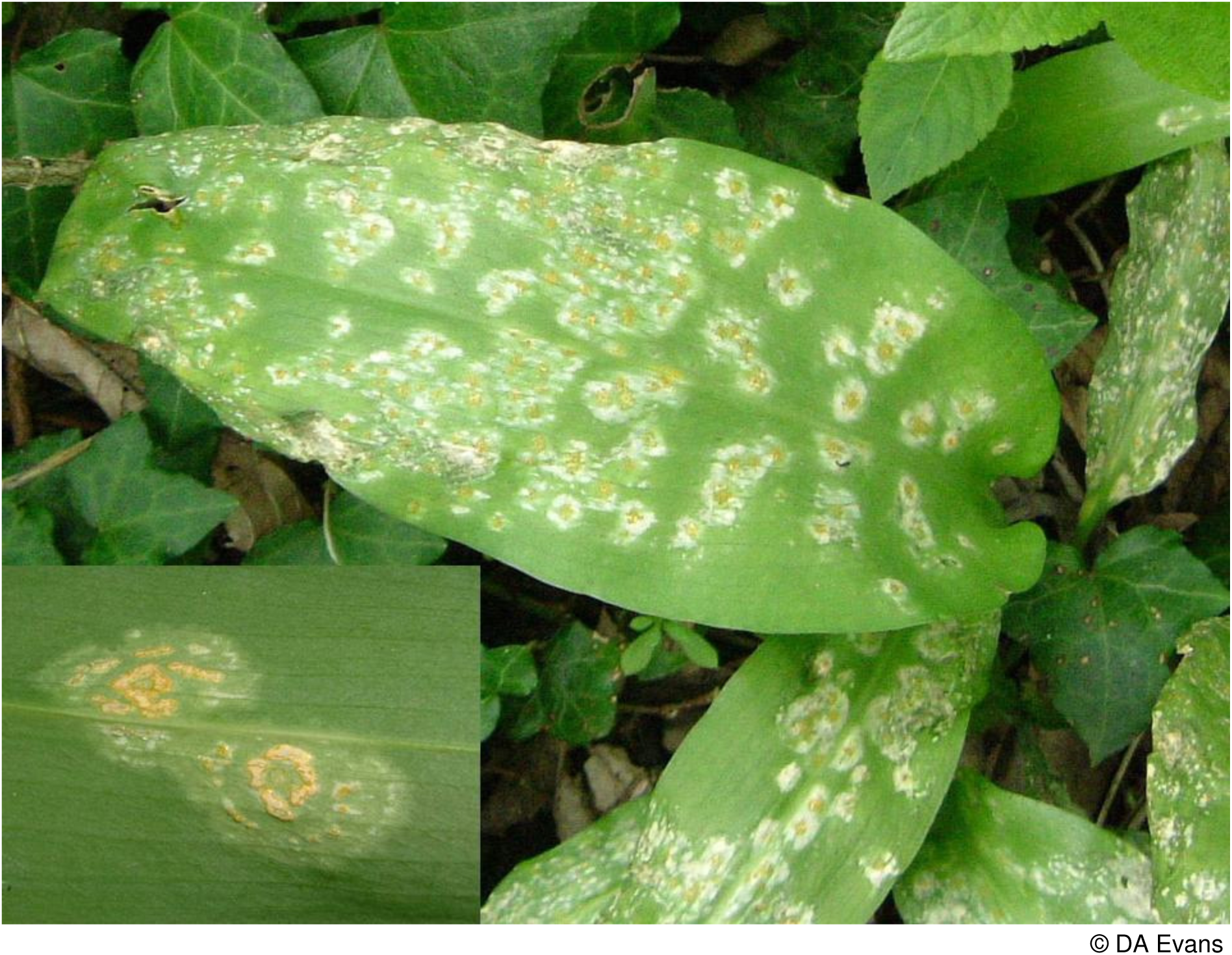
Aecia on ramsons

### Melampsora allii-populina ***Kleb*.**

There is a single British record on lords-and-ladies *Arum maculatum* from Llandeilo, Carmarthenshire and a single Welsh record on black-poplar *Populus nigra* from Monmouthshire, its alternative native host. There is also a single record from the introduced western balsam-poplar *P. trichocarpa* from Carmarthenshire. Legon and Henrici (2005) consider that records on lords-and-ladies need confirmation. Until this is forthcoming this species is noted as **Data Deficient**.

### Melampsora arctica Rostr

Recently discovered on dwarf willow *Salix herbacea* in a number of high altitude sites in Snowdonia, Caernarvonshire, including four areas on Elidir Fach, on Foel Grach summit and the Mynydd Perfedd plateau. This rust has been considered to be part of the *M. epitea* complex by some authors though all recent studies examining DNA sequences (such as the work of the Scottish Montane Willow Research Group in Edinburgh and Smith *et al*. (2004) in North America) suggest the rust on dwarf sub-arctic willows is distinctly different. It is also morphologically distinct, producing urediniospores on the upper surface of the leaves unlike *M. epitea* where they are almost exclusively on the lower surface. Outside Britain it alternates with saxifrage species *Saxifraga* spp. and should be sought on these hosts in Wales. Dwarf willow was considered by Dines (2008) to be Near Threatened in Wales. This rust species should be considered to be **Endangered D**.

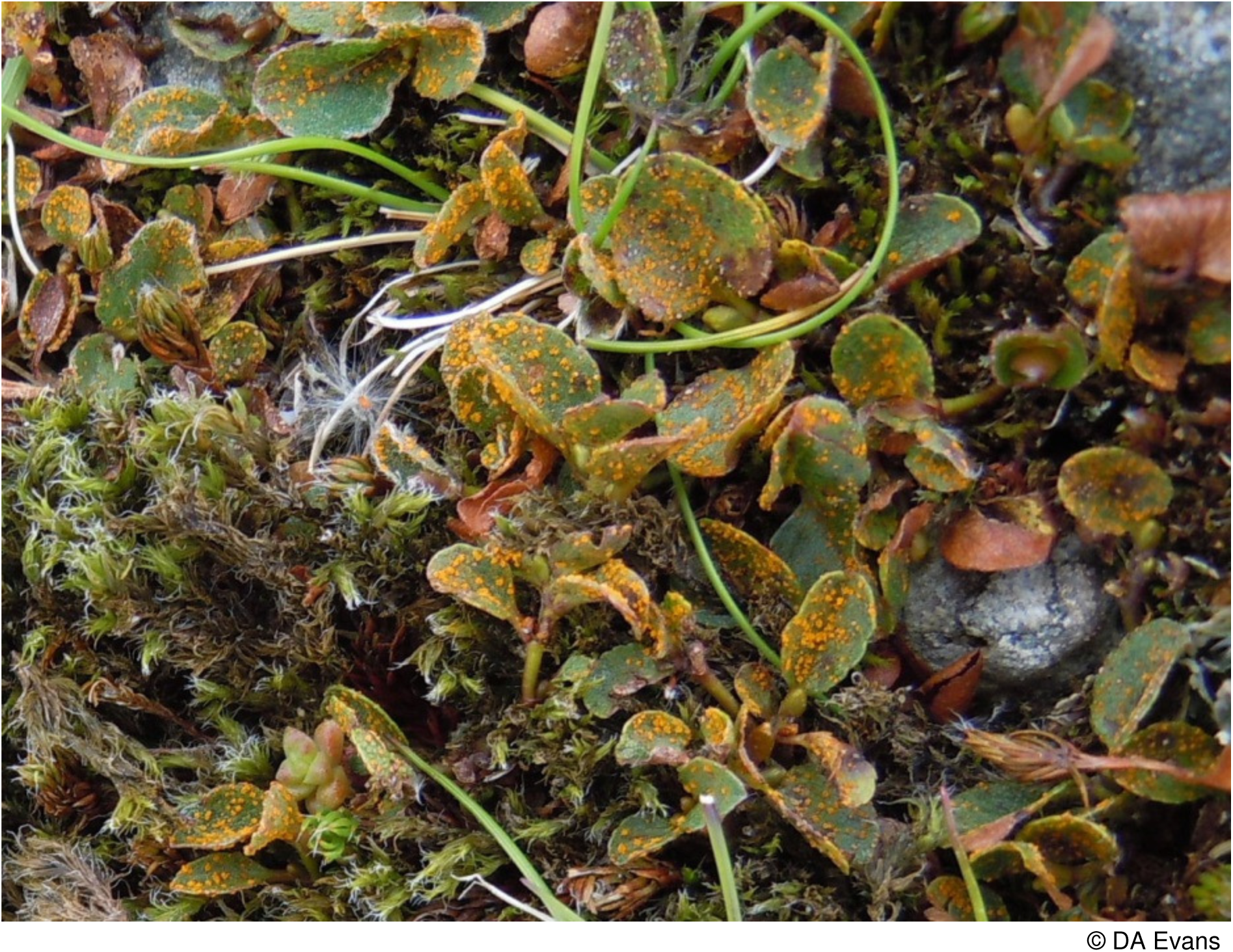
*Melampsora arctica* on dwarf willow

### Melampsora amygdalinae ***Kleb*.**

This rust species, reported from two sites in Carmarthenshire and one in Cardiganshire, is dependent on almond willow *Salix triandra* which is a rare tree in Wales. There are fewer than a dozen records of this rust from England and none from Scotland. Since the population of suitable host trees in all its known localities in Wales is likely to be less than 50 individuals and one known host tree was recently felled, this rust is considered to be **Endangered B2, D**.

### Melampsora epitea Thüm

The race on Orchidaceae alternating with creeping willow *Salix repens* and possibly eared willow *S. aurita* is sometimes distinguished by the name *M. repentis* Plowr.. It is of scarce occurrence in Wales. There are records from early marsh-orchid *Dactylorrhiza incarnata,* heath and common spotted-orchid *D. maculata* and *D. fuchsii*, northern marsh-orchid *D. purpurella*, southern marsh-orchid *D. praetermissa* and marsh and dune helleborine *Epipactis Melampsora epitea on northern marsh-orchid* *palustris* and *E. dunensis*. Particularly notable sites are the sand dune systems of Aberffraw and Newborough Warren on Anglesey. These fen-loving orchids have shown a decline at least in mid Wales (Woods 1993, Trueman *et al*. 1995 and Braithwaite *et al*. 2006). This race, if it proves to be distinct, should be considered to be Vulnerable D but for the time being is considered as **Data Deficient**.

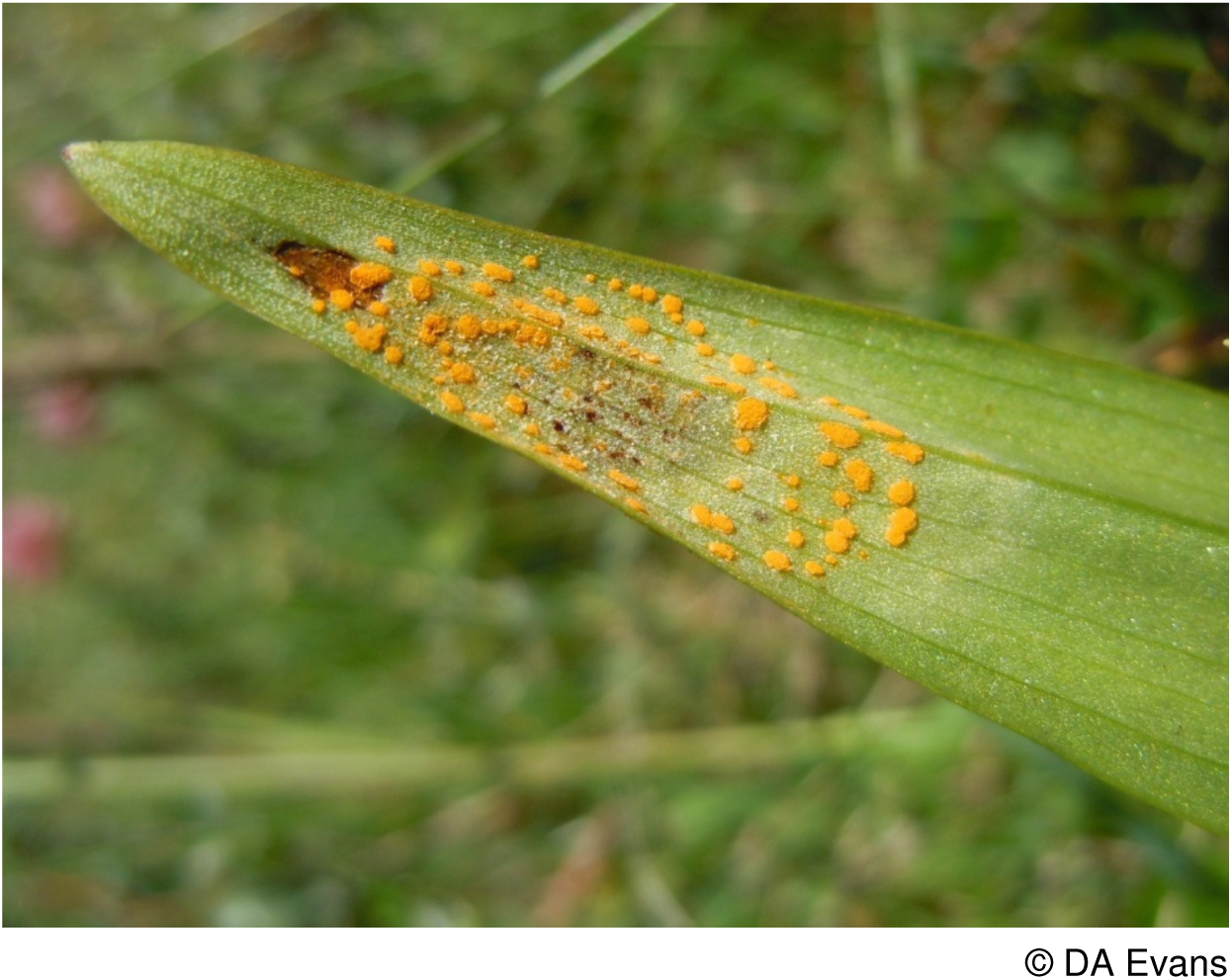

### Melampsora laricis-pentandri *Kleb*

Confined to bay willow *Salix pentandra*, populations of this rust of unknown extent occur on the Ministry of Defence Ranges on Mynydd Epynt, Breconshire, in two sites in both Monmouthshire and Carmarthenshire and one in Cardiganshire. Never common on this host, it may, however, prove to be widespread on this willow and has yet to be sought on the bay willow hybrids further north in Wales, nor has it been found on its alternate hosts of larch *Larix* spp.. For the time being it is assessed as **Data Deficient**.

### Melampsora lini var. radiolae Grove *ined*

Collected on allseed *Radiola linoides* from Aberffraw, Anglesey in 1936 by P.G.M. Rhodes, the name of this rust may never have been validly published. The specimen held in Kew is the only record of this rust on this host from Britain. Given that there are no other recent records it must be considered to be **Extinct** in Britain.

### Melampsora ribesii-viminalis *Kleb*

There are only two records from Wales on osier *Salix viminalis* and 16 from the rest of Britain (FRDBI). Recorded from Glamorgan and Carmarthenshire the sizes of the populations are unknown. Until the size can be established on this frequently occurring willow a **Data Deficient** category is proposed. This rust elsewhere alternates with currant *Ribes* species.

### Melampsora salicis-albae *Kleb*

Reported from Britain only on white willow *Salix alba* and its hybrids. There are only six records from England on the FRDBI despite the widespread occurrence of its host. Its aecia on ramsons *Allium ursinum* have not been found in the British Isles. In Wales it is reported from six sites in Carmarthenshire, two in Cardiganshire and one site in Pembrokeshire. In view of the small population, infrequency of its host and that the nearest recorded English populations are in Shropshire and Warwickshire this rust is considered to be **Near Threatened** in Wales.

### Melampsorella caryophyllacearum (DC.) J. Schröt

This rust has been found to be not infrequent in North Wales on lesser stitchwort *Stellaria graminea* with 15 monad records from Caernarvonshire and 21 monads from Anglesey. In this latter vice-county it was found once on both common chickweed *S. media* and greater stitchwort *S. holostea.* In the rest of Wales, despite the widespread occurrence of the host species this rust has recently only been reported once on common mouse-ear *Cerastium fontanum* ssp. *holosteoides* (from Pembrokeshire) and once on lesser stitchwort *Stellaria graminea* (in Cardiganshire) in the last 50 years. There is an old record of Vize (1882) who notes its presence in Montgomeryshire (host unspecified). In all of the rest of Britain there are scarcely over 20 records on native plants in this same period (FRDBI). In view of the limited extent of the population with records from only 14 hectads and the likely loss of dry grassland sites supporting its major host, lesser stitchwort (Stevens *et al*. 2010) it is considered to be **Near Threatened**.

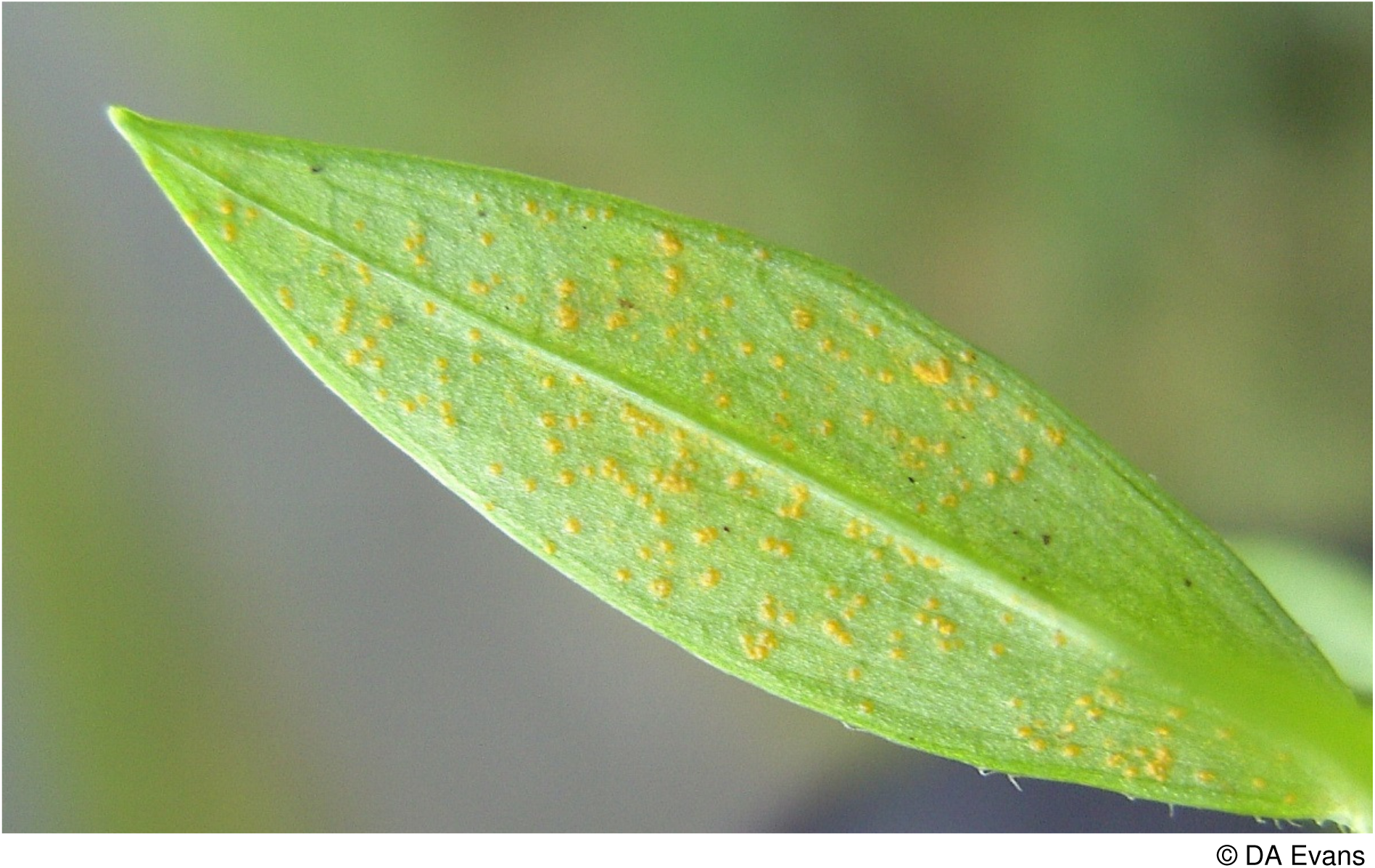
Melampsorella caryophyllacearum on lesser stitchwort

### ***Milesina carpatorum*** Hyl., Jørst, & Nannf

Found on the fronds of male-fern *Dryopteris filix-mas,* it is unfortunately impossible to separate in the field from the common *M. kriegeriana*. Evans *et al*. (2006) considered this species to be Vulnerable within Britain. There have, however, been over thirty recent records from southern England since this evaluation was made and this species might now be better considered to be of Least Concern at a British scale. In Wales, only two records have ever been made despite investigating hundreds of collections. It was found at Derwydd, Carmarthenshire in 2000 (FRDBI) and at Gamlyn Bridge, Aber-ffrwd, Cardiganshire in 2013. The discoveries in England should stimulate further searches in Wales. Given the intensive searches with little success this rust is for the time being considered to be **Critically Endangered D2** in Wales.

### Milesina vogesiaca Syd. & P. Syd

Rarely reported in Britain on hard shield-fern *Polystichum aculeatum* and even more rarely on soft shield-fern *P. setiferum* and their hybrid. Evans *et al*. (2006) considered its status to be Vulnerable D2 but subsequent to this evaluation a number of new records have appeared on the FRDBI from N. Somerset, Dorset, S. Wiltshire and SW Yorkshire and this evaluation requires revision. Nick Legon, in particular, has made a number of these records and helpfully notes that the uredinia are rather distinctive in being a very bright reddish brown in colour and are hence easily visible and separate it from the much commoner white uredinia of *M. whitei*. In Wales it is reported from three sites in Carmarthenshire on hard shield-fern and it should be sought elsewhere amongst large populations of this fern. Given the small population size this rust is considered to be **Vulnerable D1 & D2** in Wales.

### Ochropsora ariae (Fuckel) Ramsb

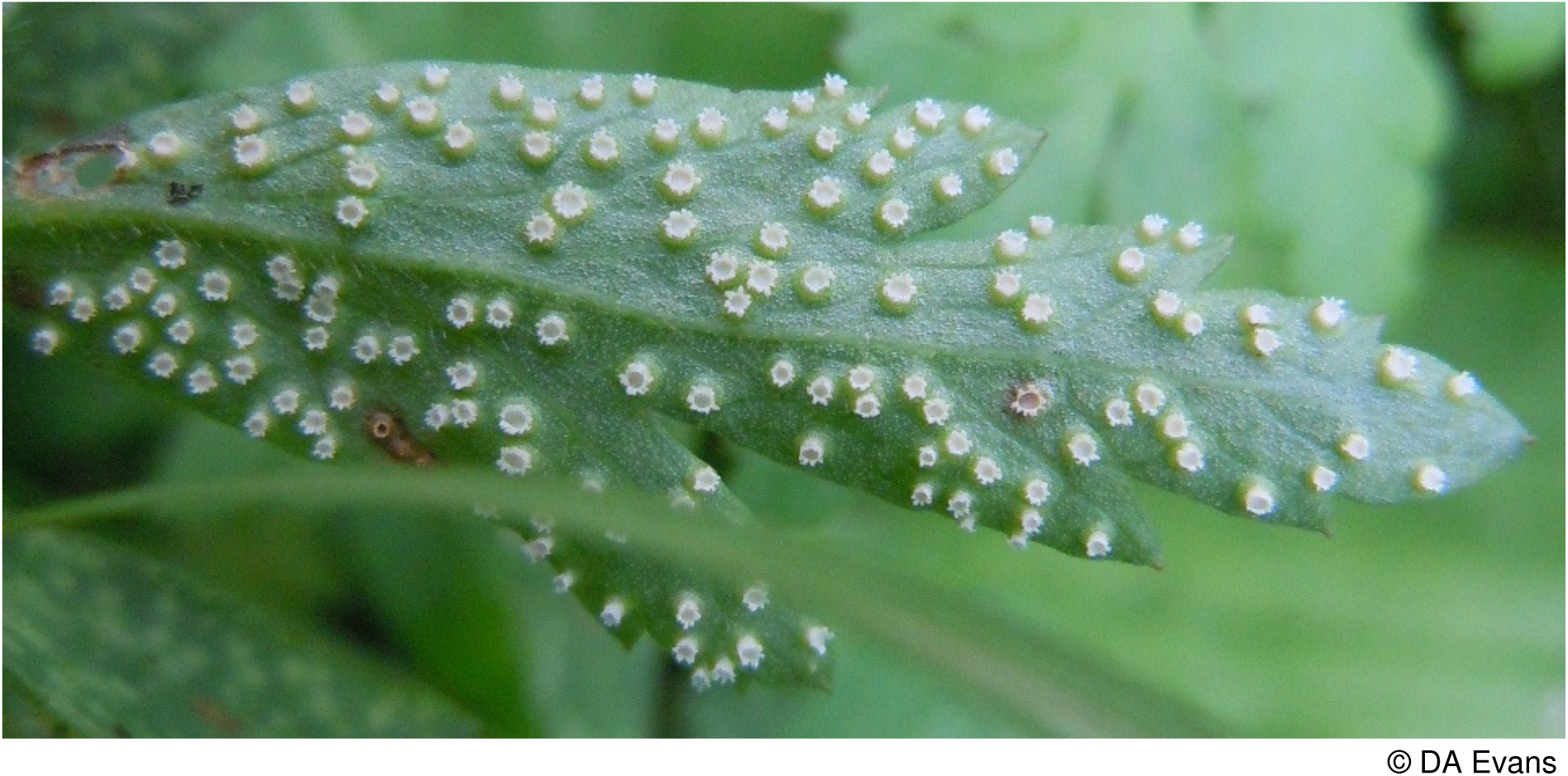

The taller leaves with narrower leaflets with scattered white aecia on the underside quickly separate infected wood anemone *Anemone nemorosa* leaves from normal leaves. There are 17 records from Wales of this scarce rust. Despite large populations of wood anemone in some sites infected leaves are few in number. It has never been reported on its alternative host rowan *Sorbus aucuparia*. In the absence of any detailed population data a status of **Near Threatened** is considered appropriate.

### Phragmidium potentillae (Pers.) P. Karst

There are seven Welsh records of this rust on trailing tormentil *Potentilla anglica*, a single record from Breconshire and three each from Carmarthenshire and Cardiganshire; and a single record on the hybrid *P*. × *suberecta* from Breconshire. The FRDBI lists only seven British records made in the last 50 years so the rust is clearly nationally scarce. An understanding of its true distribution is hampered by the regular occurrence of hybrids of the host species and the difficulty of separating this rust species from *Frommeëlla potentillae*. Given these problems for the present it is considered to be **Data Deficient**.

### Puccinia angelicae (Schumach.) Fuckel

Reported recently on wild angelica *Angelica sylvestris* from six high quality nature conservation sites in Wales viz. in mixed woodland at Llethrid, Glamorgan; on sand dunes at Tywyn Burrows, Carmarthenshire; the edge of a raised mire, Cors Caron, Tregaron, Cardiganshire; and on sand dunes at Newborough Warren (in 4 monads) and in fens on Cors Erddreiniog and at Cors Bodeilio, all in Anglesey. There are records from the early 1900’s on this host near Monmouth and Swallow Falls, Betws y Coed and recently from less than a dozen sites in Scotland and a single site in England. It is also reported from six sites on milk-parsley *Thyselium palustre* (= *Peucedanum palustre*) in eastern England and from a single site in Worcestershire on pepper-saxifrage *Silaum silaus* (FRDBI). This latter species was once more widespread in eastern Wales but wet pastures have largely been drained and its habitat lost. In view of this decline and the small number of sites this rust is placed in the **Endangered D** category.

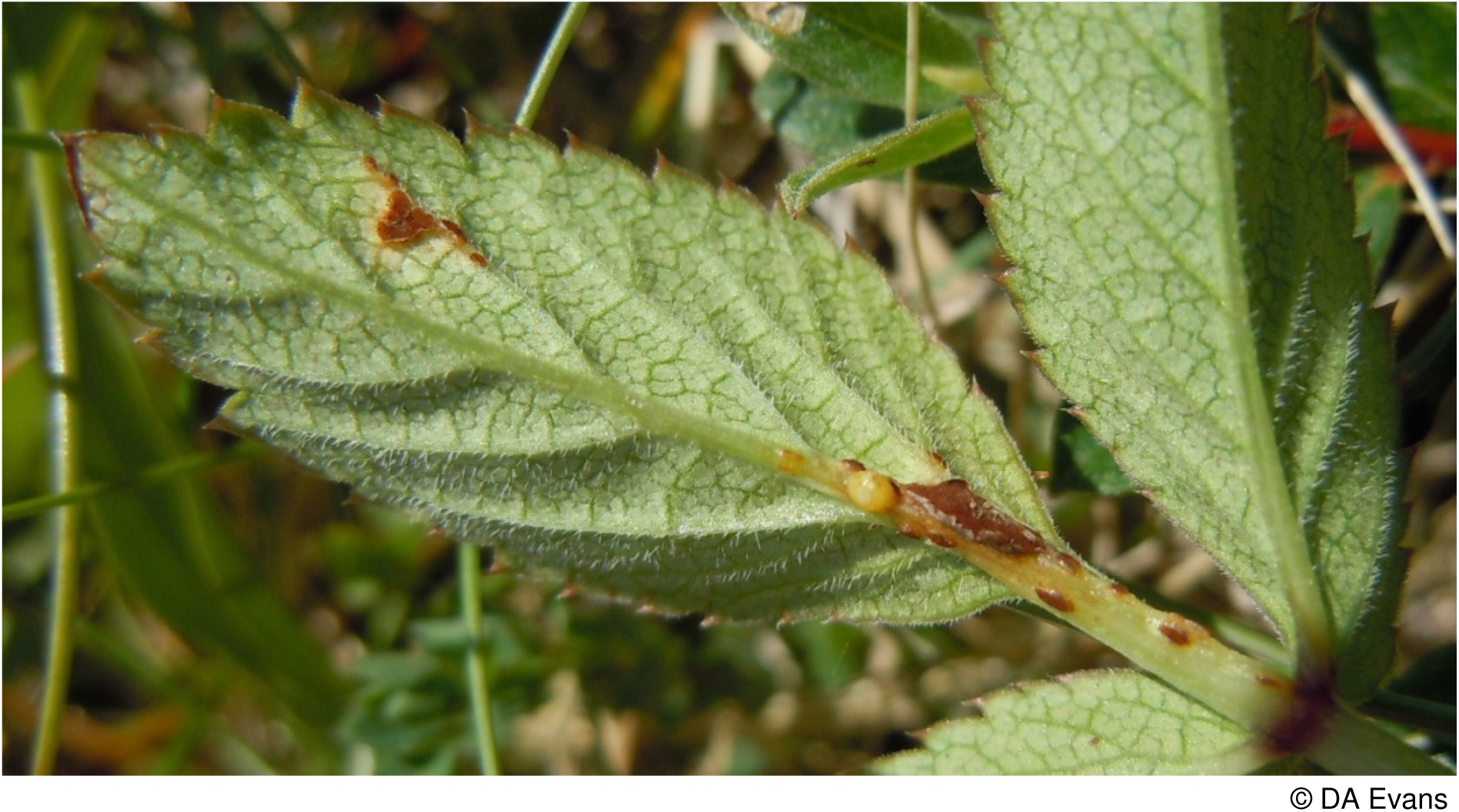

### Puccinia apii Desm

A nationally rare rust confined to wild celery *Apium graveolens*. This host is almost exclusively coastal in Wales. The FRDBI notes a record from Prestatyn in Flintshire in 1913. There are no more recent records from Wales and only two recent records from eastern England. It is considered to be **Regionally Extinct**.

*Puccinia bistortae* (F. Strauss) DC.

A nationally scarce rust with aecia on either pignut *Conopodium majus* or cow parsley *Anthriscus sylvestris* and uredinia and telia on bistort *Persicaria bistorta*. Mostly northern in distribution in Britain, there is a specimen on pignut collected in 1960 in Penglais Woods, Aberystwyth, Cardiganshire stored in the fungarium at Kew and a more recent record by Paul Smith in 2011 on pignut and bistort from the Memorial Park Meadows, Pontllanfraith, Monmouthshire. Due to the small size of its population with no nearby records from England it is considered to be **Critically Endangered**.

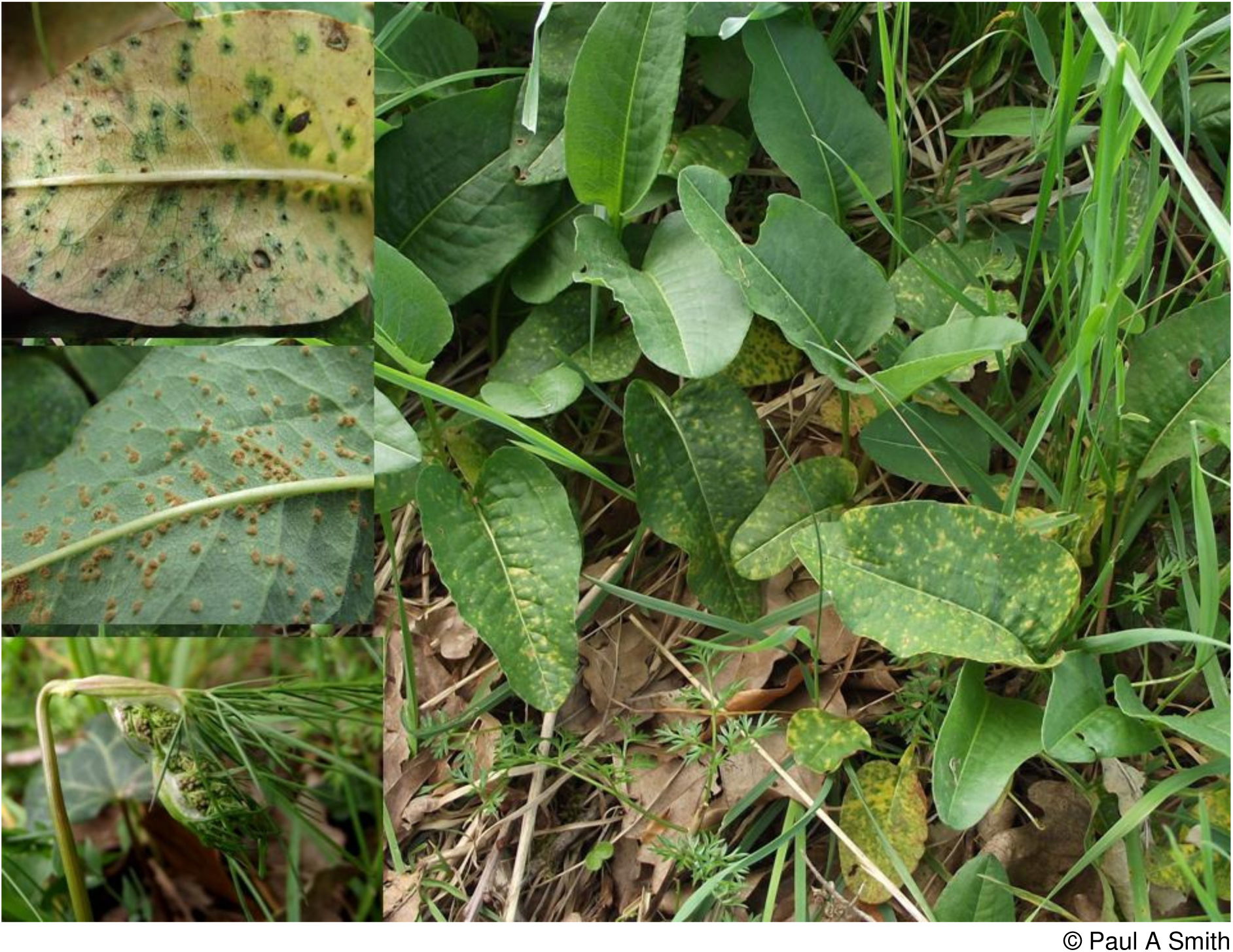
Aecia on pignut (bottom left). Uredinia (mid left) and telia (top left) on bistort (right).

### Puccinia calthae ***Link***

Found on marsh-marigold *Caltha palustris,* there are recent records of this rust from 14 hectads in Wales, mostly from high conservation quality wetlands. It appears to be less frequent than *P. calthicola,* a feature shared with England, where the FRDBI lists only a handful of sites in the last 50 years. In contrast there are 26 records from Scotland (20 in

FRDBI and 6 additions of Paul Smith pers. comm.) where this rust is more frequent, particularly in the west. Lost from at least one site in Wales recently due to the spread of woody species and given the widespread drainage of wet woodlands and pastures in the last 50 years, its host has suffered a decline (Braithwaite *et al*. 2006). There are, however, a few additional records of rusts on this host that could not be certainly separated from the next species. For the time being it is placed in the **Near Threatened** category.

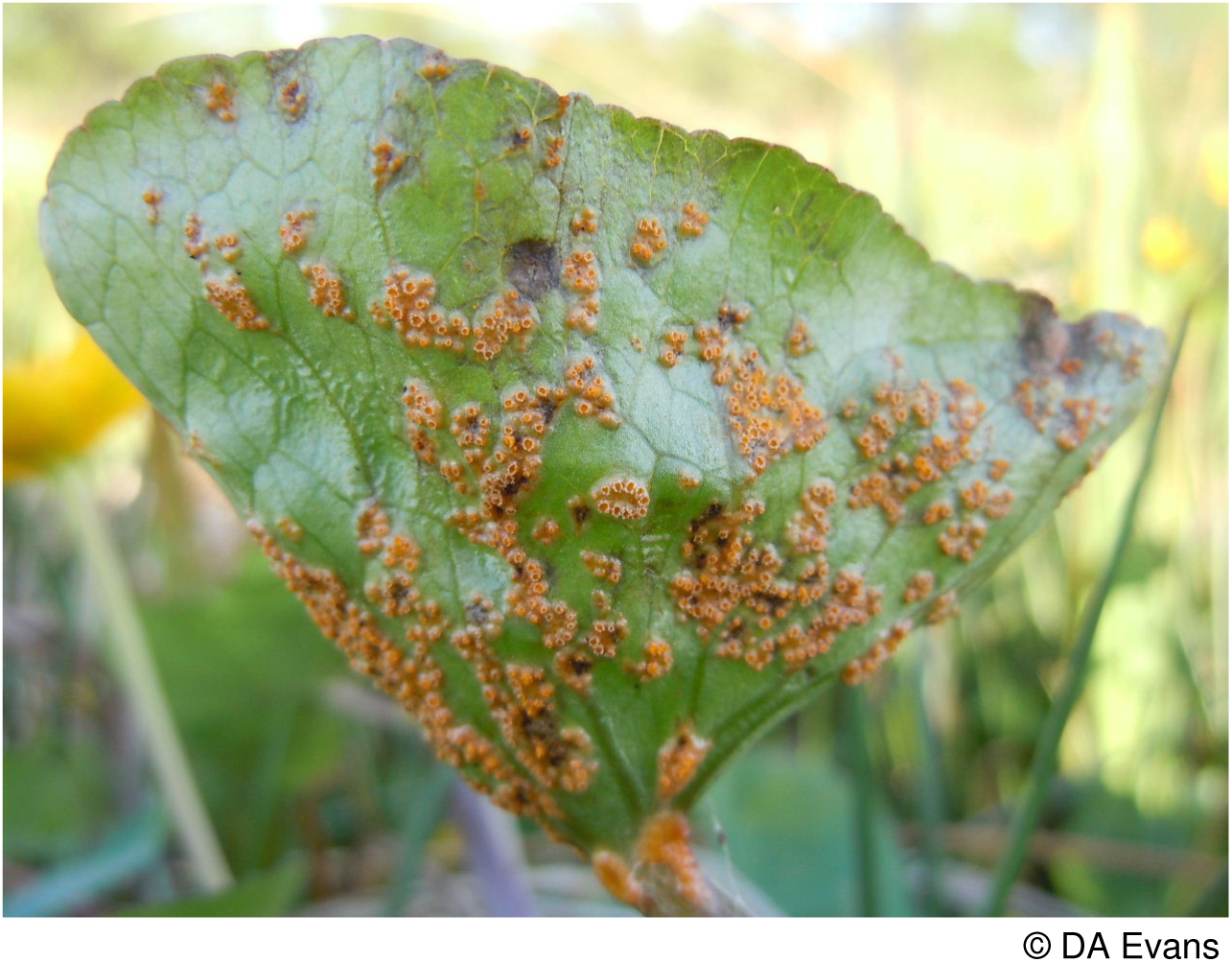
Puccinia calthae

### Puccinia calthicola J. Schröt

Found on marsh-marigold *Caltha palustris* in mostly high conservation quality wetlands like the last species. The FRDBI lists seven records from England and one from Scotland in the last 50 years, though Paul Smith has recently made 16 records from the Outer Hebrides (pers. comm.). Within Wales it is recorded from seven sites in Carmarthenshire, one in Pembrokeshire (pre 1964) and Montgomeryshire and eight sites in both Caernarvonshire and Anglesey and two each in Breconshire, Radnorshire and Cardiganshire. As with *P. calthae,* due to the widespread drainage of wet woodlands and pastures in the last 50 years, its host species has suffered a decline (Braithwaite *et al*. 2006). Notwithstanding this threat, since it occurs in more than 20 hectads in Wales, it is considered to be of **Least Concern**.

### Puccinia campanulae Carmich

Legon and Henrici (2005) note the lack of confirmed records from Britain since a specimen was collected in England in 1922. Aron (2005) however notes records from harebell *Campanula rotundifolia* at Coedydd Aber, Caernarvonshire in 1979 and on the same host on limestone at Parciau on Anglesey in 1999. There had been an earlier record on this host from Betws y Coed in Caernarvonshire in 1866 and on sheep’s-bit *Jasione montana* from near Lampeter, Cardiganshire in the 1840’s (FRDBI). Confirmation of these recent Welsh records is desirable. If confirmed this rust should be categorized as Critically Endangered D2. Until these N. Wales records are confirmed a **Data Deficient** status is proposed.

### ***Puccinia cancellata*** (Durieu & Mont.) Sacc. & Roum

Until recently this rust was only known from the uncommon sharp rush *Juncus acutus*, a distinctive rush of sand dunes. Welsh populations of this rust may be nationally significant. Over half the British population of this rush occurs in Wales (Dines 2008). The FRDBI only lists a single record from England (Braunton Burrows in Devon in 1959) and a number of records from the Channel Isles. Recent searches in Wales have located significant populations in both Glamorgan and Carmarthenshire with an isolated population in Merionethshire. What appears to be this rust has been found on sea rush *Juncus maritimus* from a number of sites in both Glamorgan and Carmarthenshire and also a large population on the Dyfi Estuary in Cardiganshire where it was first found by Arthur Chater in 2002.

Large patches of sea rush become infected in contrast to the infection pattern on sharp rush where it occurs as isolated individuals. No records of this rust on sea rush have been found in the literature and confirmation by either DNA analysis or cross-infection experiments is required before these records can be confidently accepted. For the moment the status of this rust in Wales is considered to be **Data Deficient**.

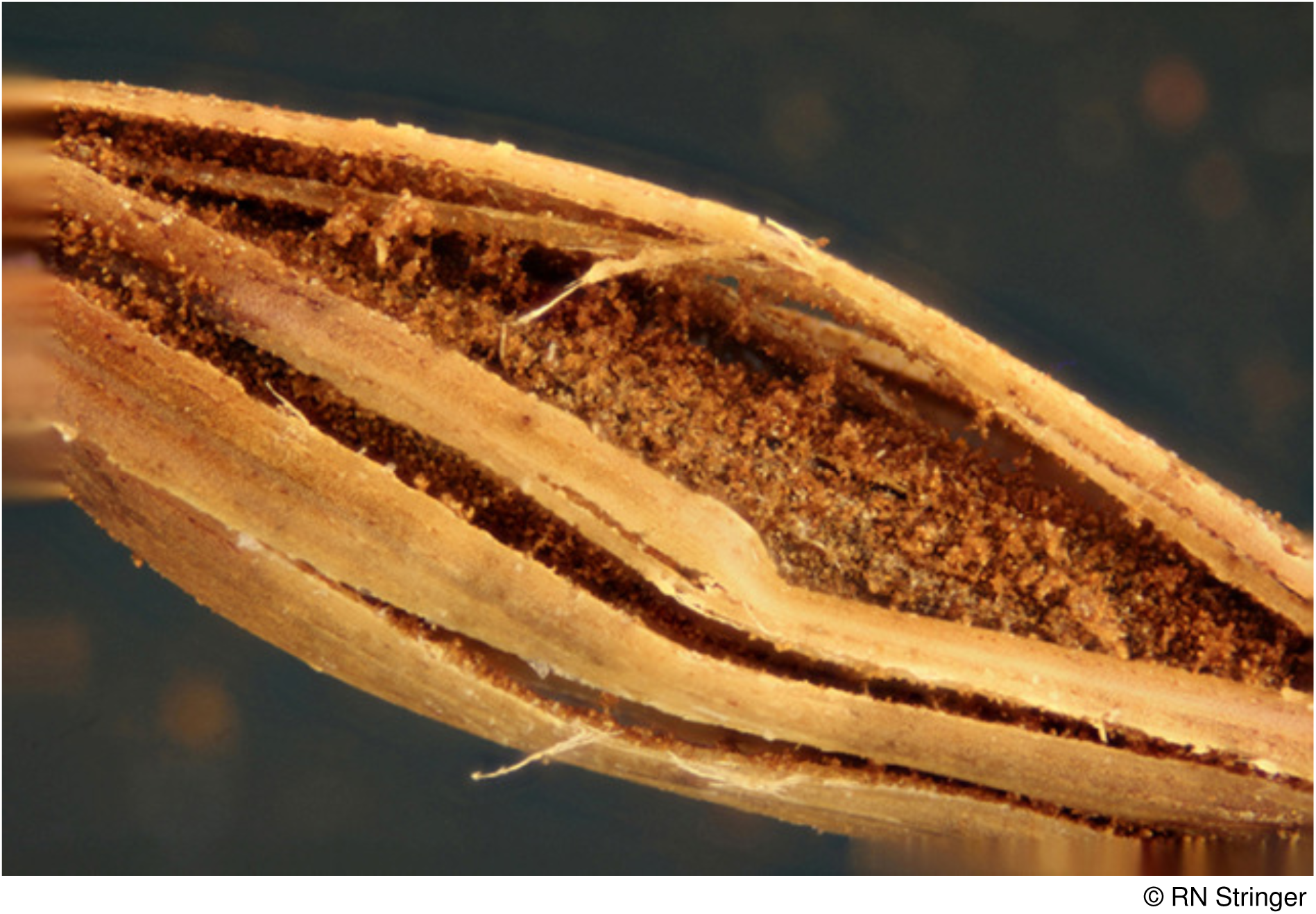
*Puccinia cancellata* on sea rush

### Puccinia caricina var. magnusii (Kleb.) D.M. Hend

Uredinia and telia are produced on greater pond-sedge *Carex riparia*, a sedge that is very much a lowland, coastal species in Wales. This rust is known from one site in Monmouthshire, five sites in Carmarthenshire, one site in Pembrokeshire, three sites in Cardiganshire and one site in Anglesey. In England it is only known from nine sites on this sedge. There is, in addition, a single record from Derbyshire of the aecia on mountain currant *Ribes alpinum*. On the continent this rust is recorded from a range of *Ribes* species.

Whether some of the records on *Ribes* ascribed to the next variety belong here is not known. For the time being it is considered to be **Near Threatened** in Wales.

### Puccinia caricina var. pringsheimiana (Kleb.) D.M. Hend

Identified with certainty only when it occurs on slender tufted-sedge *Carex acuta*, itself a rather scarce sedge of the major rivers in Wales. It is recorded from single sites in Breconshire, Radnorshire and on Anglesey. Aecia are found on *Ribes* species. On gooseberry *R. uva-crispa* there are records from Carmarthenshire, Caernarvonshire and Anglesey which are claimed as this variety. Whether the records on this host from Breconshire and Merioneth ascribed to *P. caricina s.l*. are of this variety is uncertain since the var. *magnusii* is said to alternate with *Ribes* on the continent (Wilson and Henderson 1966). This rust also lives on common sedge *Carex nigra* but cannot be distinguished by conventional means on this host. The population size of this rust therefore remains uncertain and it is placed in the **Data Deficient** category. Should it prove to be confined to tufted-sedge it would be considered to be at least Near Threatened.

### Puccinia caricina var. ribis-nigri-paniculatae (Kleb.) D.M. Hend

Known in Wales only from the greater tussock-sedge *Carex paniculata*. Despite Wilson and Henderson’s (1966) assertion that it is probably present wherever its host occurs in quantity, there are only 24 records, from fewer than 20 hectads in Wales and 25 on the FRDBI from the British Isles. The authors have noted the drainage of numerous wet flushes and wet woodlands in the last 40 years in Wales with the consequent loss of this sedge and both Braithwaite *et al*. (2006) and Preston *et al*. (2002) also note a decline. It is considered **Near Threatened** in Wales.

### Puccinia chaerophylli Purton

Providing a conservation evaluation for this species presents something of a dilemma. It is frequent on the introduced and now widely naturalised pot herb sweet cicely *Myrrhis odorata* with over a dozen records. On native Apiaceae it is a different story with just three recent records from Wales on cow parsley *Anthriscus sylvestris*, (on a hedgebank at Glynhir and at Llanddowror in Carmarthenshire and a single record from Cardiganshire, all in 2014).

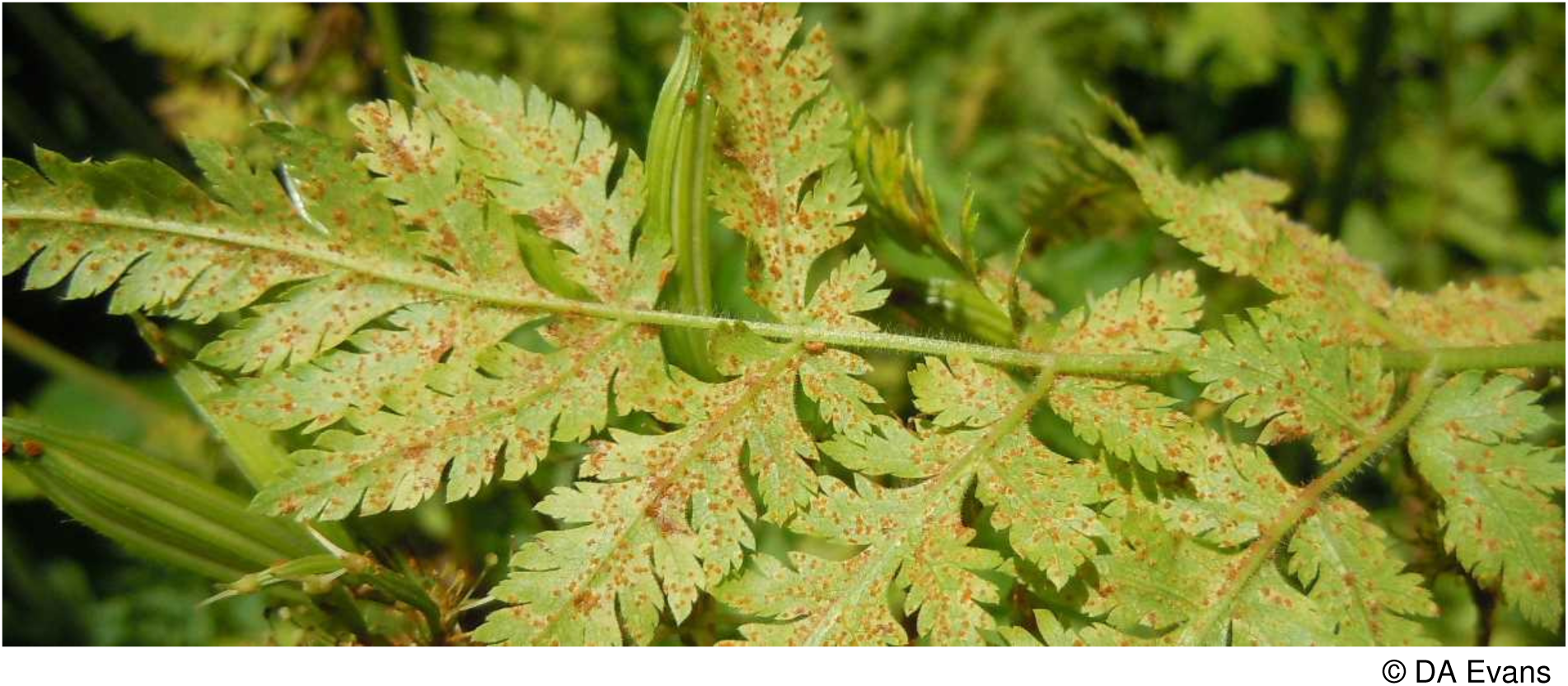
*Puccinia chaerophylli* on sweet cicely

There are old records from Newborough Warren, Anglesey in 1950 and 1956. It has been found on rough chervil *Chaerophyllum temulentum* at Bwlch in Breconshire in 1988; and at Penallt Farm in 2006 and at Llanwrda in 2002 and 2005, both sites being in Carmarthenshire. In England it is equally scarce on species other than sweet cicely, though Tom Preece notes it on cow parsley close to the Montgomeryshire border in Shropshire.

Within Mid Wales a recent expansion of the rust on sweet cicely has been noted. This has not been reflected in records on cow parsley, a species that appears to have become increasingly abundant on road verges. Different races may be involved on these two hosts. Until this is resolved a conservation evaluation of **Data Deficient** is proposed.

### Puccinia cladii Ellis & Tracy

The FRDBI has no records post 1958 and Evans *et al*. (2006) have in consequence considered this species to be extinct in Britain. It does, however, survive on great fen-sedge *Cladium mariscus* in the fen of Crymlyn Bog near Jersey Marine, Swansea, Glamorgan.

Threatened with destruction in the past this site is now a National Nature Reserve. Elsewhere this rust has been recorded in the past from the fens of eastern England and a recent record from Norfolk is reported (M. Yeo pers. comm.). A search of the few populations of this sedge in Central Wales and the fens of Anglesey has failed to find any additional sites for this rust. Due to the small size and isolation of its population and rarity in England it is considered **Critically Endangered** D.

### Puccinia commutata Syd. & P. Syd

The distinctive orange aecia on stems and leaves of common valerian *Valeriana officinalis* attract attention and where present are unlikely to be missed. Yet for many years this rust was only known from two sites in west and central Scotland (Wilson & Henderson 1966). In the 1980’s records were made in Cheshire, Lancashire and Cornwall but still the FRDBI notes fewer than a dozen records from England and Scotland. More recently this rust has been recorded from eight sites in Cardiganshire, and in single localities in Glamorgan, Pembrokeshire and Radnorshire and from a single site in Caernarvonshire and four sites on Anglesey. Most sites are of high nature conservation value and the possibility must exist that this rust is spreading. It should also be noted that it has recently been found infecting red valerian *Centranthus ruber* in sites along the southern English coast. Some of its Welsh sites have of late ceased to be grazed and colonisation by willow and birch may threaten its host. Placing this rust in the **Near Threatened** category will hopefully highlight the need for appropriate management.

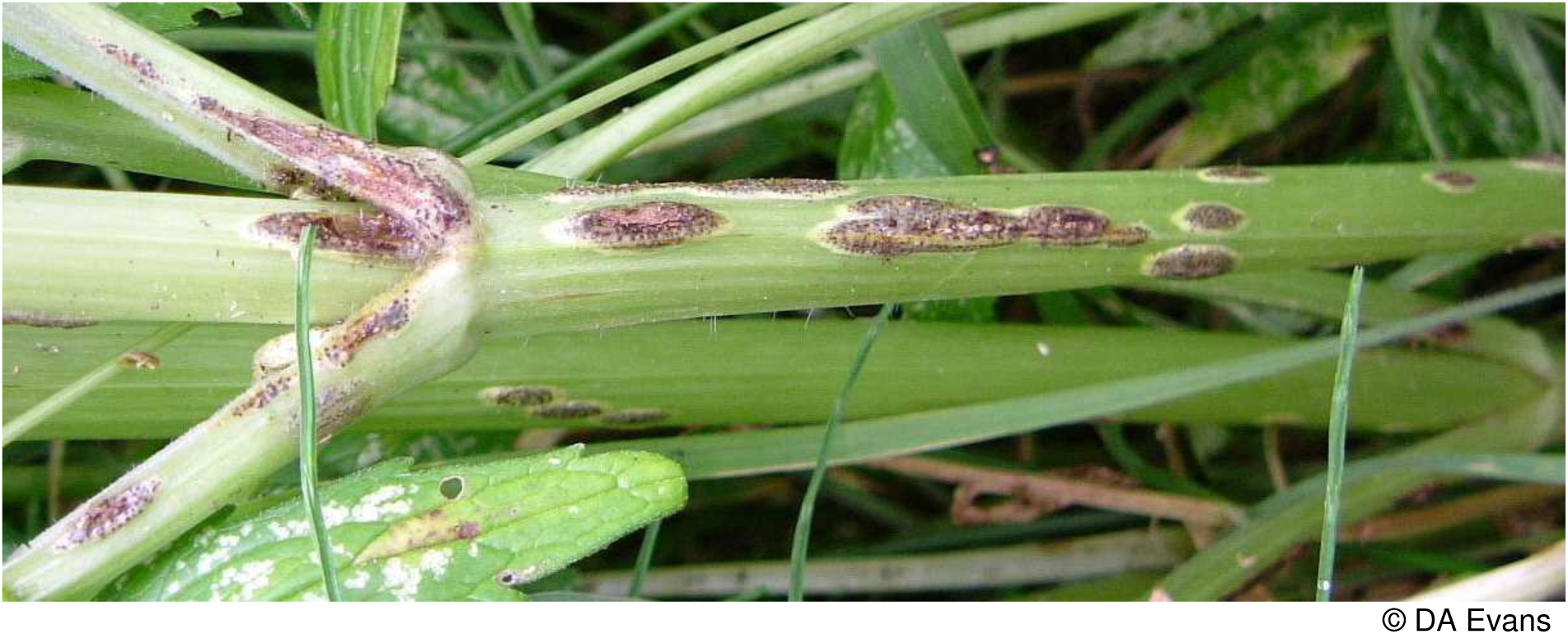

### ***Puccinia conii*** (F. Strauss) Fuckel

A parasite of hemlock *Conium maculatum*, a plant mostly confined to the coastal plains of north and south Wales, it has only been seen recently in Caernarvonshire and Anglesey where it is known from about 6 sites and from three sites in the Usk Valley between Abergavenny and Usk in Monmouthshire. The FRDBI lists only three records from Wales out of over 200 from the rest of Britain viz. collected by the Rev. Vize of Forden in Montgomeryshire in the 1800s; from Aber Castle in Pembrokeshire in 1935 and from the sand dunes of Merthyr Mawr in Glamorgan in 1955. This rust is commoner in the West Midlands and southern England where its host is more frequent. Occupancy of fewer than ten hectads in Wales, with clear evidence of decline, places this rust in the Vulnerable B category. Its population is, however, closely linked to that in England so its threat status is demoted to **Near Threatened**.

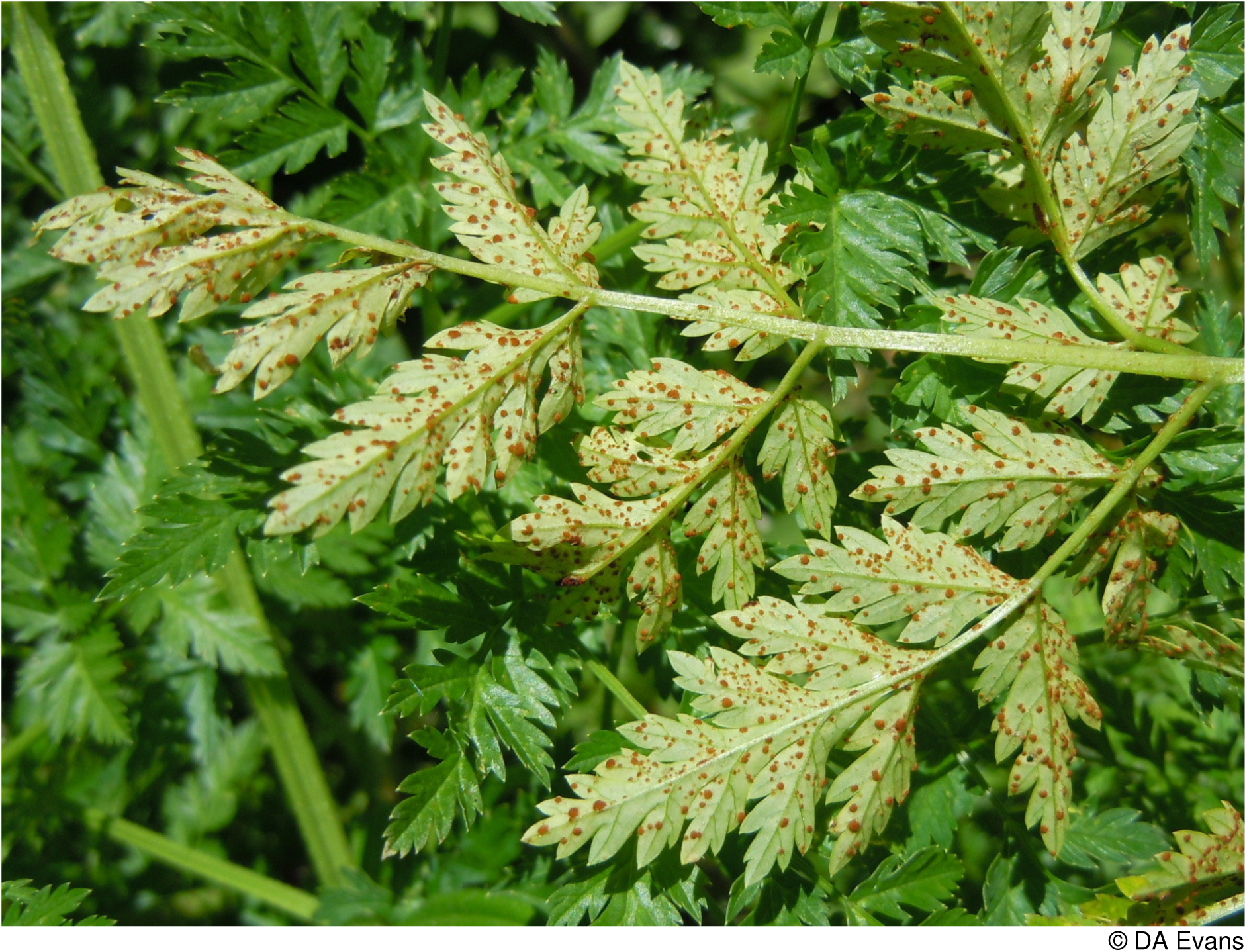

### Puccinia coronata Corda

This complex of taxa, following DNA sequencing, is currently undergoing revision. Liu and Hambleton (2013) support previous workers who considered the rust on wood fescue *Festuca altissima* to be a distinct taxon called *P. coronata* var. *gibberosa* (Lagerh.) Jørst. The sole British records are from a tiny population of this host in the Bach Howy ravine, Radnorshire and in the gorge at Devil’s Bridge, Cardiganshire. Due to the small population size, this taxon is considered to be **Critically Endangered D2**.

### Puccinia crepidicola Syd. & P. Syd

The small brown uredinia and tiny black telia are probably easily overlooked on rough hawk’s-beard *Crepis biennis,* smooth hawk’s-beard *C. capillaris* (see image below) and beaked hawk’s-beard *C. vesicaria.* There are, however, fewer than 60 recent records from the British Isles. In Wales it has recently been seen once on *C. biennis* (in Bangor, Caernarvonshire), eight times on *Crepis capillaris* in a scatter of Mid and N. Wales VCs and once in an urban habitat in SE Wales and five times on *C. vesicaria* in Carmarthenshire where it occurred on mostly open ground close to the coast and on Anglesey. Given the level of destruction of lowland flower-rich grassland and only 14 records from Wales this species is considered to be **Near Threatened**.

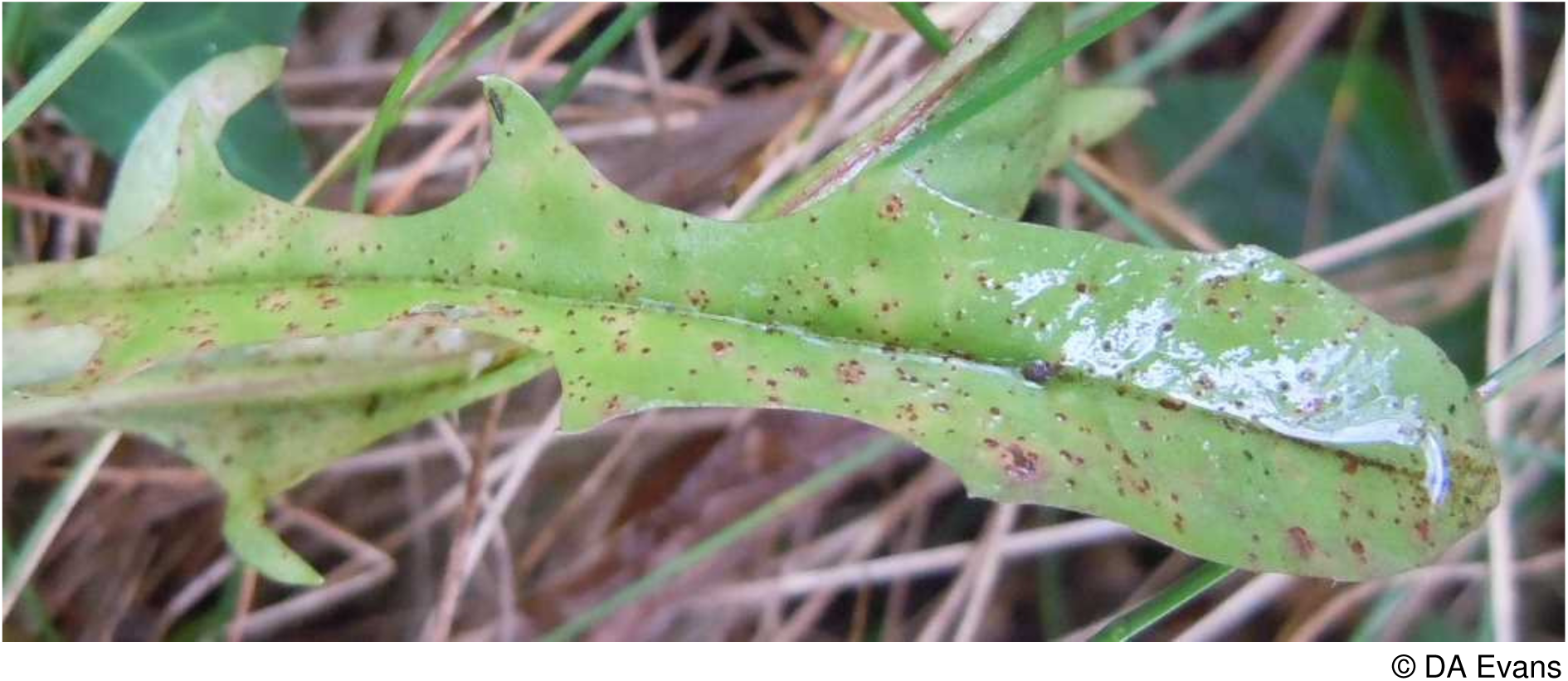
*Puccinia crepidicola* on smooth hawk’s-beard.

### Puccinia difformis J. Kunze

Given the enormous populations of its host, cleavers *Galium aparine,* the scarcity of this rust is somewhat unexpected. The aecia occur as yellow spots all over the leaf whilst the telia are said to resemble little spots of tar. In Wales it has been recorded recently from single sites in Glamorgan, Carmarthenshire, Merionethshire and Denbighshire, six sites in

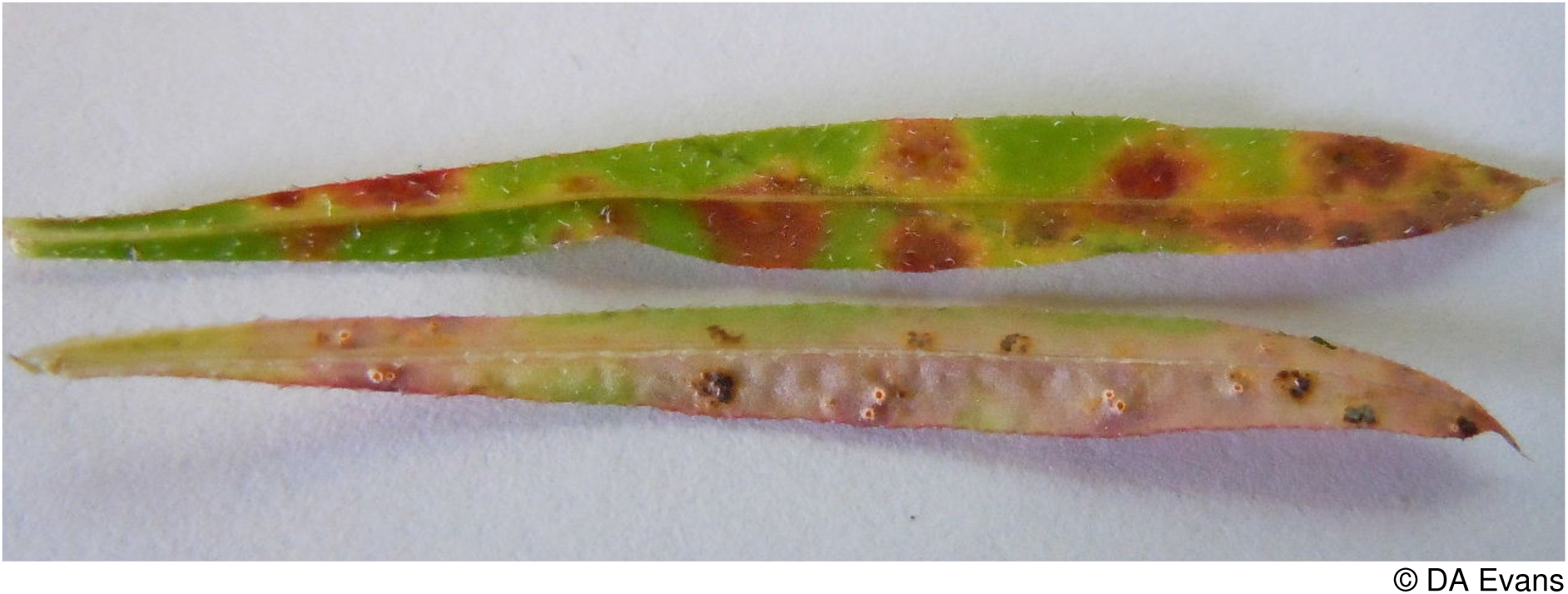

Cardiganshire, three sites in Caernarvonshire, and four sites on Anglesey. There is a specimen “on/with *Galium verum*” from Barrie Sands, Glamorgan collected by W. Gardiner in the 1840’s. Of the fifteen recent records on the FRDBI only one is from Scotland. The rest are mostly from SW England (including Herefordshire) with a number reported from river banks and fens. As it occurs in fewer than 20 hectads it is considered to be **Near Threatened** in Wales.

### Puccinia dioicae Magnus

Reported only twice in Wales on dioecious sedge *Carex dioica* from wet flushes in Breconshire and Caernarvonshire. It alternates with marsh thistle *Cirsium*

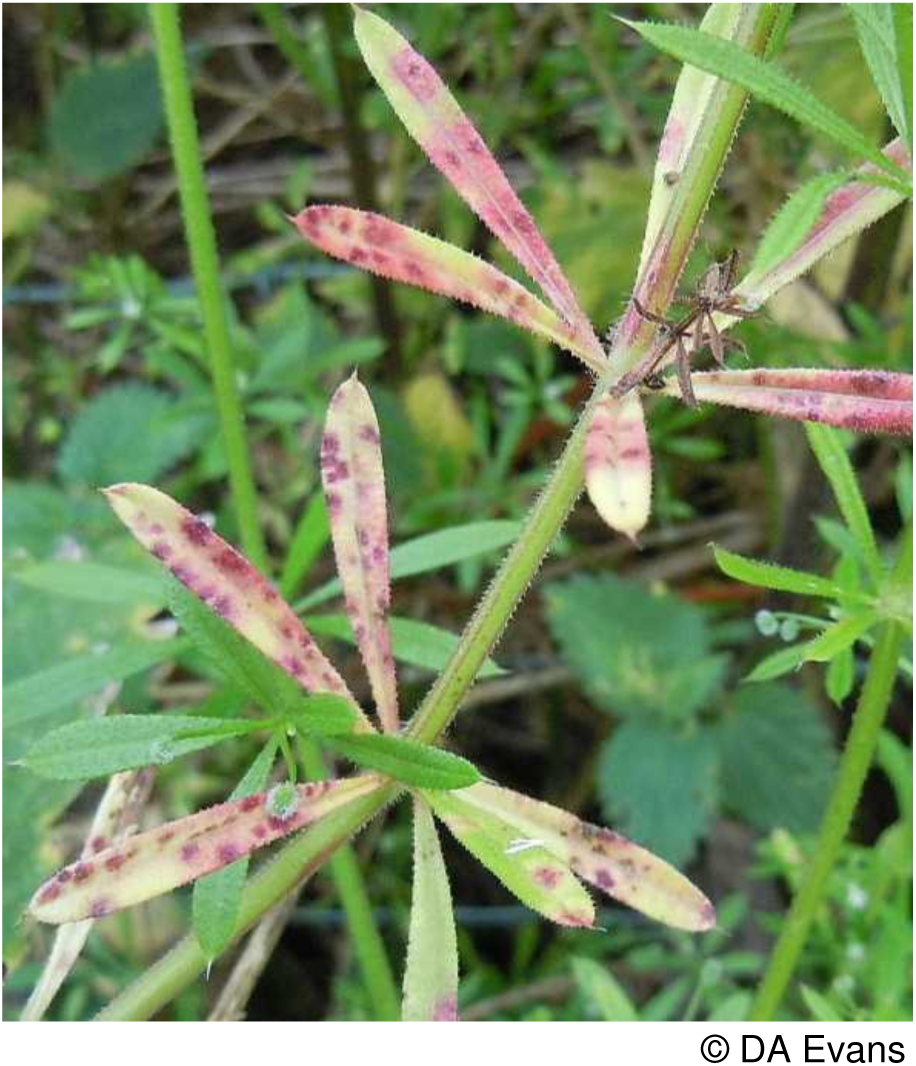

*palustre* and meadow thistle *C. dissectum*. Marsh thistle was recently found infected near dioecious sedge in Caernarvonshire (see inside rear cover for image). Three more sites have been found in this VC with infected marsh thistle and one further site in Merionethshire. The FRDBI provides a confused picture and clearly includes records of other varieties of the rust. It lists two records on dioecious sedge: one from Ben Lawers in 1959 and one from the Isle of Skye at an unknown date in the 20^th^ century. It also lists single records on meadow thistle from the North of England and on marsh thistle on Orkney. Paul Smith (pers. comm.) in his surveying in the north of Scotland has added substantially to these records recently with two records from marsh thistle, one record from dioecious sedge and three from common yellow-sedge *Carex demissa*. Detecting rust fungi on the small setaceous leaves of dioecious sedge is not easy. Searching marsh thistles close to wet flushes with dioecious sedge has proved more successful. Whilst accepting this problem of possible under-recording, the paucity of records on thistle hosts and the general scarcity of large populations of this sedge in Wales together with a paucity of records from elsewhere in Britain does point to the probability of this rust being rare and a categorisation of **Endangered D** is proposed based on the tiny known population size.

### *Puccinia dioicae* var. *schoeleriana* (Plowr. & Magnus) D.M. Hend

Orange cup-like aecia occur on common ragwort *Senecio jacobaea* and telia on sand sedge *Carex arenaria*. The FRDBI has 10 recent records from Britain; eight on common ragwort and two on sand sedge. In Wales it occurs on common ragwort in five sites in Carmarthenshire, two in Merionethshire (one dating back to 1951), and three in Anglesey.

There is an additional record from *Senecio* sp. in Monmouthshire. On sand sedge it is recorded from North Dock, Carmarthenshire and from Rhosneigr, Aberffraw and Newborough on Anglesey. With so few sites this rust is considered to be **Vulnerable D**.

### Puccinia elymi Westend

Uredinia and telia form on the leaves of lyme-grass *Leymus arenarius* and marram *Ammophila arenaria*. Marram is frequent on sand dunes around the Welsh coast but infections are only visible once the leaves have been unrolled. This makes searches somewhat slow and tedious and the rust might have been much overlooked. Yet the FRDBI only has three records on marram, all from over 50 years ago (including one from Pwllheli, Caernarvonshire). The persistent searches of Arthur Chater, Ian Morgan, Dic Davies and the authors in Wales have recently revealed populations on this host in Glamorgan, Pembrokeshire, Cardiganshire and Caernarvonshire. It is also known from two sites in Carmarthenshire and one in each of Caernarvonshire, Flintshire and Anglesey on lyme-grass. Elsewhere in Britain the FRDBI notes two records in recent years from northern Scotland. Given the paucity of records from Britain as a whole the Welsh populations must take on added significance and are considered to be **Vulnerable D2**.

### Puccinia eriophorae *Thüm*

Aecia are formed on the leaves of golden rod *Solidago virgaurea* (main image below), uredinia and telia on deergrass *Trichophorum cespitosum* agg. (see insert below). Until 2010 it was known from only a handful of sites in Scotland. Then in that year Debbie Evans found populations on both golden rod and deergrass on Moel yr Ogof, Beddgelert and in the

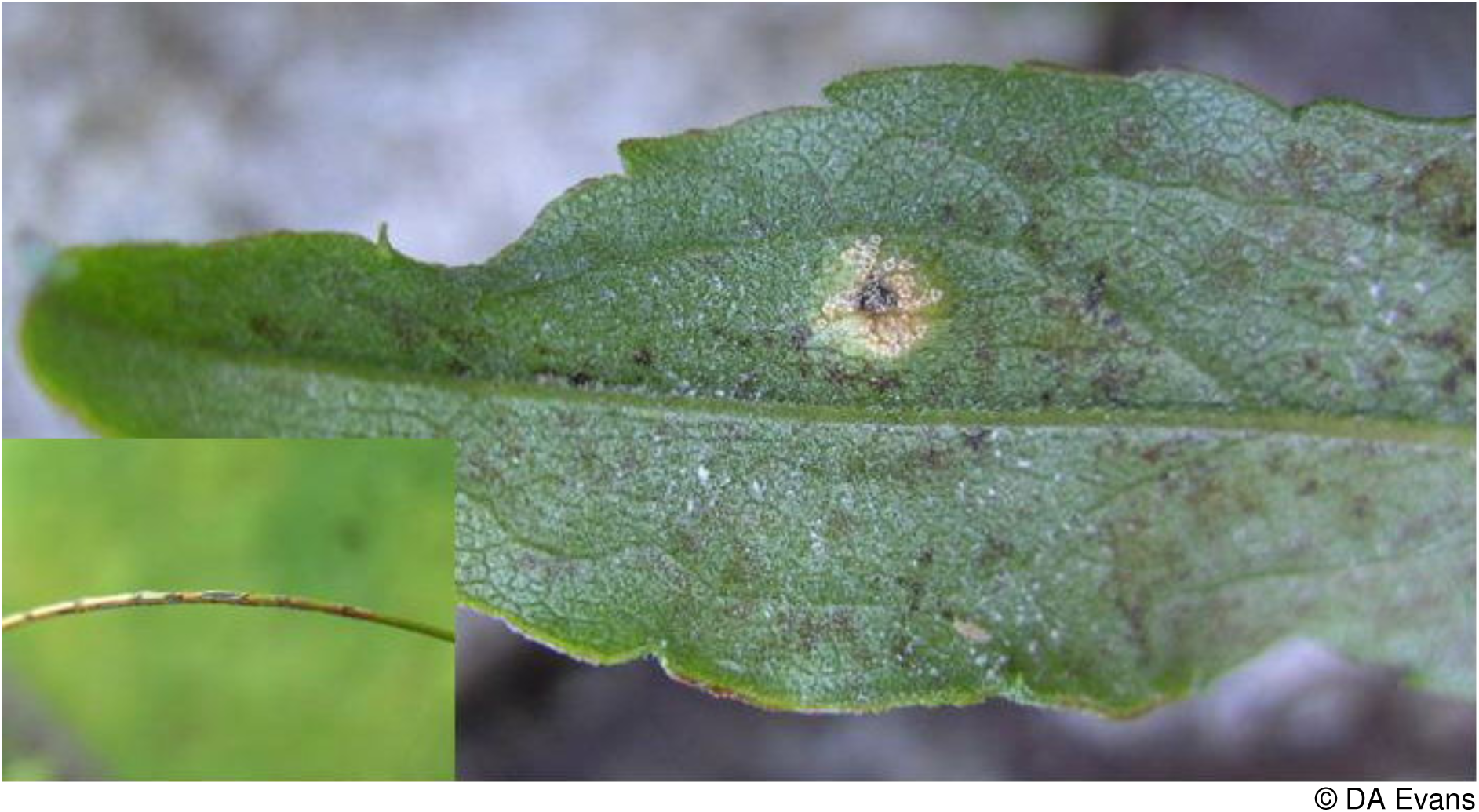

Ogwen Valley, Caernarvonshire. Evans *et al*. (2006) consider this species to be Vulnerable D in Britain. The host species occur together rather uncommonly in Wales, deergrass being favoured by heavy grazing and burning and goldenrod favoured by only light grazing.

Braithwaite *et al*. (2006) note a decline in the extent of goldenrod. Even if this rust is overlooked it is unlikely to be anything but rare and is isolated from the nearest Scottish populations. It is considered to be **Endangered D** in Wales since it is known from two populations. If on further survey the total population numbers fewer than 50 individuals it might be considered to be Critically Endangered.

### Puccinia fergussonii Berk. & Broome

A rarely-encountered rust on marsh violet *Viola palustris,* with ten recent records from Scotland (mostly in the north) and five records in northern England on the FRDBI. There are fewer than 20 recent records from 12 hectads in a wide scatter of localities in Wales.

Braithwaite *et al*. (2006) note a marked decline in the distribution of marsh violet and the demise of fritillary butterflies dependant on marsh violet has highlighted the rather narrow set of conditions required by this plant. The spread of conifer forests and intensive sheep grazing must also have reduced the size of its population. Given the relatively small population of the rust on fewer than 250 marsh violet colonies in Wales and threats to its host, this rust is considered to be **Endangered D**.

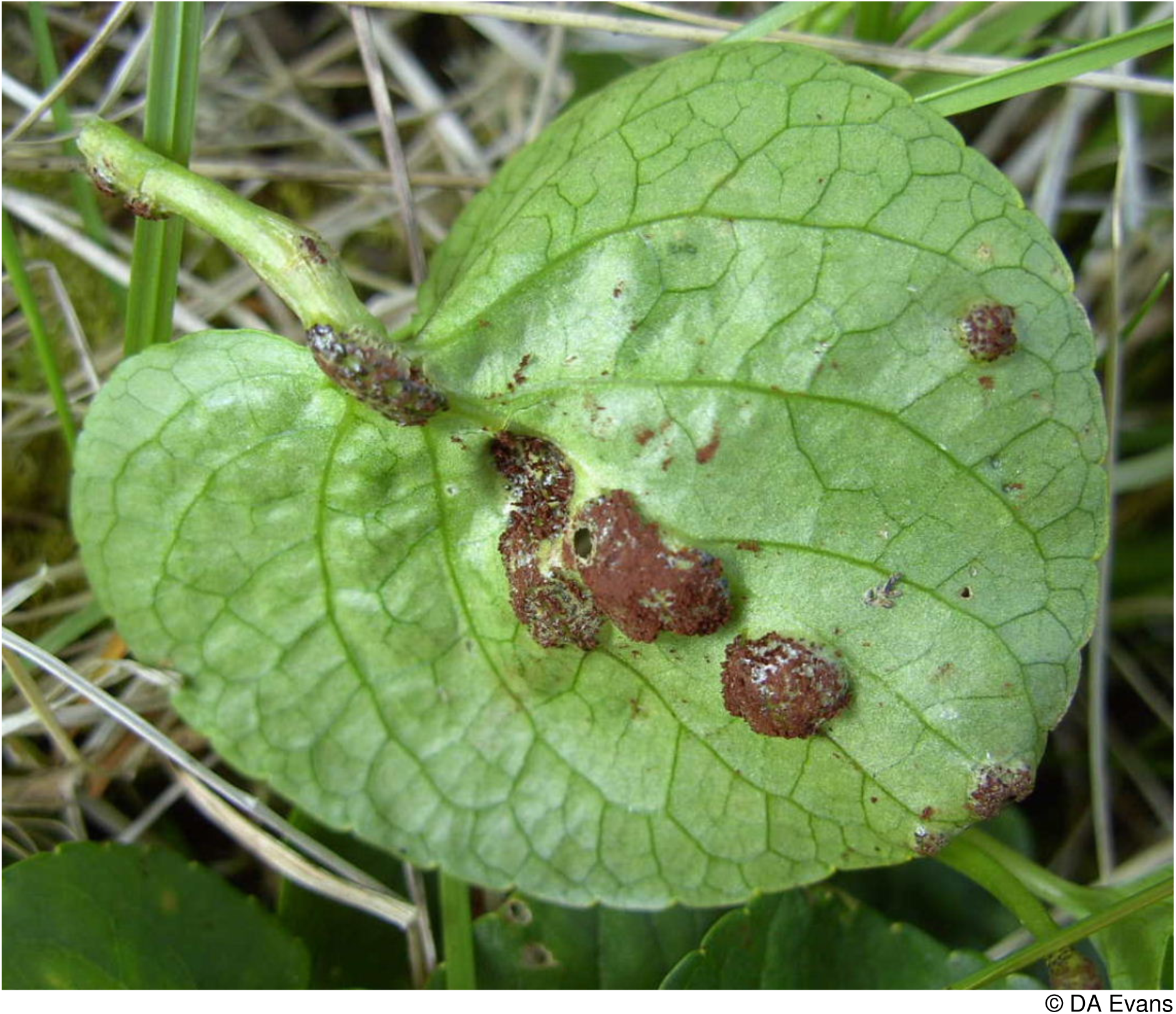

### Puccina galii-cruciatae ***Duby***

*P. galii-cruciatae* is the least common of the three rusts that occur on crosswort *Cruciata laevipes* with but single records from Glamorgan, Breconshire and Carmarthenshire. Elsewhere in Britain the FRDBI notes 10 recent records. Whilst mostly from southern Britain there is a collection of records from Shropshire close to the Welsh border. Older records were mainly from the north of England. In view of the small population size and continued threat to its host from agricultural intensification and particularly eutrophication of roadside verges (Preston *et al*. 2002,) this rust is placed in the **Endangered B2, D** category.

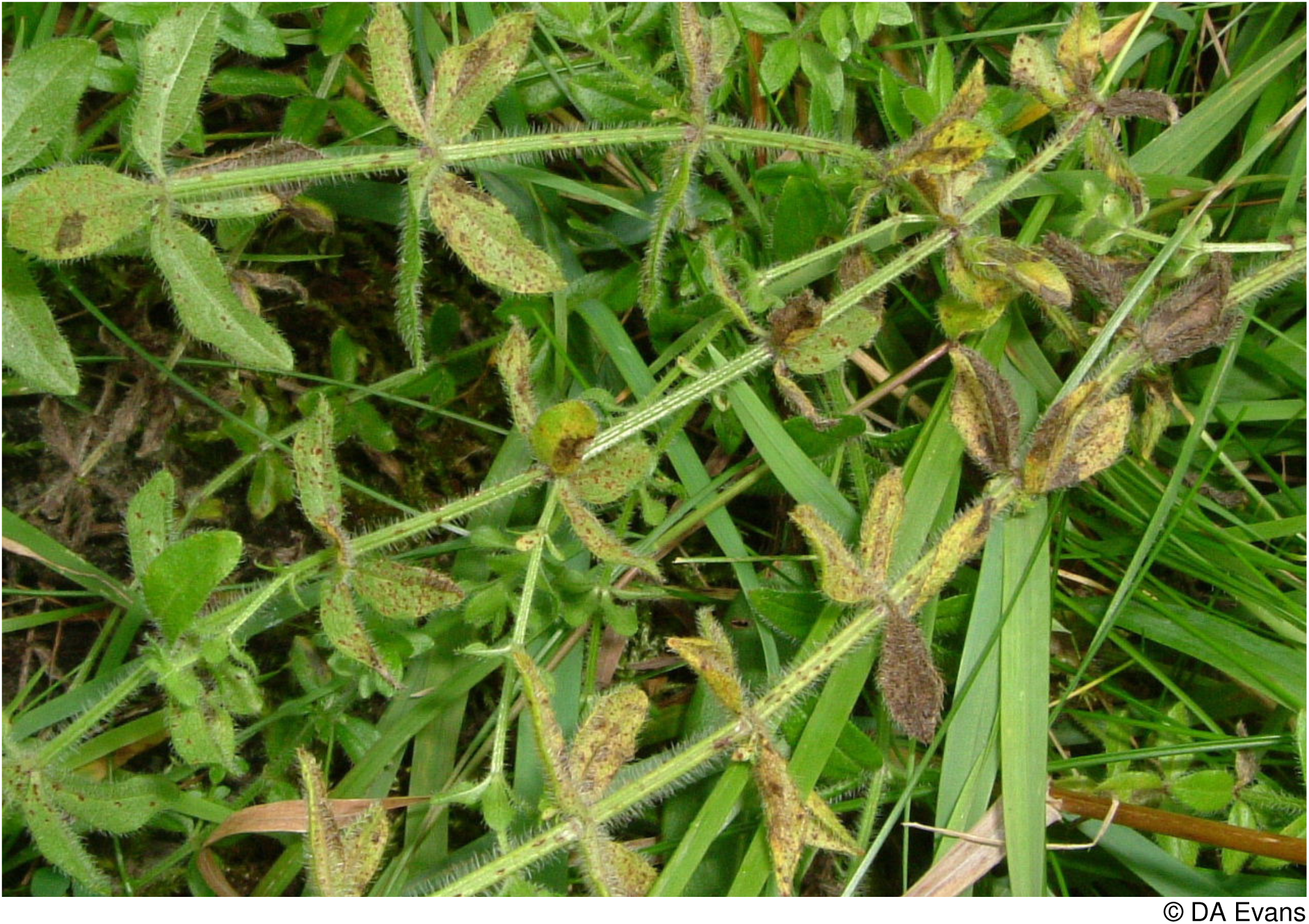
*Puccina galii-cruciatae* on *Cruciata laevipes* (see p38)

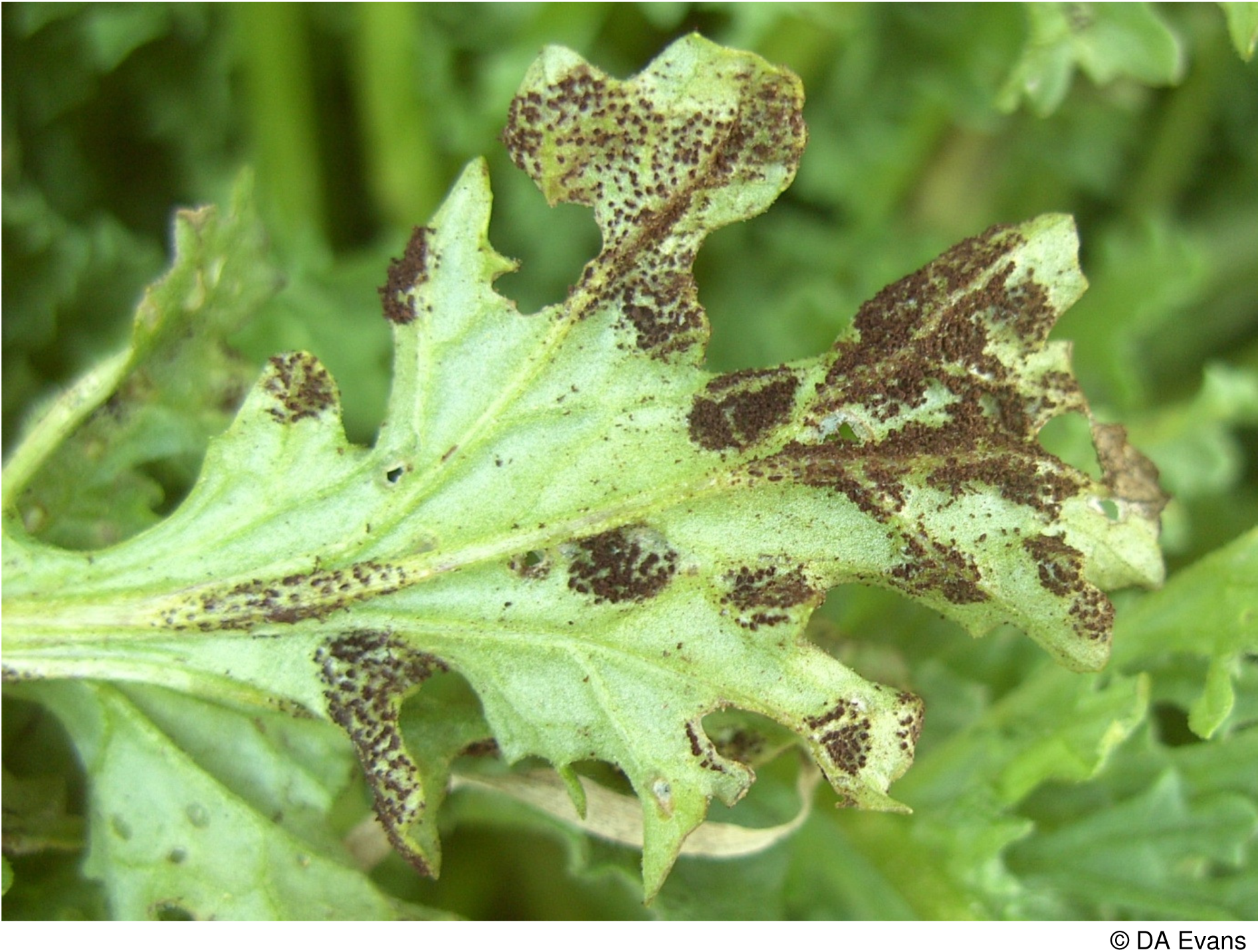
*Puccinia glomerata* on common ragwort (see p40)

### Puccinia glomerata Grev

Occurring on both common ragwort *Senecio jacobaea* (see image overleaf) and marsh ragwort *S. aquatica,* there are surprisingly few records from Wales from such widespread species. On marsh ragwort there is a single record from Montgomeryshire and on common ragwort six records from Carmarthenshire, three in close proximity to one another from Ynyslas in Cardiganshire, three from Caernarvonshire, three from Anglesey and one each from Montgomeryshire, Denbighshire and Flintshire. There are ten recent records from common ragwort in England and nine from Scotland on the FRDBI. On marsh ragwort there are four from England and eight from Scotland. Given the small number of known sites, the loss of wetland due to agricultural drainage and the consequent loss of habitat for marsh ragwort and the general persecution of common ragwort due to its toxicity to grazing stock, this rust is placed in the **Near Threatened** category.

### Puccinia graminis Pers

Earlier work splitting this rust into two subspecies has not been confirmed by DNA analysis. There is, however, some evidence of host specificity such as on sweet vernal-grass (Wilson and Henderson 1966). In view of its very wide host range and potential impact on commercial forage grasses further work should be stimulated and it is placed in the **Data Deficient** category.

### Puccinia heraclii Grev

Infecting hogweed *Heracleum sphondylium*, there are no recent records from Wales. The FRDBI lists records from Lampeter in Cardiganshire in 1841, Llandudno in Caernarvonshire in 1865, Corwen in Merionethshire in 1865 and 1872 and from Bala in 1866, Harlech in 1900 and Nannau in 1909 in the same vice-county. It should be sought in Wales since there were records in both 2003 and 2012 from Richard’s Castle in Herefordshire not far from the Welsh border. For the moment it is considered **Regionally Extinct**.

### Puccinia hydrocotyles (Mont.) Cooke

Considered by Evans *et al*. (2006) to be Critically Endangered B in Britain, this rust of marsh pennywort *Hydrocotyle vulgaris* has subsequently been found in at least five new localities and Vulnerable may now be a more up to date category. In Wales it is very uncommon with one record from Carmarthenshire, three from Cardiganshire and four from Newborough Forest in Anglesey. It occurs in dune slacks and fen meadows in generally high nature conservation value sites. Infections are relatively easy to spot and the paucity of records probably genuinely reflects its rarity. The widespread drainage of wetlands in the last 60 years has much diminished its potential number of hosts (Braithwaite *et al*. 2006). Given the relatively small population which must number fewer than 250 infected marsh pennywort plants it is considered to be **Endangered D**.

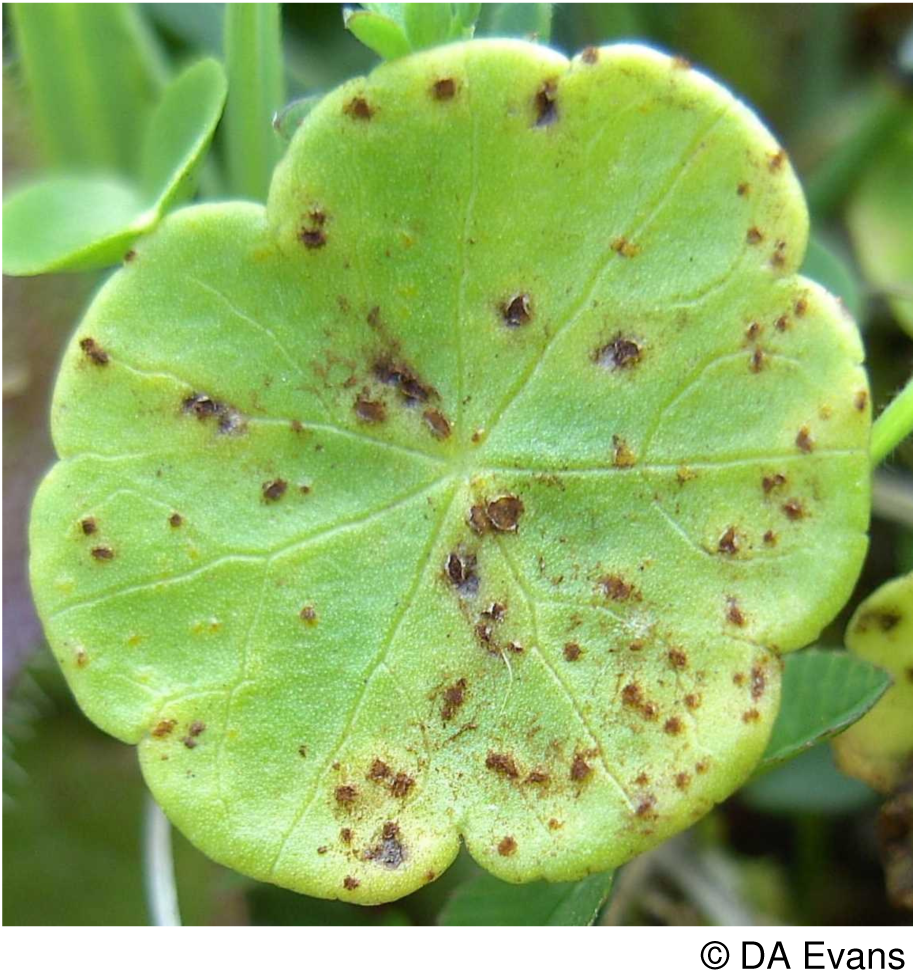

### Puccinia hysterium (F. Strauss) Röhl

The orange aecia can be present on goat’s-beard *Tragopogon pratensis* leaves and stems in great abundance, causing distortion and rendering infection very obvious. This rust is widespread in central and southern England with over 45 recent (post 1964) records in the FRDBI. It is rare in Wales with but single recent records from Glamorgan and Pembrokeshire. Vize records it from the Parish of Forden, Montgomeryshire pre 1882 (Vize 1882). The host plant is never very frequent in Wales except in the lowlands and the Welsh population probably represents an outlier of a population centred on England. As such its threat status has been downgraded by a criterion in line with IUCN advice and it is considered **Vulnerable D1**.

### Puccinia luzulae ***Lib*.**

This rust of hairy wood-rush *Luzula pilosa* in Wales is reported from only a handful of sites. The FRDBI lists recent records from the Hengwrt Estate near Dolgellau, Plas Tan-y-bwlch near Maentwrog and Morfa Harlech near Harlech, all in Merionethshire. There are old records from Cardiganshire (site unspecified), Forden, Montgomeryshire and Swallow Falls, Caernarvonshire. Aron (2005) in the latter vice-county lists a record from Halfway Woods in 1973 and in 2013 Nigel Stringer recorded this rust from near Llandyfaelog, Carmarthenshire. Searches of many populations of hairy wood-rush have failed to locate this rust elsewhere. It is placed in the **Vulnerable D** category.

### Puccinia maculosa Schwein

The many small, neat brown spots of the uredinia and telia are easy to locate on the leaves of wall lettuce *Mycelis muralis.* There have, however, been few records from Wales and this species is distinctly uncommon with probably fewer than 250 individual host plants colonised. Three modern records have been made from Breconshire, two from Radnorshire, three from Carmarthenshire, one from Merioneth and two from Denbighshire, one of which was made pre 1964. There are also two pre 1964 records from Caernarvonshire. Since there are no recent records from anywhere on the Welsh Marches in the FRDBI the Welsh population is considered to be isolated and a conservation evaluation of **Endangered D** to be appropriate.

### Puccinia major Dietel

The last Welsh record was on marsh hawk’s-beard *Crepis paludosa* at Torrent Walk in Merioneth in 1887. It had previously been recorded from Aberdyfi in the same vice-county in 1874. The nearest recent British records are from West Yorkshire. The host persists in a number of sites in ravines and wet woodlands in Wales. For the time being this rust must be considered to be **Regionally Extinct**.

### Puccinia nemoralis ***Juel***

Alternating between common cow-wheat *Melampyrum pratense* and purple moor-grass *Molinia caerulea,* this rust is only known for certain in Britain from Wales. It is reported on both hosts from Coed Cymerau National Nature Reserve and Traeth Glaslyn, Merioneth (Aron 2005), near Bethesda (Debbie Evans pers. comm.) and near Garth-Marthin, Caernarvonshire (FRDBI).

There is also a record on cow-wheat from near Dolgellau in 1886. Evans *et al*. (2006) have placed it in the Vulnerable D2 category on account of its limited population size. Braithwaite *et al*. (2006) note a reduction in distribution of common cow-wheat attributed to a general increase in shade and/or eutrophication of woodlands. A cessation of grazing in many North Wales oakwoods has resulted in the spread of holly and ivy leading to increased shade. Following Evans *et al*. (2006) it is categorised as **Vulnerable D2** in Wales.

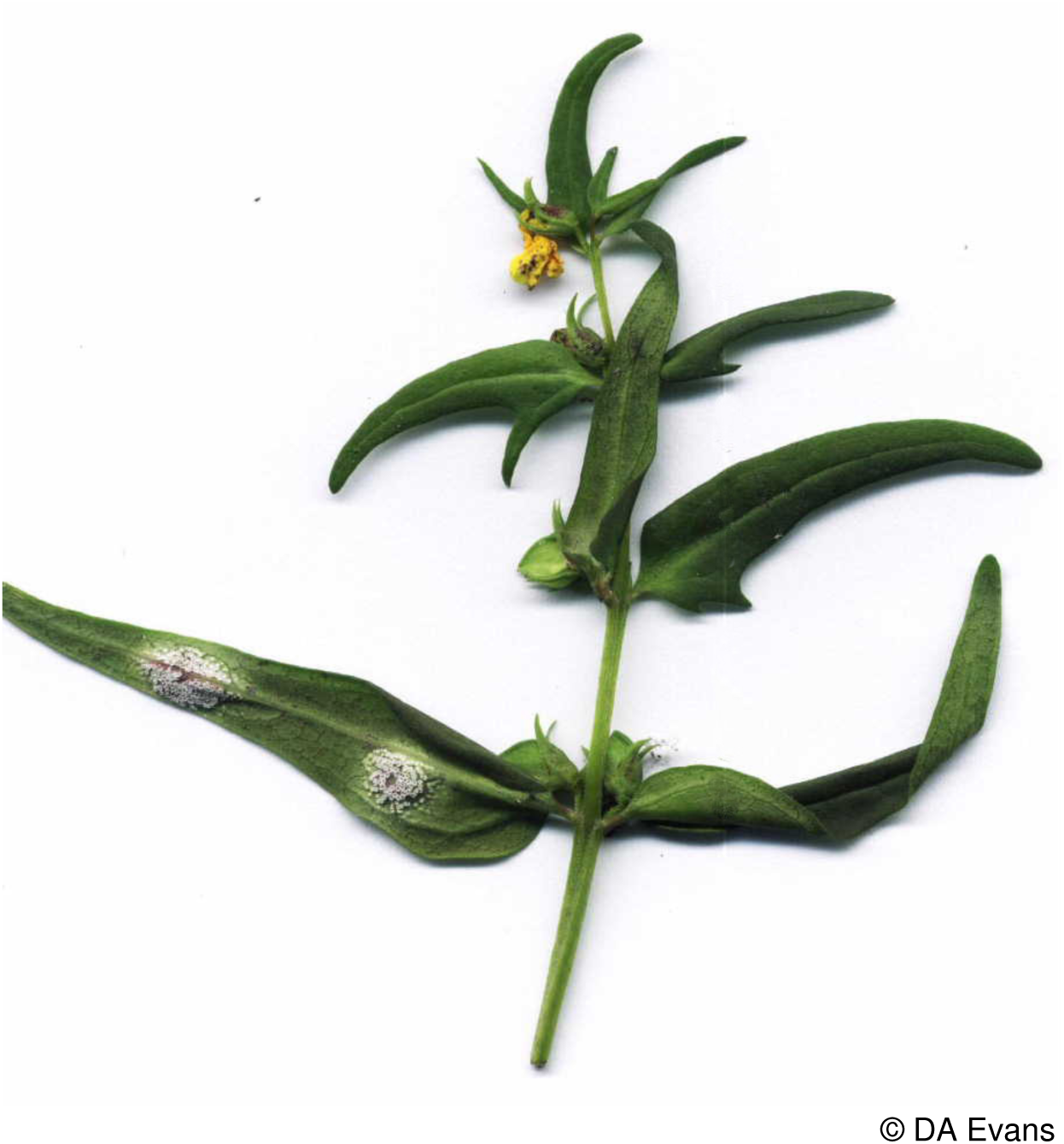

### ***Puccinia nitida*** (F. Strauss) Röhl

Occurring on fool’s parsley *Aethusa cynapium* this rust has been found recently in single localities in Monmouthshire, Radnorshire, Carmarthenshire, Cardiganshire, Montgomeryshire and in 1945 from Caernarvonshire. Weeds in general have declined in the last 50 years due to a reduction in the area of arable crops and more recently by the use of very effective weed killers (Braithwaite *et al*. 2006). Its status in Wales might therefore be considered to be Endangered D1. There is, however a widespread population in England. As such its threat status has been downgraded by a threat category in line with IUCN advice and it is considered to be **Vulnerable D2**.

### Puccinia opizii Bubák

The spermogonia and aecia on garden lettuce *Lactuca sativa* and great lettuce *L. virosa* have not been reported in Wales. Uredinia and/or telia have, however, been reported from one site in each of Breconshire and Radnorshire and three sites in Cardiganshire on prickly-sedge *Carex muricata* ssp. *pairae* (= ssp. *lamprocarpa*); one site in Glamorgan on spiked sedge *C. spicata* and four sites in Carmarthenshire and one site in Cardiganshire on greater tussock-sedge *C. paniculata*. All other British records in FRDBI are from eastern England and distant from the Welsh populations. In view of the recent drainage of many bogs and the loss of dry lowland grassland (Stevens *et al*. 2010) reducing the abundance of its hosts, this rust is considered to be **Near Threatened**.

### Puccinia oxyriae Fuckel

Found only on mountain sorrel *Oxyria digyna,* Evans *et al*. (2006) considered this rust extinct in Britain. In 2007 it was, however, found on Ben Lawers in Scotland indicating that it should now be considered to be Critically Endangered. In Wales it was found in the Devil’s Kitchen, Cwm Idwal, Caernarvonshire by E.A. Ellis in 1941 and by Debbie Evans in Cwm Glas Bach, Snowdon in 2007. Only a single infected leaf was detected in this latter site. Given that it is known recently only from a single plant in a single locality on a host threatened by grazing and climatic warming, this rust is considered to be **Critically Endangered D1 & 2** in Wales.

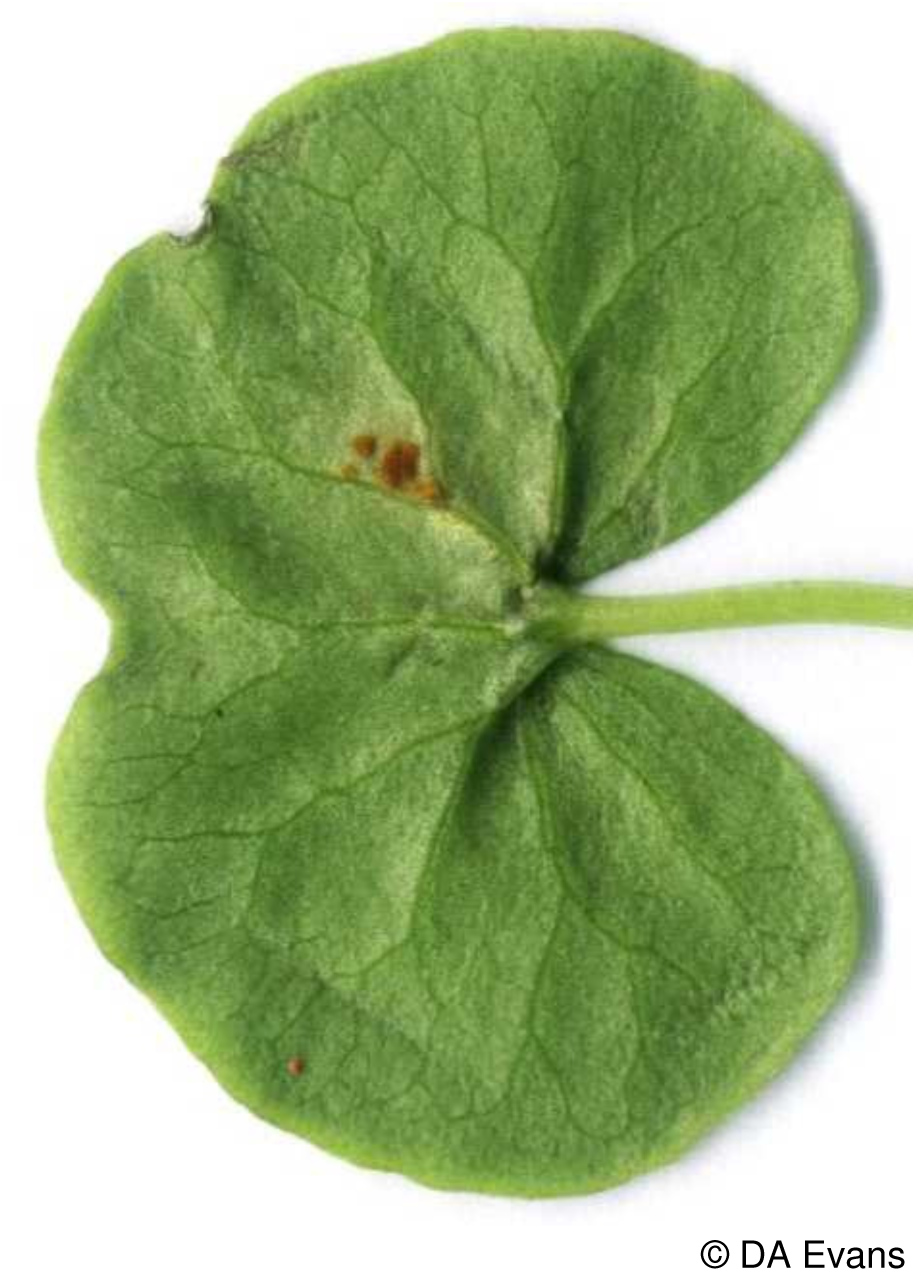

### Puccinia pimpinellae (F. Strauss) Link

Noted in Wales only on burnet-saxifrage *Pimpinella saxifraga* and then very rarely.

There are four recent records *viz*. one each from Breconshire and Caernarvonshire and two from Radnorshire. In Breconshire it occurred in short turf of the old castle mound, Builth Wells; the Caernarvonshire site was in very exposed coastal turf on the Little Orme, whilst one of the Radnorshire localities was from a roadside verge and the other a disused railway cutting. There are in addition two old records, one from Lampeter, Cardiganshire in the 1800s and the other from Sketty Park, Glamorgan in 1915. There are over 30 recent records from England, most being from greater burnet-saxifrage *P. major* in Yorkshire with very few records from a scatter of localities on burnet-saxifrage (FRDBI). In view of the limited number of sites in Wales, the probability that fewer than 50 plants are infected in total and the threat to the short turf/dry grassland of its host (Stevens *et al*. 2010) this rust is considered to be **Critically Endangered D2**.

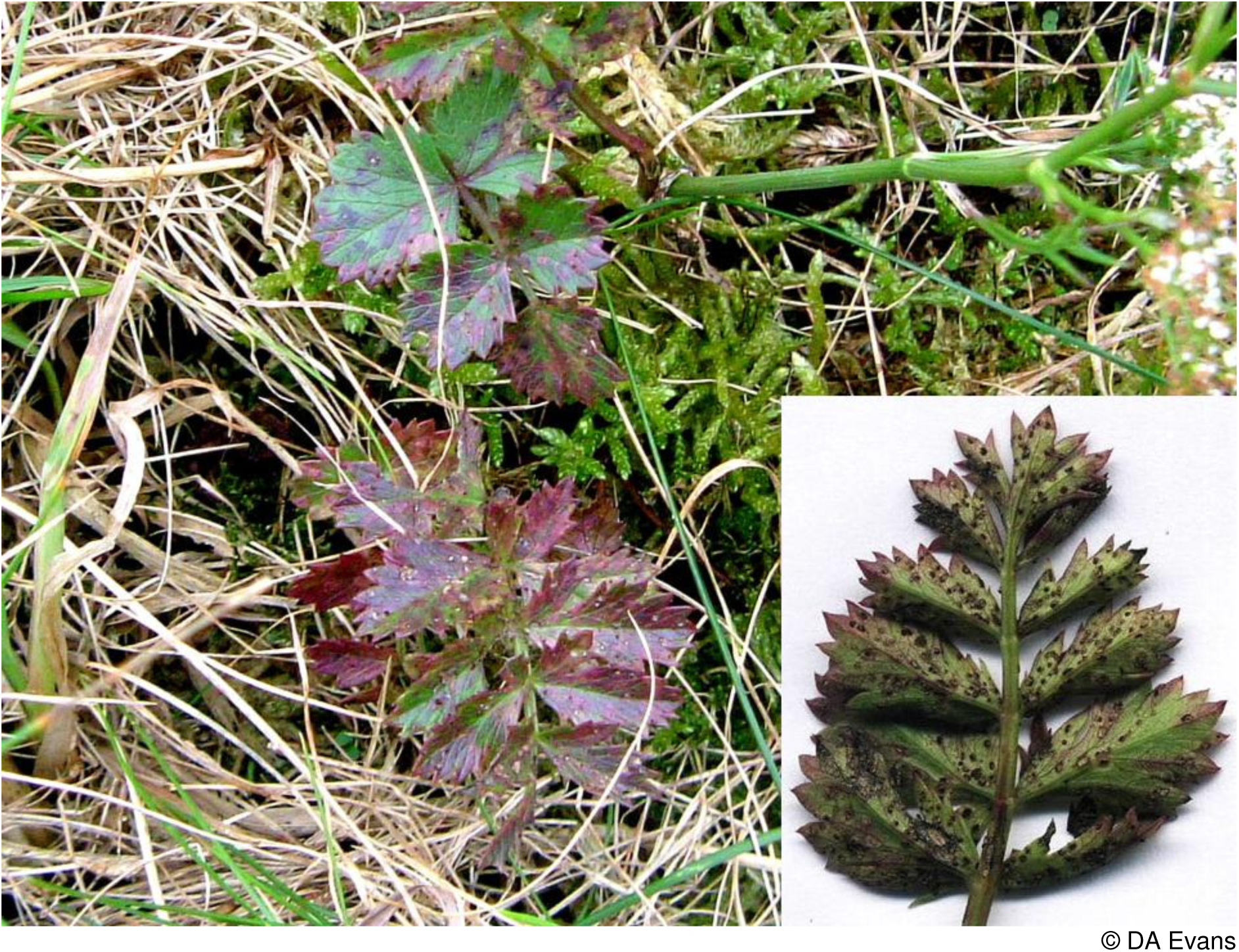

### Puccinia polygoni-amphibii var. convolvuli Arthur

The only localised records from Wales are on black-bindweed *Fallopia convolvulus*. It is recorded from Bridgend, Glamorgan in 1957 and more recently from near Llysdinam, Breconshire, Bigyn allotments in Carmarthenshire and Henfaes University Farm, Aber, Caernarvonshire. The FRDBI lists only 22 recent records on black-bindweed and cut-leaved crane’s-bill *Geranium dissectum* from all of England. The recent widespread use of pre-emergence weed-killers on, for example root crops, has greatly diminished the weed flora of farmlands (Braithwaite *et al*. 2006). This species is considered to be **Vulnerable D**.

*Puccinia polygoni-amphibii* var. *convolvuli* on black-bindweed

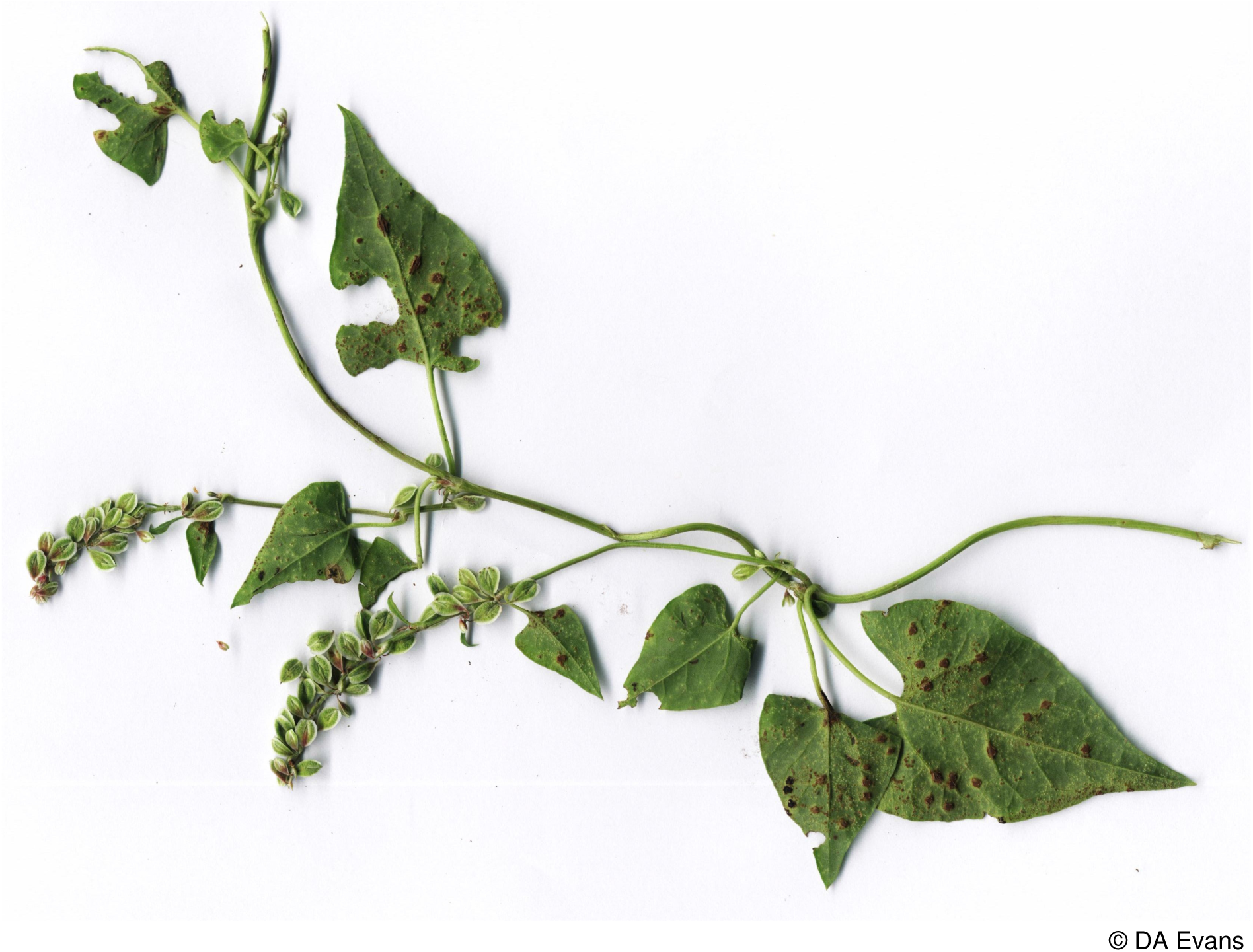

### Puccinia primulae (DC.) Duby

This conspicuous rust of primrose *Primula vulgaris* that presents no difficulty in identification is unaccountably rare in Wales. The FRDBI, whilst having nearly 350 records from Britain and Ireland has only two pre 1990 records from Wales viz. a 1913 record from Arthog Bog, Merioneth and a 1987 record from West Williamston in Pembrokeshire and the database of the ABFG one record. To these can be added two recent records from Pembrokeshire, six from Carmarthenshire, two from Cardiganshire, one from Caernarvonshire and six sites on

*Puccinia primulae* on cowslip

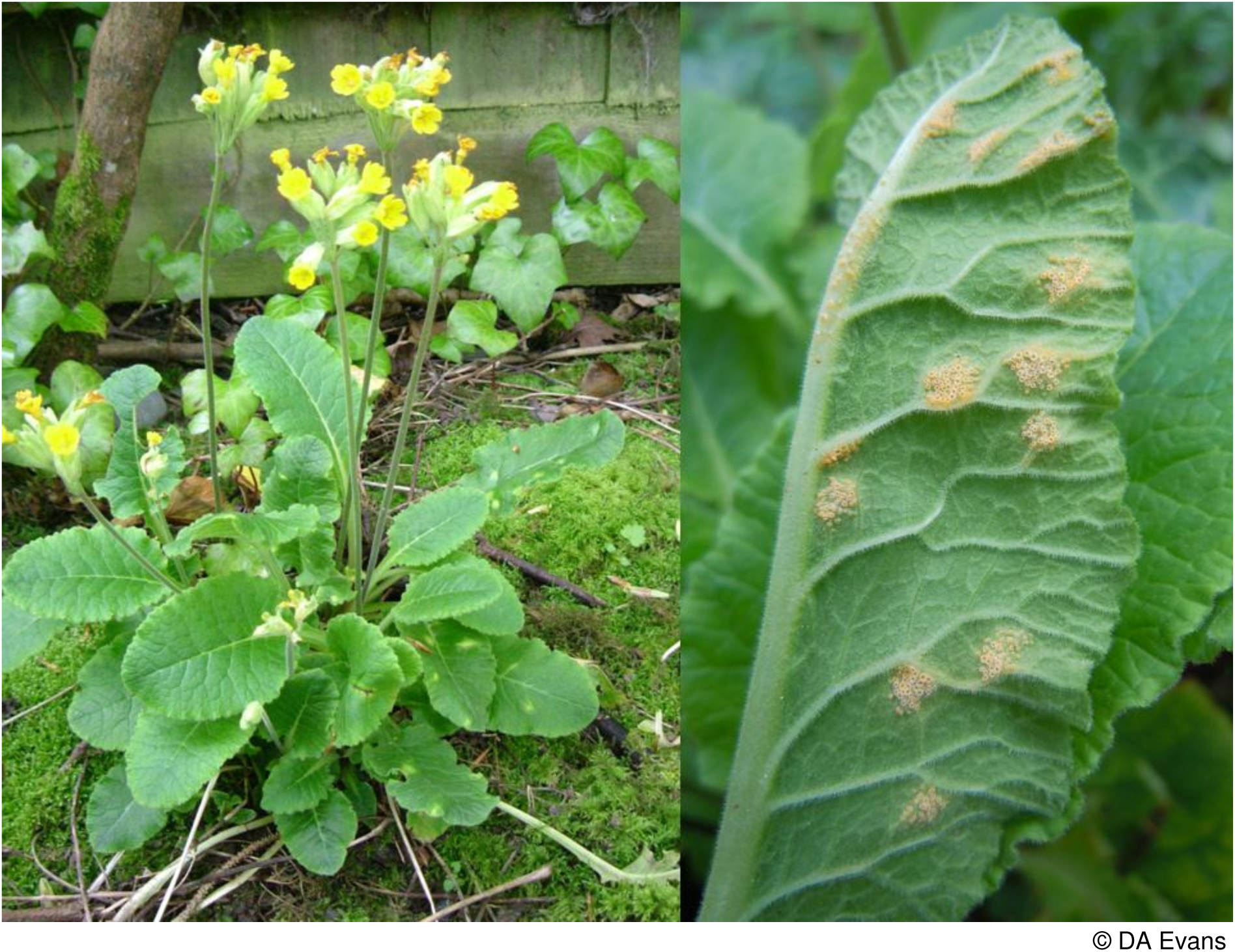

Anglesey. This latter vice-county has a concentration of records from around Holyhead including a record on a coloured primrose in an abandoned garden and a very rare record on cowslip *P. veris* in Penrhos Park (see images above). Notably in at least its North Wales sites it appears regularly year after year in the same places. Given the threats to its host plants from agricultural intensification, the eutrophication of hedgerows, heavy grazing pressure and plantation forestry this rust has to be considered to be **Near Threatened**.

### Puccinia pygmaea var. pygmaea Erikss

This variety is confined to wood small-reed *Calamagrostis epigejos*, a host species almost exclusively coastal in Wales. The FRDBI contains 18 records from Britain, almost all from the North of England and East Anglia. The single known recent Welsh population at Mwnt in Cardiganshire is very isolated. There is a 1928 record presumably on this host from Red Wharf Bay, Anglesey. It is considered to be **Critically Endangered D1** in Wales.

### Puccinia saxifragae Schltdl

Recorded in Wales from both starry saxifrage *Saxifraga stellaris* and meadow saxifrage *S. granulata*, but with very few records. On the former species it has been recorded less than a dozen times viz. recently from a single site in Merioneth and from 5 sites in Snowdonia, Caernarvonshire. Historically it has been recorded twice from Cadair Idris, Merioneth (in 1923 and 1948) and from Llyn Idwal and Y Gribin, Caernarvonshire in 1941. J.H. Salter found it above Llyn Llygad Rheidol, Cardiganshire in 1922. There are no recent records on the FRDBI from England. There are even fewer records from meadow saxifrage with just four ever from Britain viz. Warwickshire in the 1900’s, East Gloucestershire in 1918, Dovedale in Derbyshire in 1924 and East Norfolk in 1987. To these four can be added a single Welsh record from a woodland edge in the Edw Valley, Radnorshire where it occurred over a few square metres. This site has no formal protection. Starry saxifrage is at the southern-most edge of its British range in Wales and is a species at threat from a warming climate. Braithwaite *et al*. (2006) note a significant decrease in meadow saxifrage between 1987 and 2004. In view of the small size of the known extant population in Wales it is considered to be **Endangered D**.

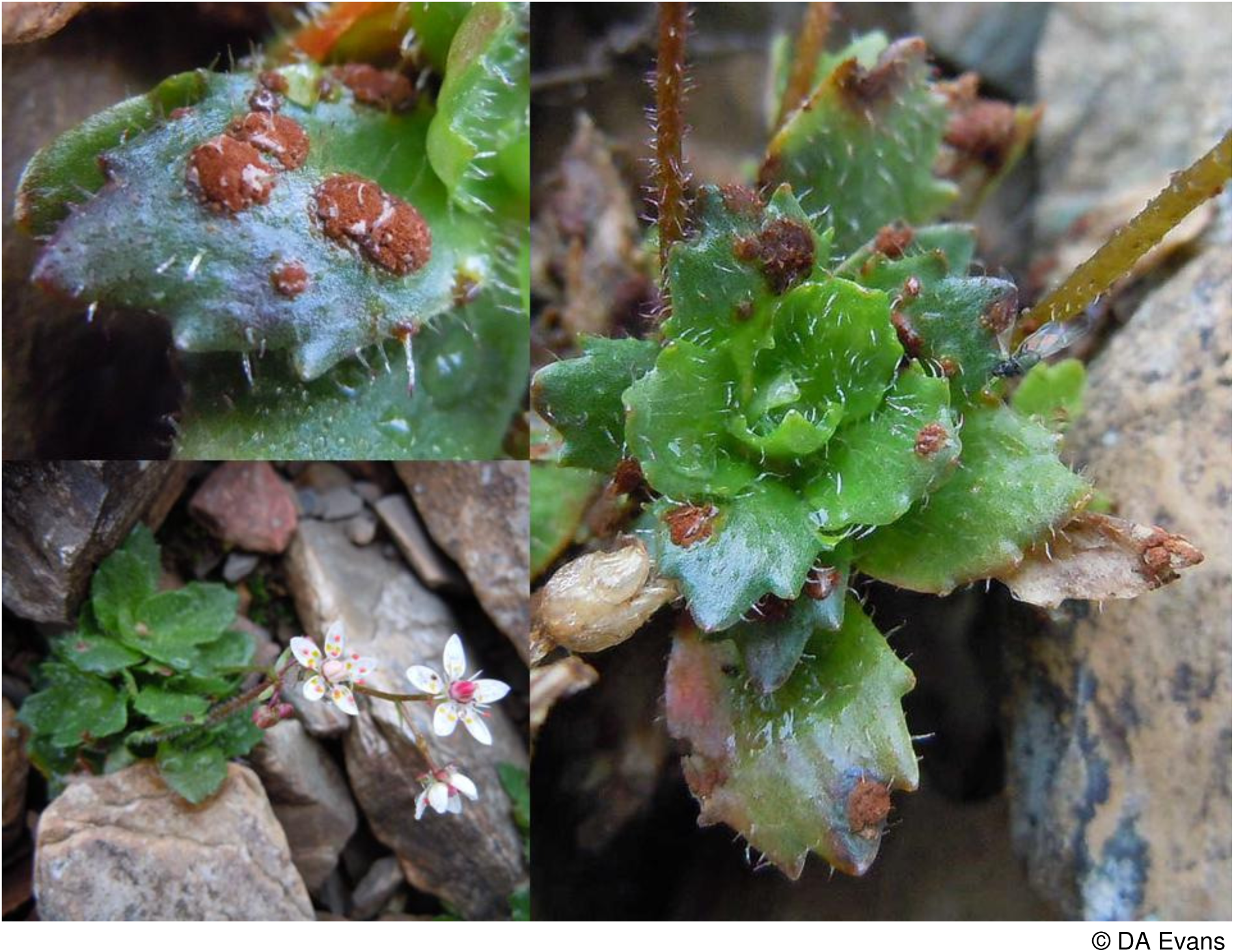
*Puccinia saxifragae* on starry saxifrage

### Puccinia schroeteriana ***Kleb*.**

This rust was discovered new to Britain by Debbie Evans in 2008. Uredinia and telia occurred on tawny sedge *Carex hostiana* and aecia on saw-wort *Serratula tinctoria* at Cors Bodeilio, Anglesey. It has subsequently been found on both hosts at Cors Goch in the same vice-county. The nearest known sites are in continental Europe. It is placed in the **Critically Endangered D2** category.

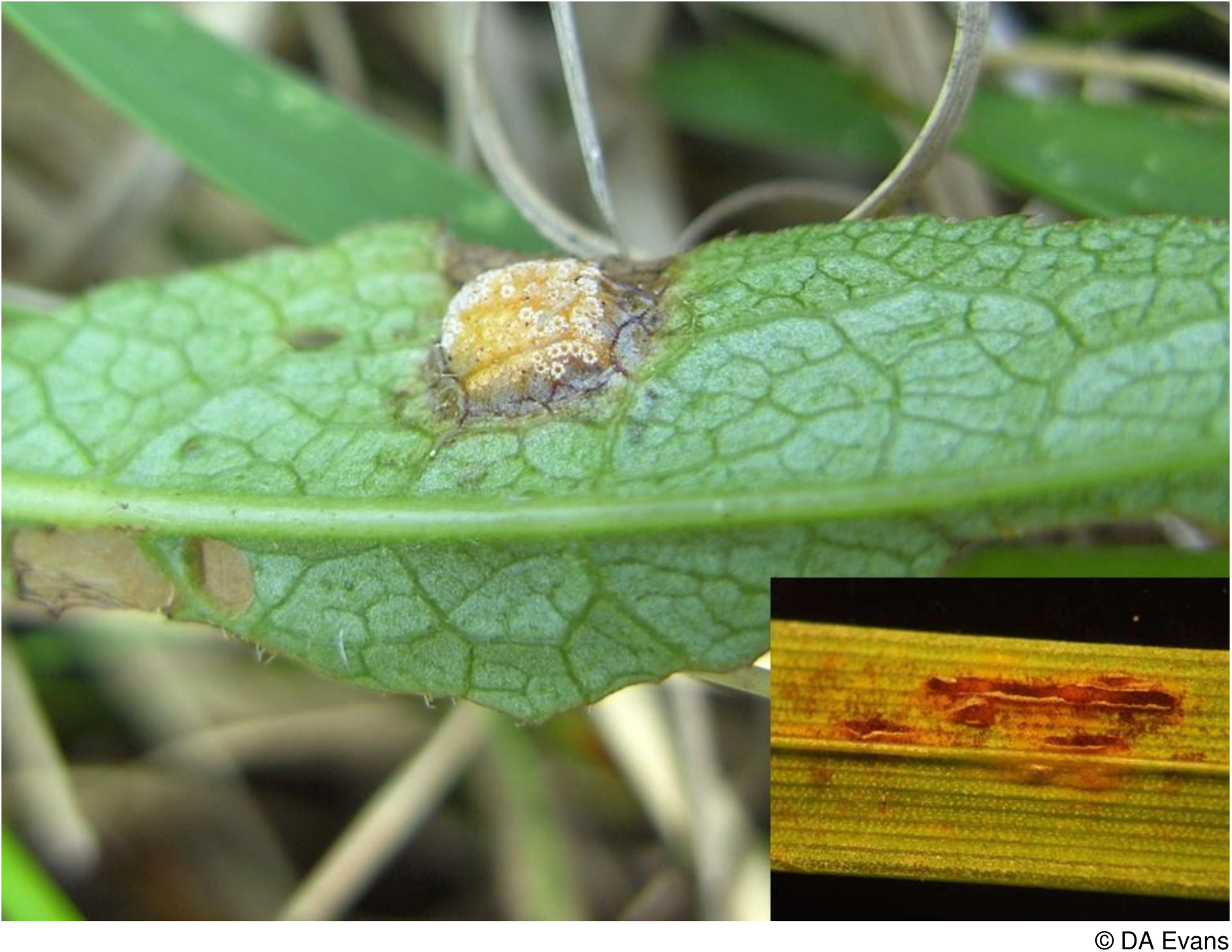
*Puccinia schroeteriana* on saw-wort (main image) and tawny sedge (insert)

### Puccinia scirpi ***DC*.**

This rust fungus alternates between fringed water-lily *Nymphoides peltata* and common club-rush *Schoenoplectus lacustris.* Wilson and Henderson (1966) note a record from Llangorse Lake in Breconshire, though give no date or details of the recorder. The FRDBI lists no records from Wales, the nearest being on common club-rush in Moccas Park, Bredwardine, Herefordshire in 1951. There is but a single recent record from Britain (Northamptonshire in 2010) and Evans *et al*. (2006) consider this species to be Critically Endangered. Legon *et al*. (2006) cast doubt on the Breconshire record but given that both hosts grow in Llangorse Lake and there is the record from nearby Herefordshire, it is entirely plausible that this rust may still occur in Wales. Pending a survey it is placed in the **Data Deficient** category.

### Puccinia scorzonerae (Schumach.) Jacky

This is a national Biodiversity Action Plan species and is listed as Critically Endangered D by Evans *et al*. (2006). It appears to be frequent on the relatively large populations of viper’s-grass *Scorzonera humilis* on a single site in Glamorgan though has not been found recently at the two other known sites for the host in Dorset. Cheffings *et al*. (2005) list the host plant as a British national RDB Vulnerable species as it is in Wales (Dines 2008). This rust is entirely dependent on this single host species and the Welsh population is at the very edge of the European range of the rust. One Dorset host population is believed to be at risk from sea level rise and management of the other site may well have been sub-optimal in the recent past. The Glamorgan population is protected to some extent by a SSSI designation. Given that the future of this rust is dependent on the management of a single site it is considered to be **Critically Endangered D1** in Wales as well as in Britain.

### ***Puccinia thymi*** (Fuckel) P.Karst

There have only ever been a handful of records on native species of thyme *Thymus* spp. in Britain. Evans *et al*. (2006) consider it to be Vulnerable D2. In Wales it has been found twice, on both occasions on wild thyme *T. polytrichus*, with one record from Pembrokeshire and one from ORS outcrops in Cwm Sere in the Brecon Beacons, Breconshire. In view of the small population, this rust is considered to be **Endangered D**.

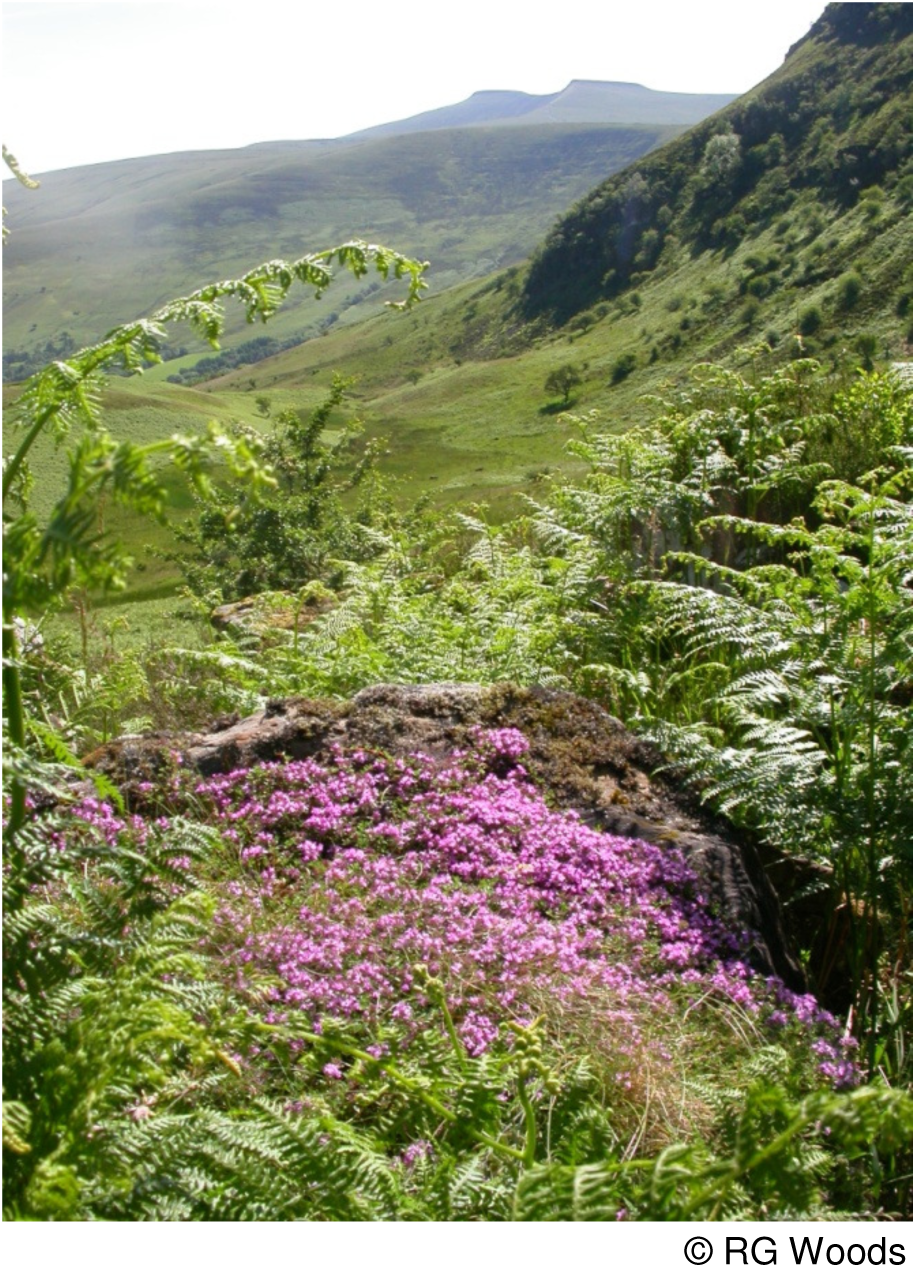

Habitat of *Puccinia thymi* on wild thyme in the Brecon Beacons with Cwm Sere in the distance.

### Puccinia uliginosa ***Juel***

Aron (2005) has but a single record from common sedge *Carex nigra* from Lligwy on Anglesey in 1948 with no indication as to the recorder. This rust alternates with grass-of-parnassus *Parnassia palustris* (see image on facing page). There are eight records on the FRDBI with only two on common sedge from Orkney in the 19^th^ century and the Isle of Ulva, Scotland in 1968. The few records of this rust on grass-of-parnassus are confined to the north of England and Scotland. Anglesey supports the main population of this flower in Wales so the possibility of this rust occurring there cannot be discounted. Given the long time interval since the last record, this rust is considered to be **Regionally Extinct**.

Opposite: (top) *Puccinia uliginosa* on grass-of-parnassus (bottom) *Puccinia urticata* var. *biporula* on pale sedge (see p50).

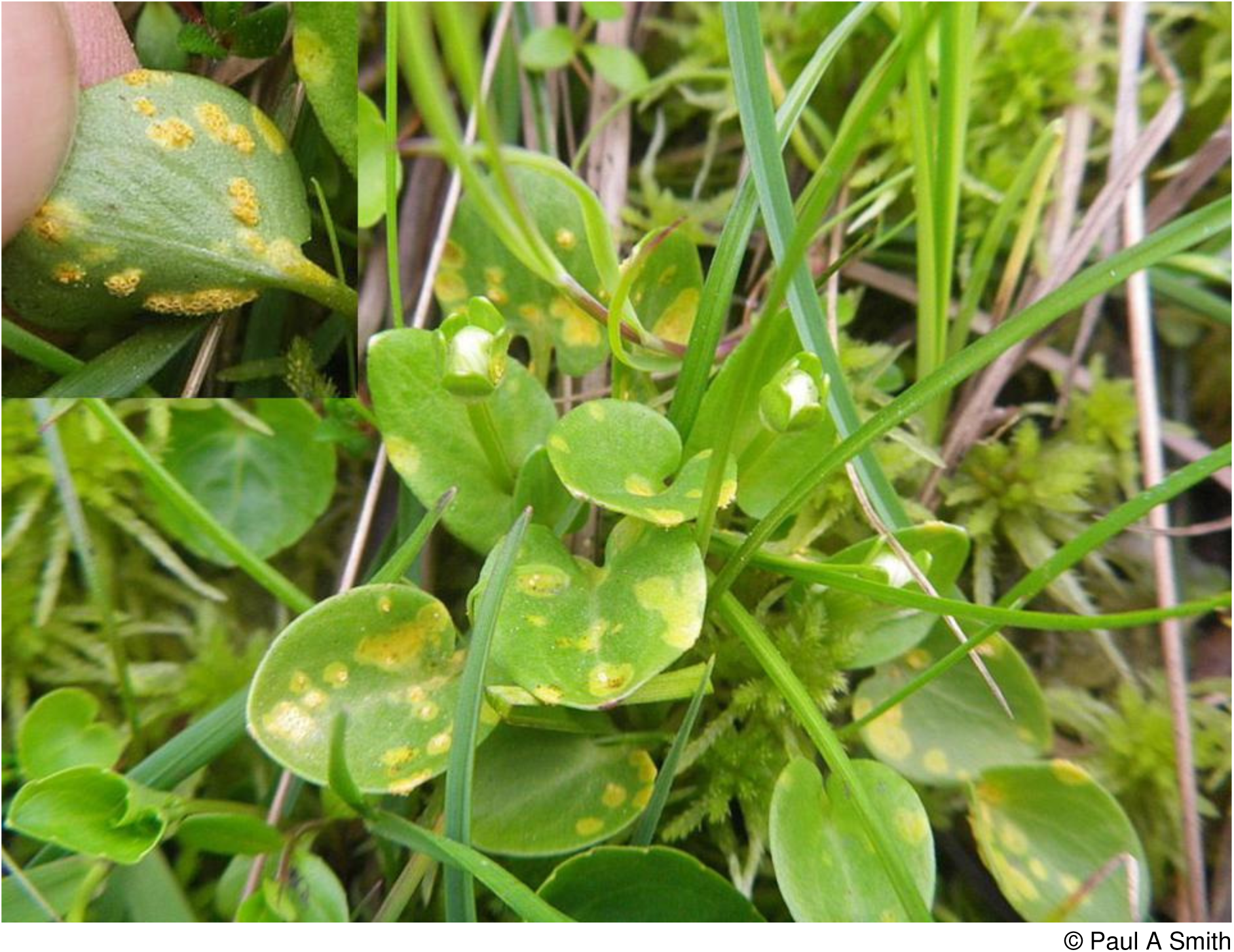

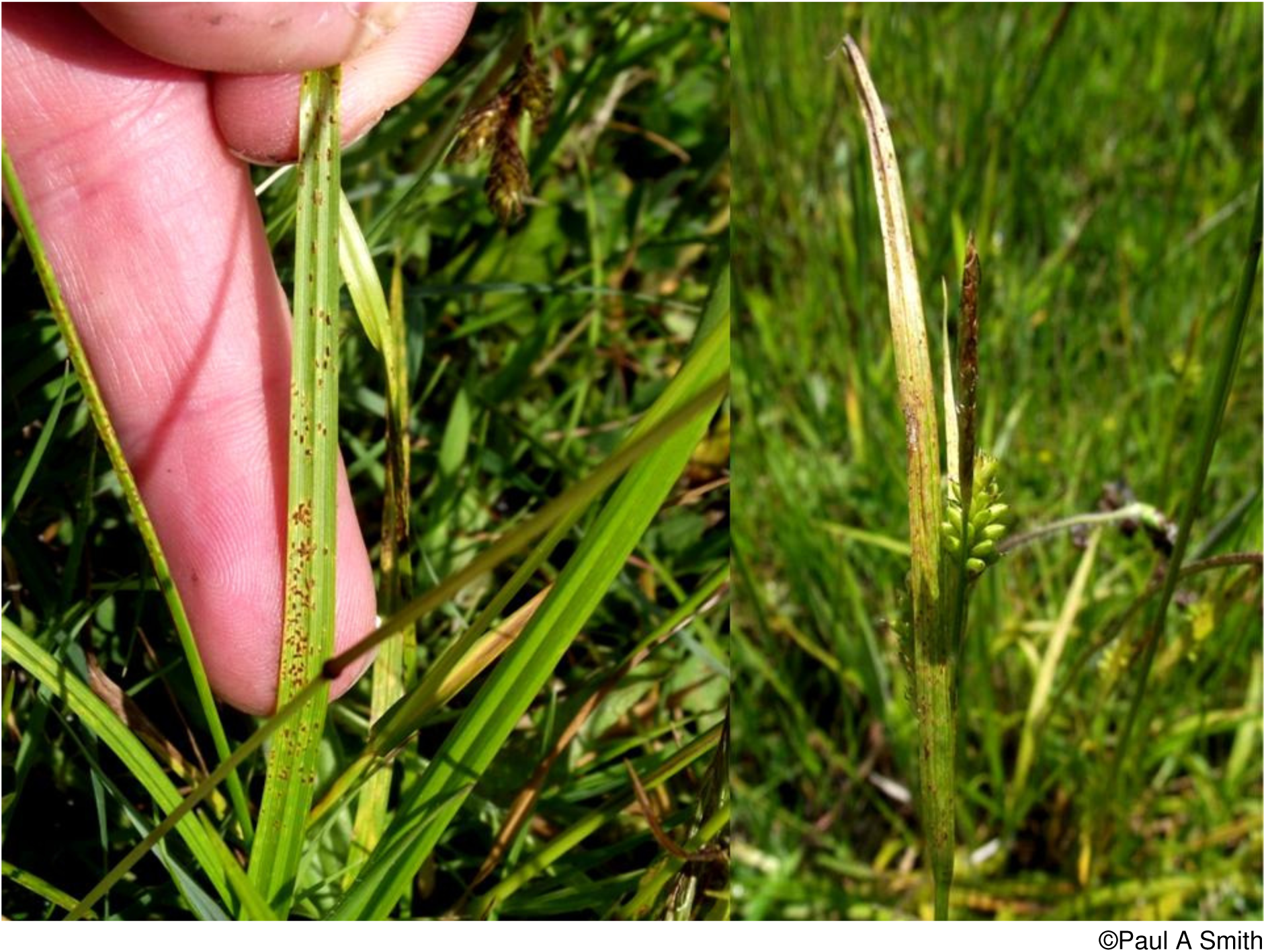

### Puccinia urticata complex

This species alternates between common nettle *Urtica dioica* and sedge *Carex* species. It is not possible to separate the varieties from their aecia on nettles so their abundance can only be measured by their records from sedges.

### Puccinia urticata var. biporula Zwetko

Found very rarely on pale sedge *Carex pallescens* with only five known records from Britain. In Wales it was found in a single site in Glamorganshire (see image on previous page), in a garden border in north Breconshire where a variegated form of the sedge collected from a nearby field was being grown and from near Tre’r-ddol, Cardiganshire. There is on the FRDBI a record from Wester Ross, Scotland and one from Buckinghamshire. In view of the small known population it is considered **Endangered D**.

### Puccinia urticata var. urticae-acutae (Kleb.) Zwetko

Known in Wales from a handful of records on common sedge *Carex nigra*, with a single record from Breconshire, three records from Carmarthenshire and one record from Anglesey. Said by Wilson and Henderson (1966) to be common on this host, it is possible the collection of material may have been avoided by some recorders due to the difficulty of separation of rust species on this host. For the present it is considered **Data Deficient**.

### Puccinia urticata var. urticae-inflatae (Hasler) Zwetko

Found on the upper surface of the leaves of bottle sedge *Carex rostrata* and contrary to the assertion of Wilson and Henderson (1966) that it is frequent on this host in Scotland and England, the FRDBI only lists a single recent record from Scotland (East Perth) and lists no records from England. Paul Smith (pers. comm.), however, reports three recent records from the Outer Hebrides. In Wales it is reported from one site in Radnorshire, four in Carmarthenshire, two in Pembrokeshire, three in Cardiganshire and one in both Montgomeryshire and Anglesey. Whilst natural succession might alter the abundance of bottle sedge, most of its sites are considered to be fairly secure. However the apparent lack of populations in England leaves the Welsh population somewhat isolated and in consequence it is placed in the **Near Threatened** category.

### Puccinia urticata var. urticae-ripariae (Hasler) Zwetko

This rust occurs on greater pond-sedge *Carex riparia*, there being just over a dozen modern records from England on the FRDBI. In Wales it is known only from three sites in Carmarthenshire and two in Cardiganshire. Its host is by no means abundant and is mostly coastal, where a number of its colonies are threatened by rising sea levels. It also probably occurs on Anglesey but this record requires confirmation. This rust is considered to be **Vulnerable D1 & D2**.

### Puccinia veronicae-longifoliae ***Savile***

Found on spiked speedwell *Veronica spicata* ssp. *hybrida*, only a single record is known from Wales. Wilson and Henderson (1966) note a collection made by N. Robertson in 1950 from Denbighshire. The record in Aron (2005) from Vaynol Park, Caernarvonshire in 1972 has proved not to be on this host and must be discounted. With no recent records, this rust is considered to be **Regionally Extinct** in Wales. There are no localised records of this rust from anywhere in Britain on the FRDBI and it is quite possible that this rust is extinct in the wild in Britain.

### Puccinia virgae-aureae (DC.) Lib

A rarely recorded rust of golden rod *Solidago virgaurea*. The FRDBI only lists three recent localities from the north and west of Scotland, a single English record from West Sussex and a single Welsh record from by the Nant Lledr, Dolwyddelan, Caernarvonshire in 1969. In 2008 and 2011 it was found on rocks just under the summit of Snowdon on Clogwyn Du’r Arddu, Caernarvonshire and on a trackside near Carmel in Carmarthenshire in 2013. A search of this latter site in 2014 failed to locate any host plants. Given the small size of the known populations and threats and its remoteness from all other British records, this rust is considered to be **Critically Endangered D2**.

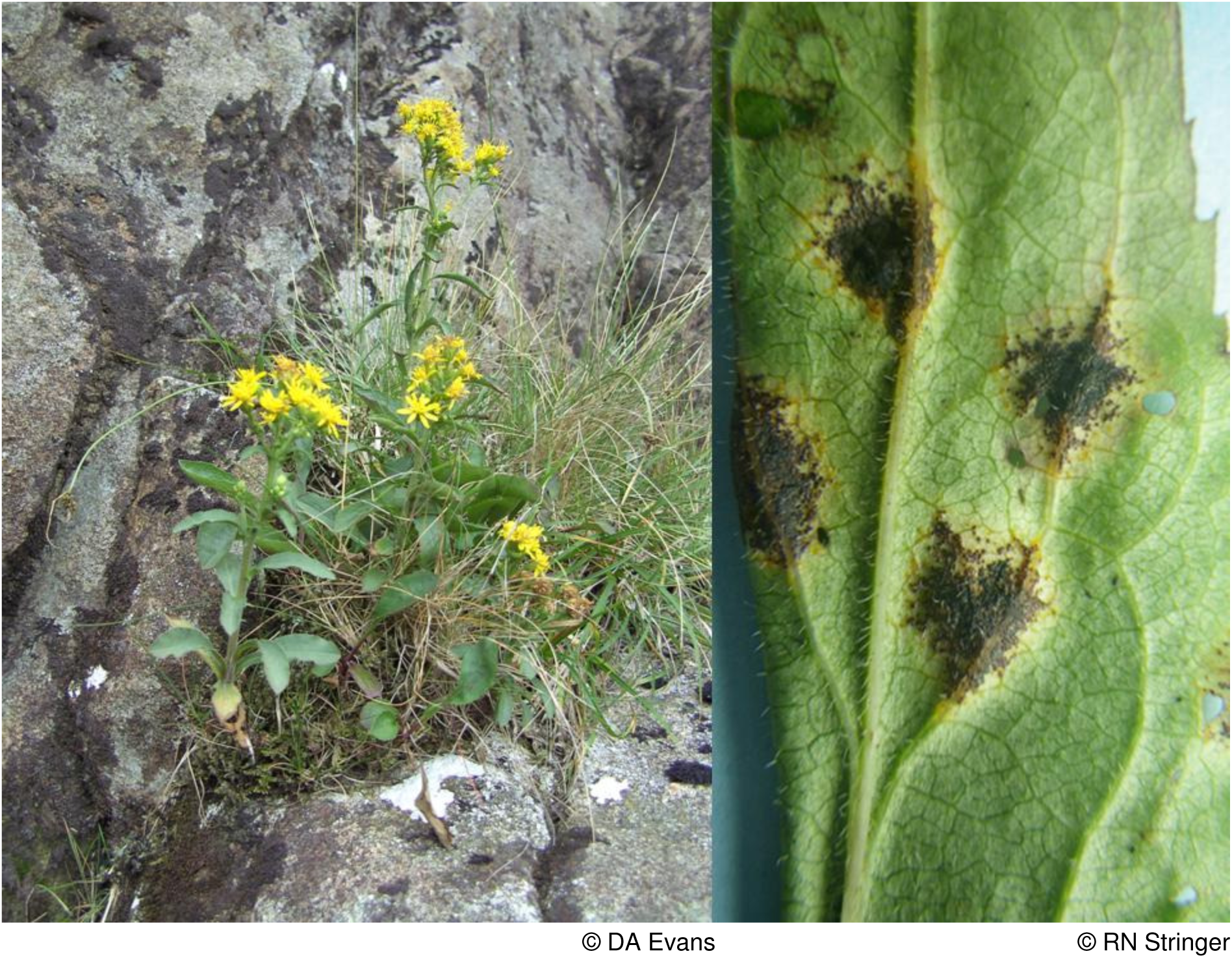
*Puccinia virgae-aureae* on golden rod in Snowdonia

### Pucciniastrum agrimoniae (Dietel) Tranzschel

Despite being a distinctive and readily observed rust on agrimony *Agrimonia eupatoria* it has rarely been reported from Wales. There are old records from the Forden area of Montgomeryshire in the 1870’s and from Dyserth in Flintshire in 1882. More recently the FRDBI notes records made in 2011 from Springdale Farm, Monmouthshire and Morfa Harlech and Shell Island in Merionethshire. In 2001 it was found on the limestone of Llanymynech Hill, Montgomeryshire. There are also recent records from three sites in Caernarvonshire and one in each of Glamorganshire and Denbighshire. This rust is widespread in England and the Welsh populations are western outposts of a species that in England is probably of “Least Concern”. In Wales it qualifies for “Vulnerable” status but its threat status has been downgraded by a category in line with IUCN advice and it is considered to be **Near Threatened**.

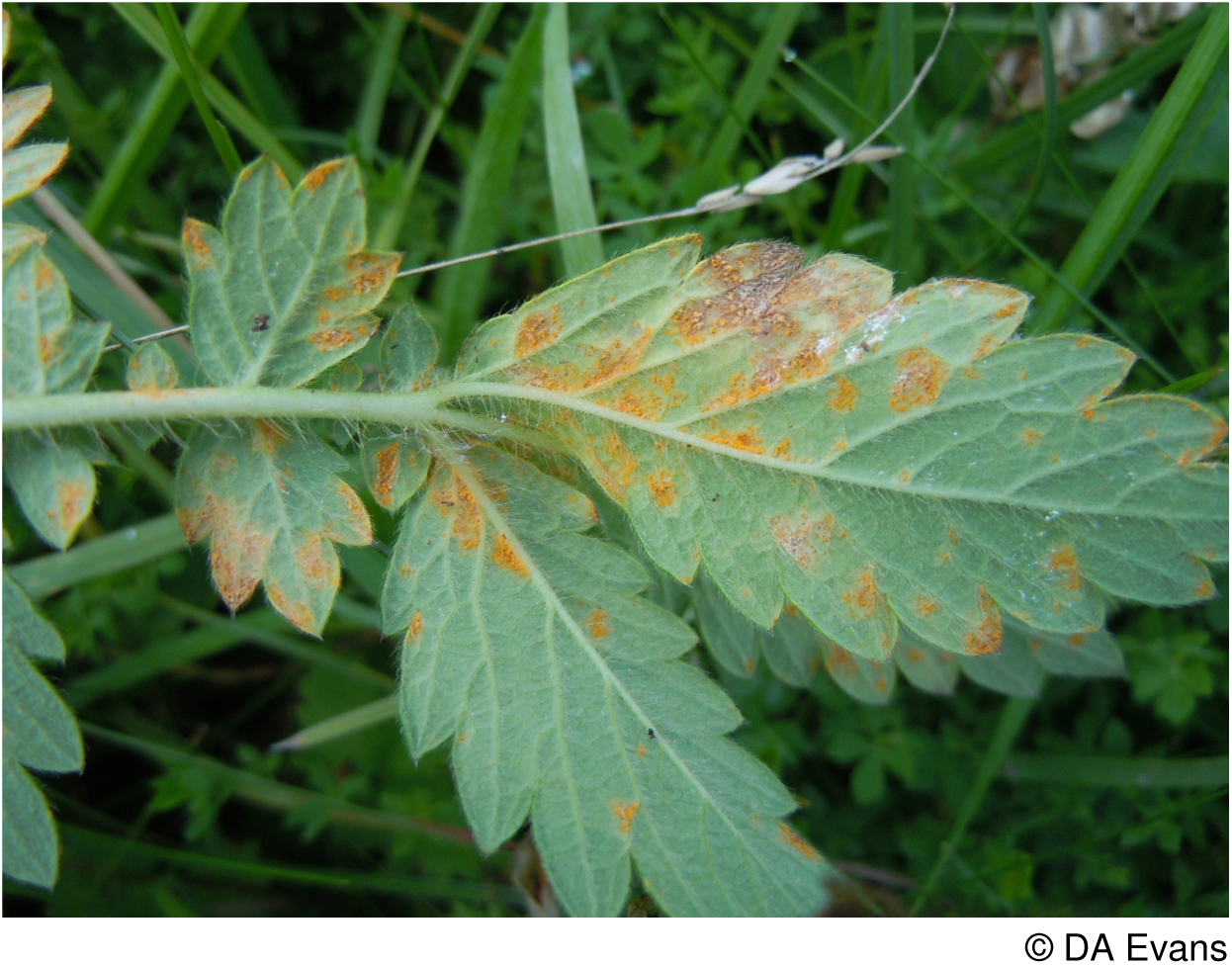
*Pucciniastrum agrimoniae* on agrimony

### Trachyspora intrusa (Grev.) Arthur

The orange aecia often developing over the whole lower leaf surface and the yellowing elongated leaves rising above the uninfected foliage render this a conspicuous species. It is widespread on lady’s-mantle *Alchemilla* species in Scotland and the North of England but is of sparse occurrence in Wales. There are a number of old records from North Wales with the FRDBI noting specimens collected from Cadair Idris, Merioneth in 1841, 1886 and 1944, Cwm Idwal, Caernarvonshire in 1941 and 1950, Bala, Merioneth in 1866 and 1942 and from the Swallow Falls, Betws y Coed, Caernarvonshire in 1885. More recently this rust has been found in five sites on *A. glabra* and once on an undetermined lady’s-mantle species high in the mountains of Snowdonia and once at Caeau Tan y Bwlch on the Lleyn, Caernarvonshire on *A. xanthochlora* in a species-rich pasture. There is also a single record from by Lake Vyrnwy, Montgomeryshire in 1991. In mid and south Wales it is known from two sites in Monmouthshire, two species-rich grassland sites in Radnorshire, three upland sites in the Brecon Beacons in Breconshire, at Parc, Cwm Darran in Glamorganshire and three species-rich grassland sites in Carmarthenshire. Where a host species is recorded the records are equally shared between *A. glabra* and *A. xanthochlora*. Given the loss of many lowland grassland sites (Stevens *et al*. 2010) and the knowledge that both Radnorshire localities are threatened by development and the apparent decline in North Wales, this rust fungus is placed in the **Near Threatened** category.

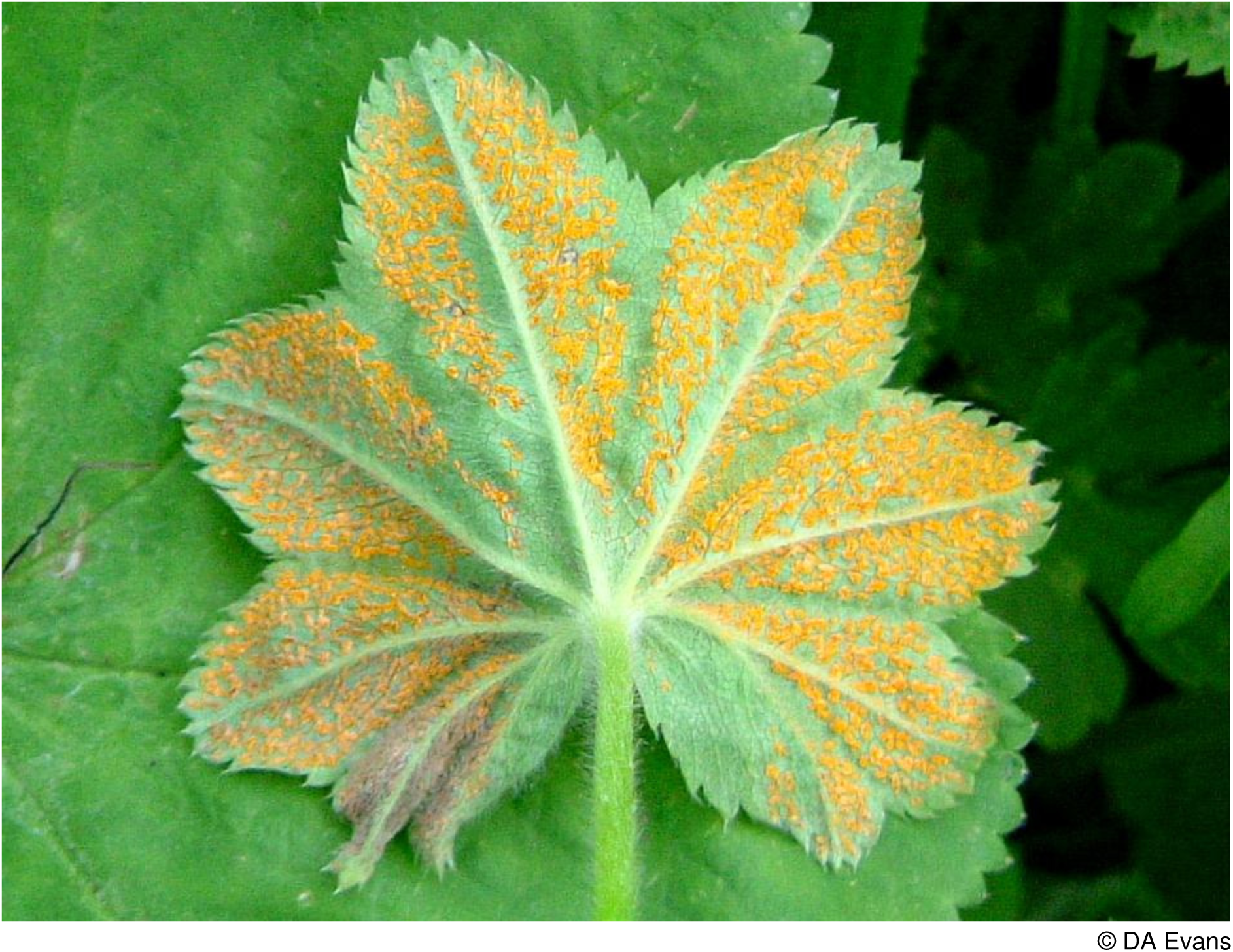
*Trachyspora intrusa* on *Alchemilla xanthochlora*

### Triphragmium filipendulae (Lasch) Pass

Evans *et al*. (2006) consider this rust to be Vulnerable D2 within Britain, with 19 recent records on the FRDBI from a wide range of sites in England and rarely in Scotland. In Wales it has recently been discovered for the first time on dropwort *Filipendula vulgaris* in two sites in North Wales. It occurs in three monads on the limestone of the Great Orme and to the east of Llandudno on the North Wales Wildlife Trust reserve of Bryn Pydew, both in Caernarvonshire. In Wales it is considered to be **Vulnerable B & D1**.

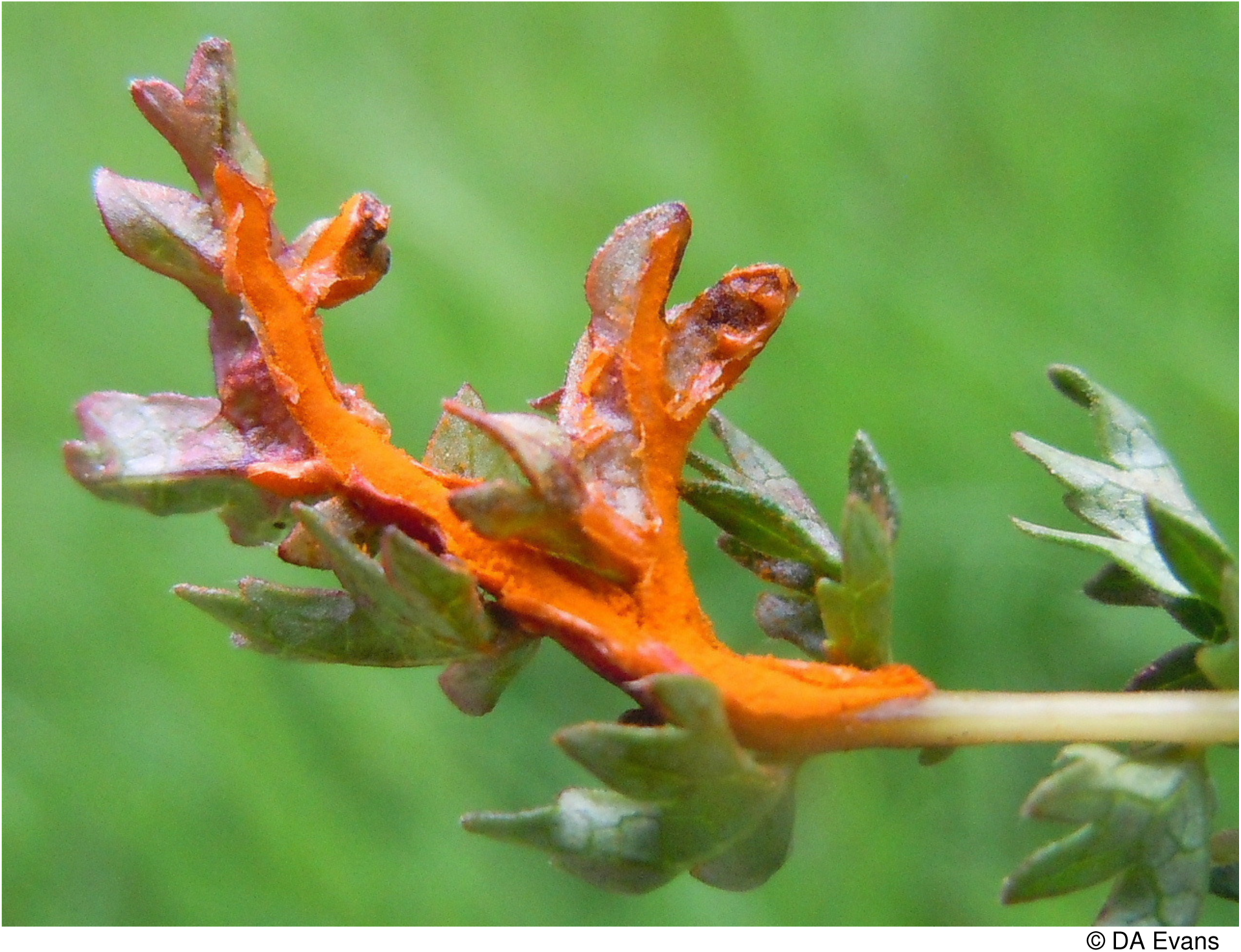
*Triphragmium filipendulae* on dropwort

### Uredo morvernensis Dennis

Described new to science in 1983 from a collection made on wild thyme *Thymus polytrichus* in Morvern, Westerness, Scotland, only three other records of this species have been traced. It was identified in a collection made in NE Galway, Ireland in 1947 and has recently been found in limestone grassland and on an upland Old Red Sandstone cliff in Carmarthenshire. Both of these sites are subject to statutory protection but since the known population in Wales is fewer than 50 individuals it is considered to be **Critically Endangered D2**.

### Uromyces ervi (Wallr.) Westend

Reported only rarely on the hairy tare *Vicia hirsuta*, a common plant of waysides and waste ground. There are recent records from a wide scatter of sites in England on the FRDBI. In Wales it has been found in single sites in Monmouthshire, Breconshire, Radnorshire, Carmarthenshire, Montgomeryshire and Merioneth and in four sites in Cardiganshire, seven sites in Caernarvonshire and four in Anglesey. There is also a single record on smooth tare *V. tetrasperma* from Cardiganshire. In view of the small and scattered populations from fewer than 20 hectads this rust is considered to be **Near Threatened**.

*Uromyces ervi* on hairy tare

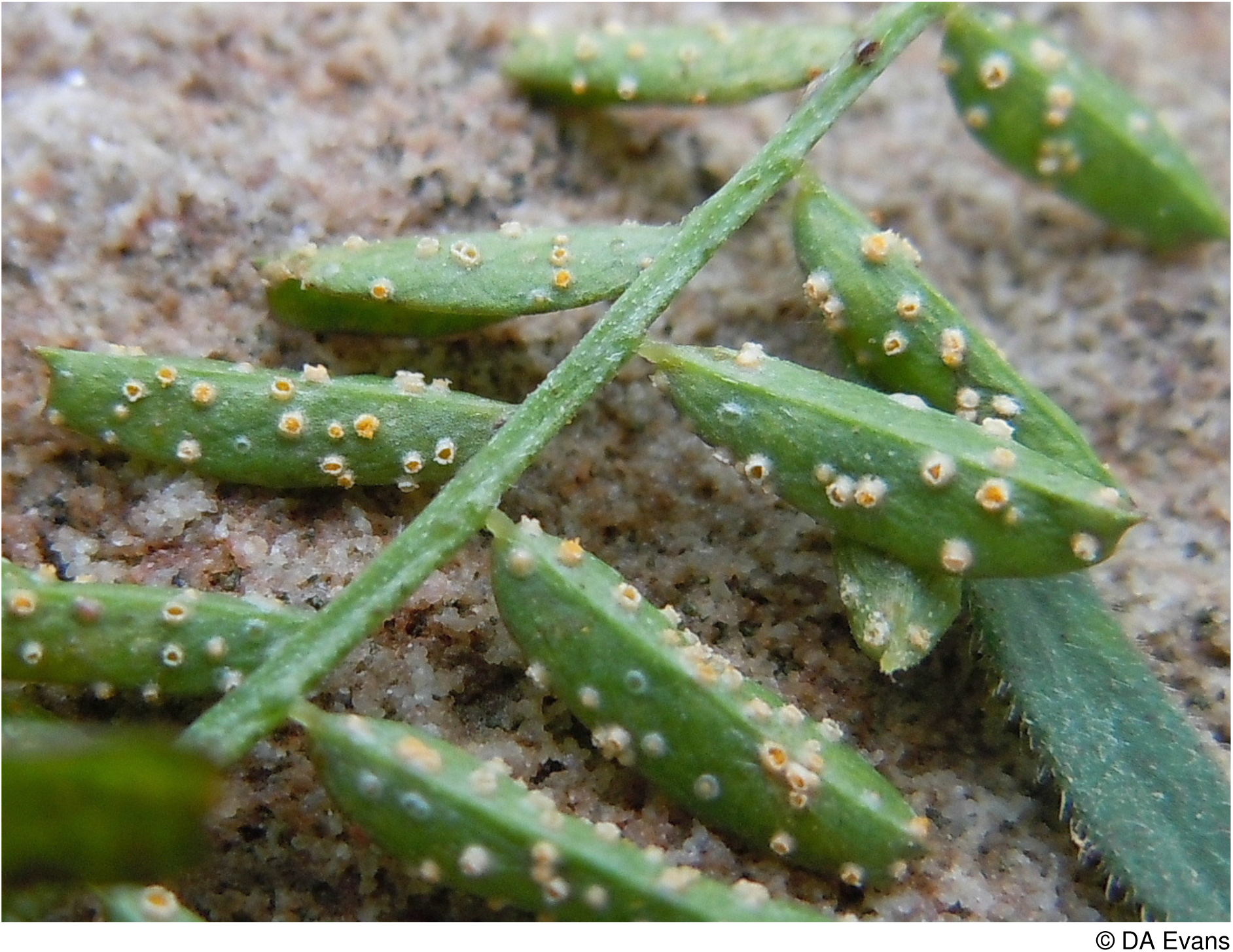

### Uromyces inaequialtus ***Lasch***

Occurring on Nottingham catchfly *Silene nutans*, the only modern records on the FRDBI are from the Channel Isles, though Legon & Henrici (2005) note an unsubstantiated record from Kent. There is but a single record from Wales. It was reported from Prestatyn in Flintshire 1921. The host still occurs in this area but the rust must be considered to be **Regionally Extinct**.

### Uromyces scrophulariae Fuckel

Recorded on water figwort *Scrophularia auriculata* from three sites in Carmarthenshire in 1997 and two sites in 2014; there was only one previous record from Wales (Forden, Montgomeryshire in the 1800’s). The Carmarthenshire populations have in part been strimmed and none of the 1997 populations could be re-found in subsequent years. It has, however, persisted close to the Welsh border in Herefordshire where this rust, first reported in 1951 was re-found in 2009 (FRDBI). There are only 7 recent records from England in the FRDBI. For the time being it is placed in the **Endangered D** category.

### Uromyces sommerfeltii Hyl., Jørst. & Nannf

Discovered new to Britain on goldenrod *Solidago virgaure*a amongst upland rock outcrops in Cardiganshire in 2003. Its single-celled teliospores differentiate it from *Puccinia virgae-aureae* with two-celled teliospores. Only a single plant was seen to be infected and a more recent search in its general area has failed to find any additional populations. In 2008 it was found on this host on Creigiau Gleision at over 550m near Capel Curig and in 2010 on Moel yr Ogof near Beddgelert, both in Caernarvonshire. In view of the tiny known population it is considered to be **Endangered D**.

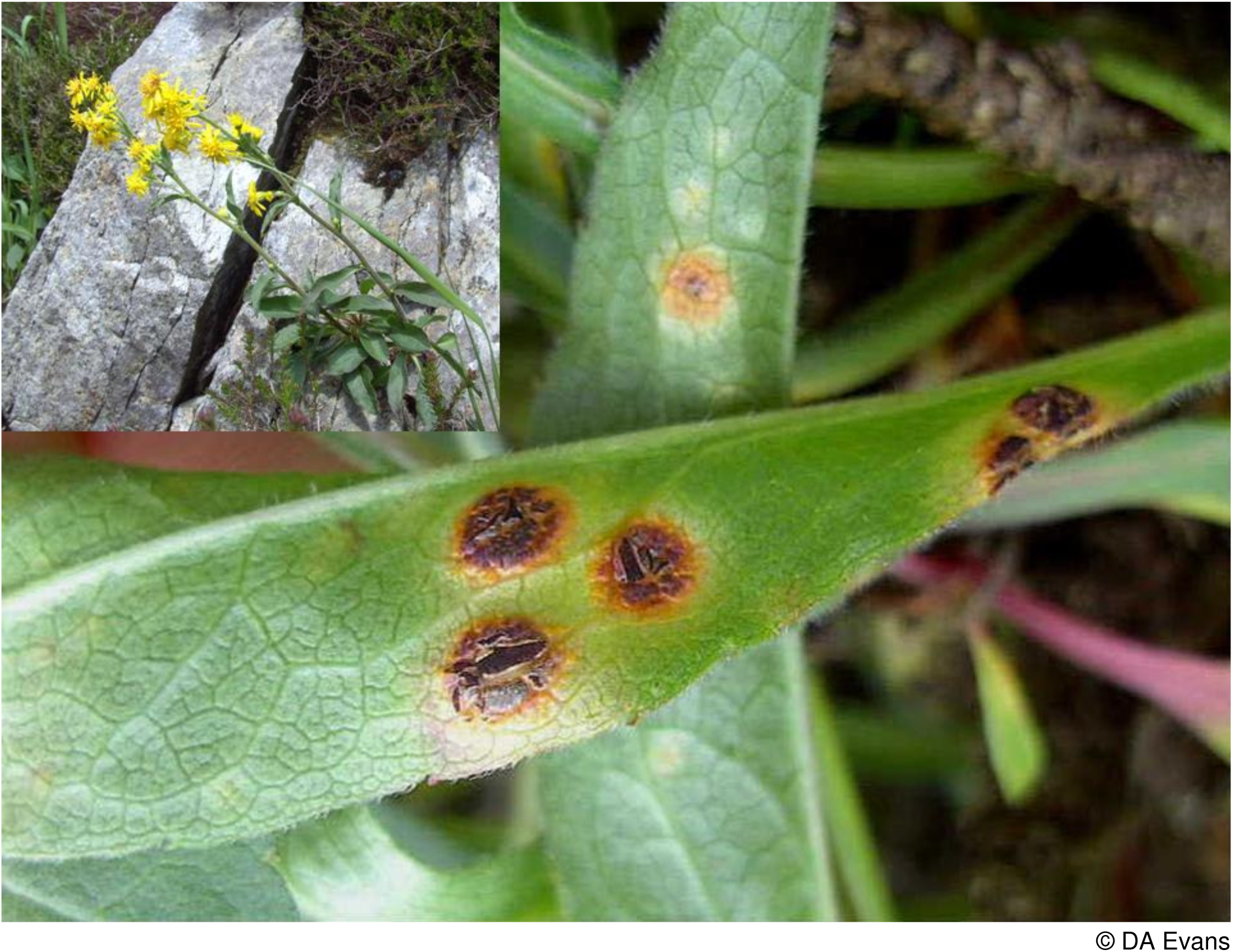

### Uromyces sparsus (J. Kunz & J. C. Schmidt) Lév

Wales supports a notable population of this rust fungus. It is recorded on lesser sea-spurrey *Spergularia marina* from seven sites on the coast of Carmarthenshire, a single site in Glamorgan, four sites on the Menai Straits in Caernarfonshire and two sites on Anglesey.

There are recent records from greater sea-spurrey *S. media* near the mouth of the Afon Leri in Cardiganshire and from the Menai Straits and the Inland Sea near Valley, Anglesey. The FRDBI only notes a single recent record from England in N. Essex, though there are old records from Suffolk, Norfolk, Northumberland and Somerset. Many of the Welsh populations appear to be under threat from coastal erosion and are in consequence considered to be **Near Threatened**.

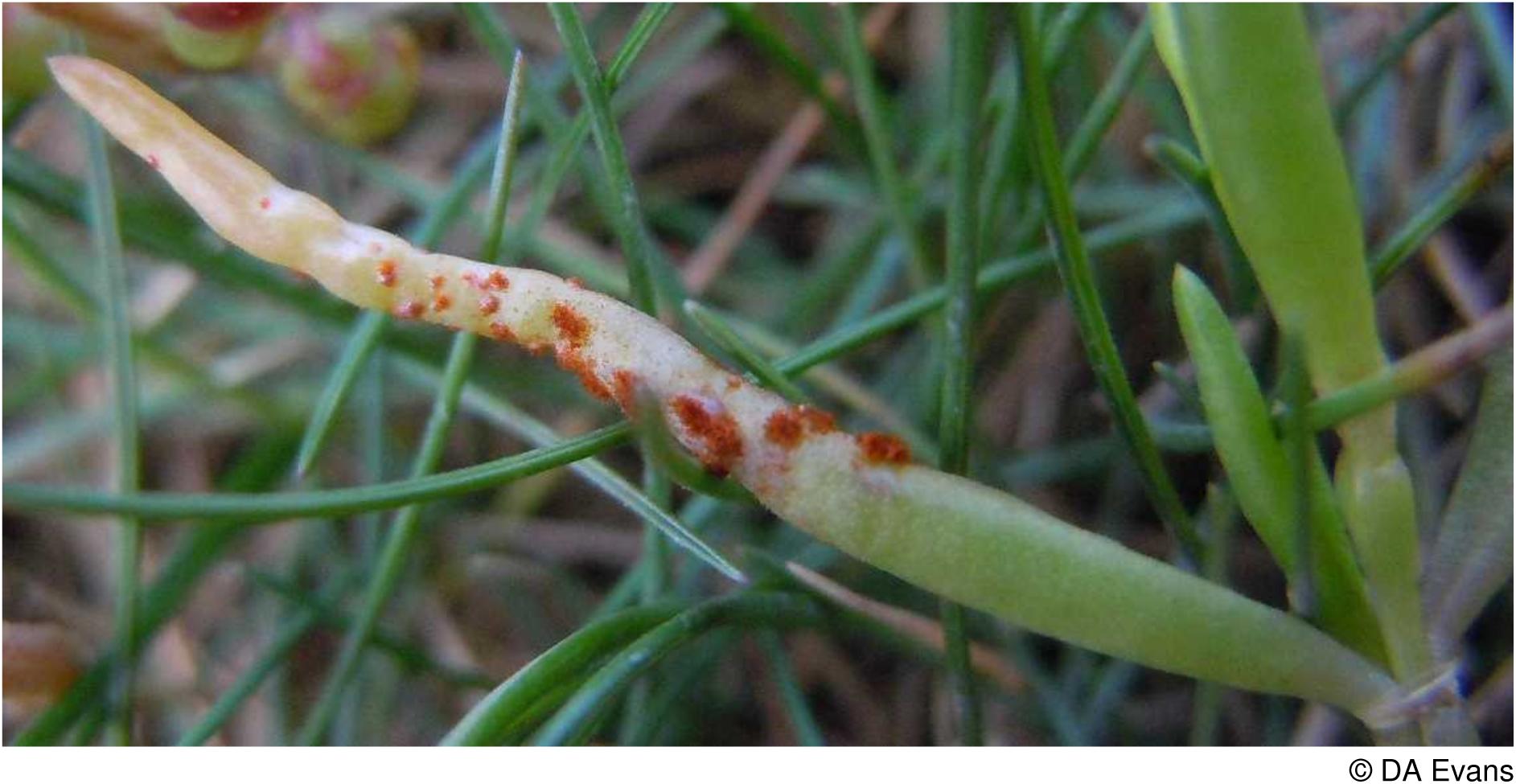
*Uromyces sparsus* on lesser sea-spurrey

### Uromyces viciae-fabae var. orobi (Schumach.) Jørst

Occurring on bitter-vetch *Lathyrus linifolius* var. *montanus,* the FRDBI lists ten recent records from Scotland and five from England. In Wales it is reported from two nearby roadside verges in Radnorshire, two sites in Montgomeryshire and one site each in Merioneth and Caernarvonshire. The host has almost certainly declined with the loss of upland hay meadows (Stevens *et al*. 2010) and this rust is considered to be **Vulnerable D.**

## 10 Rust Red Data List and Census Catalogue

See Section 7 for an explanation of the columns.

| Rust Fungus Taxon | Host Taxon | Cons. Eval. | Vice-county |
| --- | --- | --- | --- |
| *Chrysomyxa abietis* | *Picea abies* (H/N) | **NE** | 44, 46 |
| *Chrysomyxa empetri* | *Empetrum nigrum* | **NT** | 49, 50 |
| *Chrysomyxa empetri* | *Empetrum nigrum* ssp. hermaphroditicum | **NT** | 49 |
| *Chrysomyxa empetri* | *Empetrum nigrum* ssp. *nigrum* | **NT** | 35, 41, 42, 44, 46, [48], 49 |
| *Chrysomyxa pirolata* | *Pyrola rotundifolia* ssp. *maritima* | **CR D** | 52 |
| *Chrysomyxa rhododendri* | *Rhododendron* sp. (H/N) | **NE** | 44 |
| *Coleosporium tussilaginis* | *Campanula rotundifolia* | **LC** | 35, 42-44, 46, 48-51, 52 |
| *Coleosporium tussilaginis* | *Campanula persicifolia* (H) | **NE** | 42, 43, 52 |
| *Coleosporium tussilaginis* | *Euphrasia arctica* ssp. *borealis* | **DD** | 44, 49 |
| *Coleosporium tussilaginis* | *Euphrasia confusa* | **DD** | 44, 52 |
| *Coleosporium tussilaginis* | *Euphrasia confusa* × *nemorosa* | **DD** | 46 |
| *Coleosporium tussilaginis* | *Euphrasia confusa* × *tetraquetra* | **DD** | 46 |
| *Coleosporium tussilaginis* | *Euphrasia nemorosa* | **DD** | 41, 42, 44, 46, 51 |
| *Coleosporium tussilaginis* | *Euphrasia officinalis* | **LC** | 35, 41, 42, 44 |
| *Coleosporium tussilaginis* | *Euphrasia officinalis* ssp. *anglica* | **DD** | 46 |
| *Coleosporium tussilaginis* | *Euphrasia officinalis* ssp. *pratensis* | **DD** | 43, [49] |
| *Coleosporium tussilaginis* | *Euphrasia* sp. | **LC** | 41, 42, 44, 46, 49, 52 |
| *Coleosporium tussilaginis* | *Euphrasia tetraquetra* | **DD** | 42, 46 |
| *Coleosporium tussilaginis* | *Melampyrum pratense* | **LC** | 42, 43, 47-49 |
| *Coleosporium tussilaginis* | *Melampyrum pratense* ssp. *pratense* var. *hyans* | **LC** | 44 |
| *Coleosporium tussilaginis* | *Melampyrum pratense* ssp. *pratense* var. *pratense* | **LC** | 44 |
| *Coleosporium tussilaginis* | *Odontites vernus* | **LC** | 44, 46, 47, 49, 52 |
| *Coleosporium tussilaginis* | *Odontites vernus* ssp. *serotinus* | **LC** | 41, 42, 44, 46, 50, 52 |
| *Coleosporium tussilaginis* | *Parentucellia viscosa* | **DD** | 44, 49 |
| *Coleosporium tussilaginis* | *Petasites fragrans* (H/N) | **LC** | 44 |
| *Coleosporium tussilaginis* | *Petasites hybridus* | **LC** | 35, 41-44, 46, 47, 49, 51, 52 |
| *Coleosporium tussilaginis* | *Petasites japonicus* (H/N) | **LC** | 35, 42 |
| *Coleosporium tussilaginis* | *Pinus contorta* (H/N) | **LC** | 44 |
| *Coleosporium tussilaginis* | *Pinus nigra* ssp. *laricio* (H/N) | **LC** | 35, 44 |
| *Coleosporium tussilaginis* | *Pinus sylvestris* (H) | **LC** | 44 |
| *Coleosporium tussilaginis* | *Rhinanthus minor* | **LC** | 42, 44, 47, 49, 52 |
| *Coleosporium tussilaginis* | *Rhinanthus minor* ssp. *minor* | **LC** | 41, 42, 43, 44, 47 |
| *Coleosporium tussilaginis* | *Rhinanthus minor* ssp. *stenophyllus* | **LC** | 44 |
| *Coleosporium tussilaginis* | *Senecio inaequidens* (N) | **LC** | 44, 46, 49, 50 |
| *Coleosporium tussilaginis* | *Senecio squalidus* (N) | **LC** | 35, 41, 44, 49 |
| *Coleosporium tussilaginis* | *Senecio sylvaticus* | **DD** | 42, 44, 47, 49 |
| *Coleosporium tussilaginis* | *Senecio vulgaris* | **LC** | 35, 42, 44-47, 52 |
| *Coleosporium tussilaginis* | *Senecio vulgaris* var. *vulgaris* | **LC** | 35 |
| *Coleosporium tussilaginis* | *Sonchus arvensis* | **LC** | 42, 43, 47, 49, 52 |
| *Coleosporium tussilaginis* | *Sonchus asper* | **LC** | 41, 43, 47, 49 |
| *Coleosporium tussilaginis* | *Sonchus oleraceus* | **LC** | 35, 44, 46 |
| *Coleosporium tussilaginis* | *Tussilago farfara* | **LC** | 35, 41-44, 46, 47, 49-52 |
| *Cronartium quercuum* | *Quercus cerris* (H/N) | **NT** | 49 |
| Cronartium quercuum | Quercus ilex (H/N) | **NT** | 41 |
| Cronartium quercuum | Quercus petraea | **NT** | 42, 45, 46, 48, 49 |
| Cronartium quercuum | Quercus robur | **NT** | 44, 45 |
| Cronartium ribicola | Ribes nigrum (H/N) | LC | 35, 42, 43, 44, 46, 49 |
| Cronartium ribicola | Ribes rubrum cultivar (H) | LC | 43 |
| Cumminsiella mirabilissima | Mahonia aquifolium (H/N) | LC | 35, 41-44, 46, 47, 49-52 |
| Cumminsiella mirabilissima | Mahonia sp. (H) | LC | 49, 52 |
| Cumminsiella mirabilissima | Mahonia × wageri (H) | LC | 46 |
| Endophyllum euphorbiae-silvaticae | Euphorbia amygdaloides | **CR D** | 35, [47] |
| Endophyllum sempervivi | Sempervivum tectorum (H) | **NE** | 42, [47] |
| Frommeëlla tormentillae | Potentilla anglica | LC | 41, 44, 46 |
| Frommeëlla tormentillae | Potentilla erecta | LC | 41, 42, 44, 46 |
| Frommeëlla tormentillae | Potentilla erecta ssp. erecta | LC | 42-44, 46 |
| Frommeëlla tormentillae | Potentilla reptans | LC | 43, 44, 46, 49 |
| Frommeëlla tormentillae | Potentilla × mixta | LC | 41, 44, 49, 52 |
| Frommeëlla tormentillae | Potentilla × suberecta | LC | 49 |
| Gymnosporangium clavariiforme | Crataegus monogyna | **VU D** | 35, 43, 46, 49, 52 |
| Gymnosporangium clavariiforme | Juniperus communis (H and Wild) | **VU D** | 46, 47, 49, 52 |
| Gymnosporangium clavariiforme | Juniperus sp. (H) | **NE** | 47 |
| Gymnosporangium confusum | Crataegus monogyna | **DD** | 35, 43, 49 |
| Gymnosporangium confusum | Mespilus germanica | **DD** | 49, 52 |
| Gymnosporangium cornutum | Sorbus aucuparia | **DD** | 49 |
| Gymnosporangium sabiniae | Pyrus communis (H) | NE | 49, 52 |
| Gymnosporangium sp. | Crataegus monogyna | **DD** | 42 |
| Hyalopsora polypodii | Cystopteris fragilis | **NT** | 41-44, 47, 48, 50, [51] |
| Kuehneola uredinis | Rubus fruticosus agg. | LC | 35, 41-49, 52 |
| Kuehneola uredinis | Rubus silurum | LC | 48 |
| Kuehneola uredinis | Rubus sp. | LC | 35, 41, 45-52 |
| Melampsora allii-fragilis | Allium ursinum | **VU D2** | 42, 46, 49, 52 |
| Melampsora allii-fragilis | Salix × fragilis | **VU D2** | 43, 44 |
| Melampsora allii-fragilis | Salix pentandra | **VU D2** | 44 |
| Melampsora allii-populina | Arum maculatum | **DD** | 44 |
| Melampsora allii-populina | Populus nigra | **DD** | 35 |
| Melampsora allii-populina | Populus trichocarpa (H) | **DD** | 44 |
| Melampsora arctica | Salix herbacea | **EN D** | 49 |
| Melampsora amygdalinae | Salix triandra | **EN B2** | 44, 46 |
| Melampsora caprearum | Salix aurita | LC | 41, 44-46, 49, 52 |
| Melampsora caprearum | Salix caprea | LC | 35, 41-44, 46, 47, 49-52 |
| Melampsora caprearum | Salix caprea ssp. caprea | LC | 43, 44, 46, 52 |
| Melampsora caprearum | Salix × capreola | LC | 44 |
| Melampsora caprearum | Salix cinerea | LC | 42, 44, 47, 52 |
| Melampsora caprearum | Salix cinerea ssp. oleifolia | LC | 35, 41, 49 |
| Melampsora caprearum | Salix × holosericea | LC | 44, 46 |
| Melampsora caprearum | Salix moupinensis (H) | LC | 44 |
| Melampsora caprearum | Salix × multinervis | LC | 44 |
| Melampsora caprearum | Salix × reichardtii | LC | 41, 44, 50 |
| Melampsora caprearum | Salix sp. | LC | 47, 49, 52 |
| Melampsora caprearum | Salix × sericans | LC | 41, 46 |
| Melampsora epitea | Dactylorhiza fuchsii | **DD** | 52 |
| Melampsora epitea | Dactylorhiza incarnata | **DD** | 52 |
| Melampsora epitea | Dactylorhiza incarnata ssp. coccinea | **DD** | 44, 52 |
| Melampsora epitea | Dactylorhiza maculata | **DD** | 52 |
| Melampsora epitea | Dactylorhiza praetermissa | **DD** | 35, 41 |
| Melampsora epitea | Dactylorhiza purpurella | **DD** | 52 |
| Melampsora epitea | Dactylorhiza sp. | **DD** | 44, 48, 52 |
| Melampsora epitea | Epipactis dunensis | **DD** | 52 |
| Melampsora epitea | Epipactis palustris | **DD** | 48, 52 |
| Melampsora epitea | Euonymus europaeus | LC | 42-44, 46, 49, 52 |
| Melampsora epitea | Salix aurita | LC | 49 |
| Melampsora epitea | Salix caprea | LC | 35, 42, 49, 52 |
| Melampsora epitea | Salix cinerea | LC | 42, 44, 46, 49, 52 |
| Melampsora epitea | Salix cinerea ssp. cinerea | LC | 41, 42, 44, 49, 52 |
| Melampsora epitea | Salix cinerea ssp. oleifolia | LC | 35, 41-46, 48, 49, 52 |
| Melampsora epitea | Salix daphnoides (N) | LC | 44 |
| Melampsora epitea | Salix × fragilis (A) | LC | 35, 41, 44, 47, 49, 52 |
| Melampsora epitea | Salix × mollissima 'Inger' (H) | LC | 46 |
| Melampsora epitea | Salix × mullinervis | LC | 44 |
| Melampsora epitea | Salix phylicifolia (H) | LC | 44 |
| Melampsora epitea | Salix purpurea | LC | 44, 46, 52 |
| Melampsora epitea | Salix repens | **DD** | 44, 48, 52 |
| Melampsora epitea | Salix repens var. argentea | **DD** | 51, 52 |
| Melampsora epitea | Salix × reichardtii | LC | 44, 48 |
| Melampsora epitea | Salix schwerinii × viminalis (H) | LC | 46 |
| Melampsora epitea | Salix × sericans | LC | 44 |
| Melampsora epitea | Salix sp. | LC | 41, 49 |
| Melampsora epitea | Salix triandra | LC | 44 |
| Melampsora epitea | Salix viminalis | LC | 41-44, 45, 49, 52 |
| Melampsora epitea | Salix × smithiana | LC | 44 |
| Melampsora epitea | Undetermined | LC | 50, 51 |
| Melampsora epitea /<br>Puccinia sessilis | Platanthera chlorantha | **DD** | 49 |
| Melampsora euphorbiae | Euphorbia amygdaloides | LC | 44 |
| Melampsora euphorbiae | Euphorbia cyparissias cv. (H) | LC | 44 |
| Melampsora euphorbiae | Euphorbia exigua (A) | LC | 41, 44 |
| Melampsora euphorbiae | Euphorbia helioscopia (A) | LC | 35, 41-49, 52 |
| Melampsora euphorbiae | Euphorbia lathyris (A) | LC | 44, 46, 47, 52 |
| Melampsora euphorbiae | Euphorbia paralias | LC | 44, 46, 48, 52 |
| Melampsora euphorbiae | Euphorbia peplus (A) | LC | 35, 41-44, 46, 48-52 |
| Melampsora euphorbiae | Euphorbia portlandica | LC | 44 |
| Melampsora euphorbiae | Euphorbia sp. | LC | 45, 46, 49 |
| Melampsora euphorbiae | Undetermined | LC | 47 |
| Melampsora hypericorum | Hypericum androsaemum | LC | 35, 41-52 |
| Melampsora hypericorum | Hypericum calycinum (H) | LC | 35, 41-50 |
| Melampsora hypericorum | Hypericum cv. (H) | LC | 47 |
| Melampsora hypericorum | Hypericum × desetangsii | LC | 44 |
| Melampsora hypericorum | Hypericum hircinum (N) | LC | 41, 44, 46 |
| Melampsora hypericorum | Hypericum humifusum | LC | 44 |
| Melampsora hypericorum | Hypericum × inodorum (N) | LC | 35, 41, 44, 48 |
| Melampsora hypericorum | Hypericum pulchrum | LC | 43, 46, 49, 52 |
| Melampsora hypericorum | Hypericum sp. | LC | 44, 47, 49 |
| Melampsora laricis-pentandrae | Salix pentandra | **DD** | 35, 42, 44, 46 |
| Melampsora laricis-populina | Populus balsamifera (H) | LC | 52 |
| Melampsora laricis-populina | Populus × canadensis (H) | LC | 35, 41, 42, 44, 46-50, 52 |
| Melampsora laricis-populina | Populus × canescens (H) | LC | 41, 43, 44, 49 |
| Melampsora laricis-populina | Populus cv. (H) | LC | 41-43, 47, 49, 52 |
| Melampsora laricis-populina | Populus × jackii (H) | LC | 41, 44, 49 |
| Melampsora laricis-populina | Populus nigra | LC | 49 |
| Melampsora laricis-populina | Populus nigra cv. (H) | LC | 35, 44 |
| Melampsora laricis-populina | Populus nigra 'Italica' (H) | LC | 35, 44 |
| Melampsora laricis-populina | Populus nigra ssp. betulifolia | LC | 41, 44, 46 |
| Melampsora laricis-populina | Populus sp. | LC | 44, 47 |
| Melampsora laricis-populina | Populus trichocarpa × balsamifera 'Balsam Spire' (H) | LC | 44 |
| Melampsora laricis-populina | Populus trichocarpa × clone (H) | LC | 49 |
| Melampsora laricis-populina | Populus trichocarpa (H) | LC | 35, 44, 49 |
| Melampsora lini | Linum catharticum | LC | 35, 41-44, 46-49, 52 |
| Melampsora lini var. liniperda | Linum usitatissimum (N) | NE | [45] |
| Melampsora lini var. radiolae *ined* | Radiola linoides | **EX** | [52] |
| Melampsora populnea | Mercurialis perennis | LC | 35, 41-44, 46-52 |
| Melampsora populnea | Populus alba (N) | LC | 41, 43-46, 49, 52 |
| Melampsora populnea | Populus × canescens (N) | LC | 46 |
| Melampsora populnea | Populus sp. | LC | 47, 52 |
| Melampsora populnea | Populus tremula | LC | 41-44, 46, 47, 49, 52 |
| Melampsora populnea | Undetermined | LC | 50 |
| Melampsora ribesii-viminalis | Salix viminalis | **DD** | 41, 44 |
| Melampsora salicis-albae | Salix alba (A) | **NT** | 44-46 |
| Melampsora 'Stem-infecting form' | Salix × holosericea | NE | 44 |
| Melampsora 'Stem-infecting form' | Salix viminalis | NE | 44 |
| Melampsorella caryophyllacearum | Cerastium fontanum ssp. holosteoides | **NT** | 45 |
| Melampsorella caryophyllacearum | Cerastium fontanum ssp. vulgare | **NT** | 42 |
| Melampsorella caryophyllacearum | Cerastium sp. | **NT** | 48 |
| Melampsorella caryophyllacearum | Stellaria graminea | **NT** | 42, [45], 46, 49, 52 |
| Melampsorella caryophyllacearum | Stellaria holostea | NT | 52 |
| Melampsorella caryophyllacearum | Stellaria media | NT | [46], 52 |
| Melampsorella caryophyllacearum | Unknown | NT | [47] |
| Melampsorella symphyti | Symphytum officinale | LC | 35, 42, 45 |
| Melampsorella symphyti | Symphytum sp. (N) | LC | 50 |
| Melampsorella symphyti | Symphytum × uplandicum (N) | LC | 42, 45, 49-52 |
| Melampsorium betulinum | Alnus cordata (H) | LC | 44, 46 |
| Melampsorium betulinum | Alnus glutinosa | LC | 35, 41, 42, 44, 46, 49, 52 |
| Melampsorium betulinum | Alnus glutinosa var. macrocarpa (H) | LC | 46 |
| Melampsorium betulinum | Alnus incana (H) | LC | 42, 44, 46, 49 |
| Melampsorium betulinum | Alnus rubra (H) | LC | 44 |
| Melampsorium betulinum | Alnus sp. | LC | 41, 44 |
| Melampsorium betulinum | Betula ermanii (H) | LC | 41 |
| Melampsorium betulinum | Betula pendula | LC | 41, 43, 44, 48, 49, 51, 52 |
| Melampsorium betulinum | Betula pubescens | LC | 41-44, 46, 48, 49, 52 |
| Melampsorium betulinum | Betula pubescens ssp. pubescens | LC | 35, 41, 42, 44, 46, 49 |
| Melampsorium betulinum | Betula pubescens ssp. tortuosa | LC | 42 |
| Melampsorium betulinum | Betula sp. | LC | 35, 41-43, 48-51 |
| Melampsorium betulinum | Betula utilis ssp. jacquemontii (H) | LC | 46, 49 |
| Melampsorium hiratsukanum | Alnus cordata (H) | LC | 44 |
| Melampsorium hiratsukanum | Alnus glutinosa | LC | 41, 42, 44, 46, 48, 49, 52 |
| Melampsorium hiratsukanum | Alnus incana (H) | LC | 41, 42, 44, 46, 48, 49, 52 |
| Melampsorium hiratsukanum | Alnus rubra (H) | LC | 42, 44 |
| Melampsorium hiratsukanum | Betula pubescens | LC | 44 |
| Milesina blechni | Blechnum spicant | LC | 35, 41-50, 52 |
| Milesina carpatorum | Dryopteris filix-mas | CR D2 | 44, 46 |
| Milesina dieteliana | Polypodium cambricum | LC | 41, 42, 44 |
| Milesina dieteliana | Polypodium interjectum | LC | 41, 42-44, 46, 49, 52 |
| Milesina dieteliana | Polypodium sp. | LC | 48, 49 |
| Milesina dieteliana | Polypodium vulgare | LC | 35, 41-46, 48, 49, 52 |
| Milesina dieteliana | Polypodium × mantoniae | LC | 46 |
| Milesina kriegariana | Athyrium filix-femina | LC | 48, 49 |
| Milesina kriegariana | Dryopteris aemula | LC | 44 |
| Milesina kriegariana | Dryopteris affinis agg. | LC | 44, 49, 50, 52 |
| Milesina kriegariana | Dryopteris affinis ssp. affinis | LC | 44 |
| Milesina kriegariana | Dryopteris borrieri | LC | 44 |
| Milesina kriegariana | Dryopteris carthusiana | LC | 49 |
| Milesina kriegariana | Dryopteris dilatata | LC | 35, 41-49, 50, 52 |
| Milesina kriegariana | Dryopteris filix-mas | LC | 35, 41-49, 52 |
| Milesina murariae | Asplenium ruta-muraria | LC | 35, 41-50, 52 |
| Milesina scolopendrii | Asplenium scolopendrium | LC | 35, 41-50, 52 |
| Milesina scolopendrii | Asplenium scolopendrium var. cristata | LC | 44 |
| Milesina vogesiaca | Polystichum setiferum | VU D1, 2 | 44 |
| Milesina whitei | Polystichum aculeatum | LC | 42-44, 47 |
| Milesina whitei | Polystichum setiferum | LC | 35, 41-47, 49, 50, 52 |
| Miyagia pseudosphaeria | Sonchus arvensis | LC | 41, 44-46, 48, 49, 51, 52 |
| Miyagia pseudosphaeria | Sonchus asper | LC | 35, 41-52 |
| Miyagia pseudosphaeria | Sonchus oleraceus | LC | 35, 41, 43, 44, 46-52 |
| Miyagia pseudosphaeria | Sonchus sp. | LC | 44, 49, 52 |
| Naohidemycetes vacciniorum | Vaccinium × intermedium | LC | 46 |
| Naohidemycetes vacciniorum | Vaccinium myrtillus | LC | 35, 41-51 |
| Naohidemycetes vacciniorum | Vaccinium oxycoccos | LC | 42, 44 |
| Naohidemycetes vacciniorum | Vaccinium sp. | LC | 49 |
| Naohidemycetes vacciniorum | Vaccinium vitis-idaea | LC | 35, 42-44, 46-49 |
| Ochropsora ariae | Anemone nemorosa | NT | 41, 42, 44, 46-47, [48], 49, 50, 52 |
| Phragmidium bulbosum | Rubus caesius | LC | 41, 44-46, 48, 49, 52 |
| Phragmidium bulbosum | Rubus flexuosus | LC | 47 |
| Phragmidium bulbosum | Rubus fruticosus agg. | LC | 35, 41-52 |
| Phragmidium bulbosum | Rubus nemorosus | LC | 46 |
| Phragmidium bulbosum | Rubus polyanthemus | LC | 46 |
| Phragmidium bulbosum | Rubus pyramidalis | LC | 46, 47 |
| Phragmidium bulbosum | Rubus silurum | LC | 44, 46 |
| Phragmidium bulbosum | Rubus sp. | LC | 35, 41, 44, 49-51 |
| Phragmidium bulbosum | Rubus tuberculatus | LC | 46 |
| Phragmidium fragariae | Potentilla sterilis | LC | 35, 41-52 |
| Phragmidium mucronatum | Rosa arvensis | LC | 35-42, 44, 46, 49 |
| Phragmidium mucronatum | Rosa × alba (H) | LC | 46 |
| Phragmidium mucronatum | Rosa × andegavensis | LC | 46 |
| Phragmidium mucronatum | Rosa × bigeneris | LC | 46 |
| Phragmidium mucronatum | Rosa canina | LC | 35, 41-50, 52 |
| Phragmidium mucronatum | Rosa canina (Dumales group) | LC | 44 |
| Phragmidium mucronatum | Rosa cv. (H) | LC | 43-46 |
| Phragmidium mucronatum | Rosa × dumetorum | LC | 46 |
| Phragmidium mucronatum | Rosa laxa (H) | LC | 49 |
| Phragmidium mucronatum | Rosa micrantha | LC | 46 |
| Phragmidium mucronatum | Rosa rugosa (H/N) | LC | 35, 46, 49, 52 |
| Phragmidium mucronatum | Rosa × scabriuscula | LC | 46 |
| Phragmidium mucronatum | Rosa sherardii | LC | 43, 44, 46, 49, 52 |
| Phragmidium mucronatum | Rosa sp. | LC | 35, 41, 42, 44, 46-50, 52 |
| Phragmidium mucronatum | Rosa stylosa | LC | 46 |
| Phragmidium mucronatum | Rosa tomentosa | LC | 44, 46 |
| Phragmidium mucronatum | Rosa × verticillacantha | LC | 46 |
| Phragmidium potentillae | Potentilla anglica | DD | 42, 44, 46 |
| Phragmidium potentillae | Potentilla × suberecta | DD | 42 |
| Phragmidium rosae-pimpinellifoliae | Rosa spinosissima | LC | 41, 44-52 |
| Phragmidium rosae-pimpinellifoliae | Rosa sp. | LC | 47 |
| Phragmidium rubi-idaei | Rubus idaeus | LC | 41-50, 52 |
| Phragmidium sanguisorbae | Poterium sanguisorba ssp. balearicum (N) | LC | 46 |
| Phragmidium sanguisorbae | Poterium sanguisorba ssp. sanguisorba | LC | 41, 42-46, 49-52 |
| Phragmidium tuberculatum | Rosa arvensis | LC | 46 |
| Phragmidium tuberculatum | Rosa canina | LC | 44 |
| Phragmidium tuberculatum | Rosa canina × tomentosa | LC | 46 |
| Phragmidium tuberculatum | Rosa caesia ssp. vogesiaca × canina | LC | 46 |
| Phragmidium tuberculatum | Rosa cultivars | LC | 42, 44, 50 |
| Phragmidium tuberculatum | Rosa gallica "Complicata" (H) | LC | 49 |
| Phragmidium tuberculatum | Rosa × dumalis | LC | 46 |
| Phragmidium tuberculatum | Rosa rugosa × ? "Hollandica" (H) | LC | 44 |
| Phragmidium tuberculatum | Rosa micrantha | LC | 46, 47 |
| Phragmidium tuberculatum | Rosa rugosa (H/N) | LC | 41-43, 45-47, 49, 51, 52 |
| Phragmidium tuberculatum | Rosa sherardii | LC | 42, 44, 46 |
| Phragmidium tuberculatum | Rosa sherardii × tomentosa | LC | 46 |
| Phragmidium tuberculatum | Rosa sp. | LC | 41-49, 51, 52 |
| Phragmidium tuberculatum | Rosa stylosa | LC | 46 |
| Phragmidium tuberculatum | Rosa tomentosa | LC | 46 |
| Phragmidium tuberculatum | Rosa × vericillacantha | LC | 46 |
| Phragmidium violaceum | Rubus albionis | LC | 46 |
| Phragmidium violaceum | Rubus altiarcuatus | LC | 44, 46 |
| Phragmidium violaceum | Rubus amplificatus | LC | 46 |
| Phragmidium violaceum | Rubus anglofuscus | LC | 46 |
| Phragmidium violaceum | Rubus bartonii | LC | 46, 47 |
| Phragmidium violaceum | Rubus boudiccaae | LC | 46 |
| Phragmidium violaceum | Rubus cardiophyllus | LC | 46 |
| Phragmidium violaceum | Rubus dasyphyllus | LC | 46 |
| Phragmidium violaceum | Rubus davisii | LC | 46 |
| Phragmidium violaceum | Rubus dentatifolius | LC | 46, 47 |
| Phragmidium violaceum | Rubus dumnoniensis | LC | 46 |
| Phragmidium violaceum | Rubus effrenatus | LC | 46, 47 |
| Phragmidium violaceum | Rubus flexuosus | LC | 46 |
| Phragmidium violaceum | Rubus fruticosus agg. | LC | 35, 41-52 |
| Phragmidium violaceum | Rubus hastiformis | LC | 46 |
| Phragmidium violaceum | Rubus hylocharis | LC | 46 |
| Phragmidium violaceum | Rubus incurvatus | LC | 46 |
| Phragmidium violaceum | Rubus largificus | LC | 46 |
| Phragmidium violaceum | Rubus leyanus | LC | 46 |
| Phragmidium violaceum | Rubus lindleianus | LC | 46 |
| Phragmidium violaceum | Rubus longus | LC | 46 |
| Phragmidium violaceum | Rubus micans | LC | 46 |
| Phragmidium violaceum | Rubus nemoralis | LC | 46 |
| Phragmidium violaceum | Rubus pampinosus | LC | 46 |
| Phragmidium violaceum | Rubus perdigitatus | LC | 46 |
| Phragmidium violaceum | Rubus polyanthemus | LC | 47 |
| Phragmidium violaceum | Rubus pyramidalis | LC | 46 |
| Phragmidium violaceum | Rubus rubritinctus | LC | 44, 46 |
| Phragmidium violaceum | Rubus silurum | LC | 46 |
| Phragmidium violaceum | Rubus sp. | LC | 35, 41, 42, 45, 46, 48-52 |
| Phragmidium violaceum | Rubus tuberculatus | LC | 46 |
| Phragmidium violaceum | Rubus ulmifolius | LC | 44, 46 |
| Phragmidium violaceum | Rubus ulmifolius hybrid | LC | 46, 47 |
| Phragmidium violaceum | Rubus vestitus | LC | 44, 46, 47 |
| Puccinia acetosae | Rumex acetosa | LC | 41-46, 49, 50-52 |
| Puccinia acetosae | Rumex acetosa ssp. acetosa | LC | 35, 41-46, 48, 49 |
| Puccinia acetosae | Rumex acetosa ssp. biformis | LC | 46 |
| Puccinia acetosae | Rumex acetosella | LC | 41, 44, 46 |
| Puccinia acetosae | Rumex sp. | LC | 49 |
| Puccinia adoxae | Adoxa moschatellina | LC | 35, 41-50, 52 |
| Puccinia aegopodii | Aegopodium podagraria (A) | LC | 35, 41-44, 46-52 |
| Puccinia albescens | Adoxa moschatellina | LC | 35, 42-50, 52 |
| Puccinia allii | Allium ampeloprasum | LC | 41 |
| Puccinia allii | Allium sativum (H) | NE | 42, 43, 46, 49, 52 |
| Puccinia allii | Allium schoenoprasum cv. (H) | LC | 42-44, 46, 48, 49, 52 |
| Puccinia allii | Allium schoenoprasum | LC | 44 |
| Puccinia allii | Allium vineale | LC | 35, 41, 43-47, 49, 50, 52 |
| Puccinia allii | Allium vineale var. compactum | LC | 46, 49 |
| Puccinia angelicae | Angelica sylvestris | **EN D** | [35], 41, 44, 46, [49], 52 |
| Puccinia angelicae | Undetermined | **EN D** | 49 |
| Puccinia annularis | Teucrium scorodonia | LC | 41-50, 52 |
| Puccinia antirrhini | Antirrhinum majus (H) | LC | 35, 41-46, 48-50, 52 |
| Puccinia antirrhini | Antirrhinum molle (H) | LC | 44 |
| Puccinia antirrhini | Antirrhinum sp. (H) | LC | 44 |
| Puccinia apii | Apium graveolens | **RE** | [51] |
| Puccinia arenariae | Dianthus barbatus (H) | LC | 42, 44 |
| Puccinia arenariae | Moehringia trinerva | LC | 42-45, 46, 49, 52 |
| Puccinia arenariae | Sagina apetala | LC | 43, 44, 49 |
| Puccinia arenariae | Sagina procumbens | LC | 44, 46, 49, [52] |
| Puccinia arenariae | Silene dioica | LC | 35, 41-52 |
| Puccinia arenariae | Silene latifolia | LC | 44, 46 |
| Puccinia arenariae | Stellaria graminea | LC | 43, 44, 46, 49, 52 |
| Puccinia arenariae | Stellaria holostea | LC | 44, 46, 48, 49 |
| Puccinia arenariae | Stellaria media | LC | 42, 44, 45, 48, 49 |
| Puccinia arenariae | Stellaria uliginosa | LC | 42, 49, 52 |
| Puccinia arenariae | Undetermined | LC | 35, 41, 47 |
| Puccinia behenis | Silene dioica | LC | 41, 43-52 |
| Puccinia behenis | Silene latifolia | LC | 51 |
| Puccinia behenis | Silene latifolia ssp. alba | LC | 44, 46 |
| Puccinia behenis | Silene robotii "Polly's Favourite"(H) | LC | 49 |
| Puccinia behenis | Silene uniflora | LC | 46, 52 |
| Puccinia behenis | Silene vulgaris | LC | 44, 52 |
| Puccinia betonicae | Stachys officinalis | LC | 42-46, [47], 50, 51 |
| Puccinia bistortae | Conopodium majus | **CR** | 35, [46] |
| Puccinia bistortae | Persicaria bistorta | **CR** | 35 |
| Puccinia brachypodii | Brachypodium sylvaticum | LC | 35, 41-52 |
| Puccinia brachypodii var. arrhenatheri | Arrhenatherum elatius | LC | 35, 42-44, 46, 47, 49, 50, 52 |
| *Puccinia brachypodii* var. *arrhenatheri* | *Arrhenatherum elatius* var. *bulbosum* | LC | 44 |
| *Puccinia brachypodii* var. *arrhenatheri* | *Deschampsia cespitosa* | LC | 35, 42-44, 46, 47 |
| *Puccinia brachypodii* var. *poae-nemoralis* | *Anthoxanthum odoratum* | LC | 35, 41-46, 48, 49, 52 |
| *Puccinia brachypodii* var. *poae-nemoralis* | *Glyceria fluitans* | LC | 44 |
| *Puccinia brachypodii* var. *poae-nemoralis* | *Milium effusum* | LC | 44, 45 |
| *Puccinia brachypodii* var. *poae-nemoralis* | *Phleum arenarium* | LC | 44 |
| *Puccinia brachypodii* var. *poae-nemoralis* | *Phleum pratense* | LC | 44 |
| *Puccinia brachypodii* var. *poae-nemoralis* | *Poa annua* | LC | 41, 44, 45, 48, 49 |
| *Puccinia brachypodii* var. *poae-nemoralis* | *Poa compressa* | LC | 44 |
| *Puccinia brachypodii* var. *poae-nemoralis* | *Poa glauca* | LC | 49 |
| *Puccinia brachypodii* var. *poae-nemoralis* | *Poa humilis* | LC | 49 |
| *Puccinia brachypodii* var. *poae-nemoralis* | *Poa nemoralis* | LC | 43, 44, 47 |
| *Puccinia brachypodii* var. *poae-nemoralis* | *Poa pratensis* | LC | 44, 46 |
| *Puccinia brachypodii* var. *poae-nemoralis* | *Poa* sp. | LC | 44, 50 |
| *Puccinia brachypodii* var. *poae-nemoralis* | *Poa trivialis* | LC | 44 |
| *Puccinia brachypodii* var. *poae-nemoralis* | *Puccinellia distans* | LC | 44 |
| *Puccinia brachypodii* var. *poae-nemoralis* | *Puccinellia maritima* | LC | 41, 44 |
| *Puccinia brachypodii* var. *poae-nemoralis* | *Puccinellia rupestris* | LC | 35, 41 |
| *Puccinia brachypodii* var. *poae-nemoralis* | Undetermined | LC | 52 |
| *Puccinia buxi* | *Buxus balearica* (H) | LC | 49 |
| *Puccinia buxi* | *Buxus microphylla* (H) | LC | 35 |
| *Puccinia buxi* | *Buxus sempervirens* (H) | LC | 35, 41-52 |
| *Puccinia calcitrapae* | *Carduus crispus* | LC | 42, 43 |
| *Puccinia calcitrapae* | *Carduus nutans* | LC | 41, 42, 44, 46 |
| *Puccinia calcitrapae* | *Carduus tenuiflorus* | LC | 41, 46, 49, 52 |
| *Puccinia calcitrapae* | *Carlina vulgaris* | LC | 35, 41-44, 46-49, 51, 52 |
| *Puccinia calcitrapae* | *Centaurea nigra* | LC | 35, 41, 42, 44, 46, 48-50, 52 |
| *Puccinia calcitrapae* | *Centaurea scabiosa* | LC | 44 |
| *Puccinia calcitrapae* | *Cirsium acaule* | LC | 42 |
| *Puccinia calcitrapae* | *Cirsium dissectum* | LC | 42 |
| *Puccinia calcitrapae* | *Cirsium palustre* | LC | 35, 42-46, 48-50, 52 |
| *Puccinia calcitrapae* | Undetermined | LC | 47, 49 |
| *Puccinia calthae* | *Caltha palustris* | NT | 43-46, 48, 49, 52 |
| *Puccinia calthicola* | *Caltha palustris* | LC | 42-44, [45], 46-47, 49, 52 |
| *Puccinia campanulae* | *Campanula rotundifolia* | DD | 49, 52 |
| *Puccinia campanulae* | *Jasione montana* | DD | [46] |
| *Puccinia cancellata* | *Juncus acutus* | DD | 41, 44, 48 |
| *Puccinia cancellata* | *Juncus maritimus* | DD | 41, 44, 46 |
| *Puccinia caricina* (s.l.) | *Carex aquatilis* | LC | 44, 46 |
| *Puccinia caricina* (s.l.) | *Carex binervis* | LC | 35, 42-44, 46-49, 52 |
| *Puccinia caricina* (s.l.) | *Carex caryophyllea* | LC | 41, 49 |
| *Puccinia caricina* (s.l.) | *Carex × fulva* | LC | 42, 46 |
| *Puccinia caricina* (s.l.) | *Carex echinata* | LC | 43 |
| *Puccinia caricina* (s.l.) | *Carex hostiana* | LC | 41, 42, 46, 49, 52 |
| *Puccinia caricina* (s.l.) | *Carex laevigata* | LC | 41, 42, 44, 46, 47, 52 |
| *Puccinia caricina* (s.l.) | *Carex nigra* | LC | 42, 43, 46 |
| *Puccinia caricina* (s.l.) | *Ribes nigrum* (N) | LC | 44 |
| *Puccinia caricina* (s.l.) | *Ribes rubrum* | LC | 52 |
| *Puccinia caricina* (s.l.) | *Ribes uva-crispa* (H/A) | LC | 42, 44, 48, 49, 52 |
| *Puccinia caricina* var. *magnusii* | *Carex riparia* | NT | 35, 44-46, 52 |
| *Puccinia caricina* var. *pringsheimiana* | *Ribes uva-crispa* (H/A) | DD | 44, [48], [49], [52] |
| *Puccinia caricina* var. *pringsheimiana* | *Carex acuta* | DD | 42, 43, 52 |
| *Puccinia caricina* var. *ribesii-pendulae* | *Carex pendula* | LC | 35, 41-47, 49-52 |
| *Puccinia caricina* var. *ribis-nigri-paniculatae* | *Carex paniculata* | NT | 41, 43, 44, 49, 50, 52 |
| *Puccinia chaerophylli* | *Anthriscus sylvestris* | DD | 44, 46, [52] |
| *Puccinia chaerophylli* | *Chaerophyllum temulum* | DD | 42, 44 |
| *Puccinia chaerophylli* | *Myrrhis odorata* (H/N) | LC | 35, 42-44, 46, 47, 49, 50, 52 |
| *Puccinia chrysosplenii* | *Chrysosplenium alternifolium* | LC | 42 |
| *Puccinia chrysosplenii* | *Chrysosplenium oppositifolium* | LC | 35, 42-45, 48, 49, 50, 52 |
| *Puccinia chrysosplenii* | *Chrysosplenium* sp. | LC | 47, 49 |
| *Puccinia circaeae* | *Circaea lutetiana* | LC | 35, 41-52 |
| *Puccinia circaeae* | *Circaea* sp. | LC | 41, 49, 50 |
| *Puccinia circaeae* | *Circaea × intermedia* | LC | 44, 46, 49 |
| *Puccinia cladii* | *Cladium mariscus* | EN D | 41 |
| *Puccinia cnici* | *Cirsium eriophorum* | LC | 44 |
| *Puccinia cnici* | *Cirsium palustre* | LC | 46 |
| *Puccinia cnici* | *Cirsium vulgare* | LC | 35, 41-46, 48-52 |
| *Puccinia cnici-oleracei* | *Achillea millefolium* | LC | 42-44, 46, 48-50, 52 |
| *Puccinia cnici-oleracei* | *Achillea ptarmica* | LC | 44 |
| *Puccinia cnici-oleracei* | *Aster tripolium* | LC | 41, 44 |
| *Puccinia cnici-oleracei* | *Cirsium dissectum* | LC | 35 |
| *Puccinia cnici-oleracei* | *Cirsium palustre* | LC | 35-44, 46, 48-50, 52 |
| *Puccinia cnici-oleracei* | *Matricaria* sp. | LC | 52 |
| *Puccinia commutata* | *Valeriana officinalis* | LC | 41, 43, 45, 46, 49, 52 |
| *Puccinia conii* | *Conium maculatum* (A) | NT | 35, [41], [45], [47], 49, 52 |
| *Puccinia coronata* | *Agrostis canina* | LC | 42, 43 |
| *Puccinia coronata* | *Agrostis capillaris* | LC | 35, 41-44, 46, 48, 49, 52 |
| *Puccinia coronata* | *Agrostis gigantea* (A) | LC | 42 |
| Puccinia coronata | Agrostis stolonifera | LC | 35, 41-46, 48, 49, 50 |
| Puccinia coronata | Alopecurus geniculatus | LC | 44 |
| Puccinia coronata | Alopecurus myosuroides (A) | LC | 44 |
| Puccinia coronata | Alopecurus pratensis | LC | 44 |
| Puccinia coronata | Arrhenatherum elatius | LC | 35, 41-44, 46, 47, 49-52 |
| Puccinia coronata | Arrhenatherum elatius var. bulbosum | LC | 41-47, 49, 50, 52 |
| Puccinia coronata | Avena fatua (A) | LC | 44, 46 |
| Puccinia coronata | Avena sativa (H/ Cas) | LC | 42, 44, 46, 49 |
| Puccinia coronata | Avena sterilis subsp. ludoviciana (H) | LC | 46 |
| Puccinia coronata | Avena sp. (H ?) | LC | 41, 45, 52 |
| Puccinia coronata | Bromopsis ramosa | LC | 44 |
| Puccinia coronata | Calamagrostis epigejos | LC | 41 |
| Puccinia coronata | Calamagrostis × acutiflora 'Karl Foerster' (H) | LC | 44 |
| Puccinia coronata | Deschampsia cespitosa | LC | 41, 42, 44, 46, 49, 52 |
| Puccinia coronata | Elytrigia repens ssp. repens | LC | 44 |
| Puccinia coronata | Frangula alnus | LC | 42, 44, 46, 48, 49, 52 |
| Puccinia coronata | Glyceria maxima | LC | 44 |
| Puccinia coronata | Holcus lanatus | LC | 35, 41-49, 50, 52 |
| Puccinia coronata | Holcus mollis | LC | 35, 41-46, 48-50, 52 |
| Puccinia coronata | Holcus sp. | LC | 35, 41, 44 |
| Puccinia coronata | Lolium multiflorum | LC | 44 |
| Puccinia coronata | Lolium perenne | LC | 41-46, 49, 52 |
| Puccinia coronata | Phalaris arundinacea | LC | 41 |
| Puccinia coronata | Phleum pratense | LC | 44 |
| Puccinia coronata | Poa annua | LC | 44 |
| Puccinia coronata | Poa compressa | LC | 41, 44 |
| Puccinia coronata | Poa nemoralis | LC | 44 |
| Puccinia coronata | Poa pratensis ssp. irrigata | LC | 46, 48, 49, 52 |
| Puccinia coronata | Poa trivialis | LC | 44 |
| Puccinia coronata | Rhamnus cathartica | LC | 44, 52 |
| Puccinia coronata | Schedonorus arundinaceus | LC | 41, 44, 46 |
| Puccinia coronata | Schedonorus giganteus | LC | 35, 41-44, 46-49, 50, 52 |
| Puccinia coronata | Schedonorus pratensis | LC | 44 |
| Puccinia coronata var. gibberosa | Festuca altissima | **CR D** | 43, 46 |
| Puccinia crepidicola | Crepis biennis (N) | **NT** | 49 |
| Puccinia crepidicola | Crepis capillaris | **NT** | 35, 42-44, 46, [48], 49, 52 |
| Puccinia crepidicola | Crepis vesicaria (N) | **NT** | 44, 52 |
| Puccinia difformis | Galium aparine | **NT** | 41, 44, [45], 46, 48-50, 52 |
| Puccinia dioicae (s.l.) | Carex leporina | LC | 41-44, 46, 49 |
| Puccinia dioicae (s.l.) | Carex demissa | LC | 43, 44, 46, 48, 49, 52 |
| Puccinia dioicae (s.l.) | Carex lepidocarpa | LC | 52 |
| Puccinia dioicae (s.l.) | Carex dioica | **EN D** | 42, 49 |
| Puccinia dioicae | Cirsium palustre | **EN D** | 48, 49 |
| Puccinia dioicae var. extensicola | Aster tripolium | LC | 41, 44, 45, 46, 49, 52 |
| Puccinia dioicae var. extensicola | Carex extensa | LC | 44-46, 49, 52 |
| *Puccinia dioicae* var. *schoeleriana* | *Carex arenaria* | **VU D** | 44, 52 |
| *Puccinia dioicae* var. *schoeleriana* | *Senecio jacobaea* | **VU D** | 44, 48, 52 |
| *Puccinia dioicae* var. *schoeleriana* | *Senecio* sp. | **VU D** | 35 |
| *Puccinia elymi* | *Ammophila arenaria* | **VU D2** | 41, 45, 46, 49 |
| *Puccinia elymi* | *Leymus arenarius* | **VU D2** | 44, 49, 51, 52 |
| *Puccinia epilobii* | *Epilobium ciliatum* (N) | LC | 41, 46 |
| *Puccinia epilobii* | *Epilobium hirsutum* | LC | 44, 46 |
| *Puccinia epilobii* | *Epilobium lanceolatum* | LC | 44 |
| *Puccinia epilobii* | *Epilobium montanum* | LC | 41, 44, 46, 49, 52 |
| *Puccinia epilobii* | *Epilobium obscurum* | LC | 35, 41, 44, 46, 49, 52 |
| *Puccinia epilobii* | *Epilobium palustre* | LC | 35, 42, 43, [49] |
| *Puccinia epilobii* | *Epilobium* sp. | LC | 41, 42, 44, 46, [47], 49, 52 |
| *Puccinia eriophorae* | *Solidago virgaurea* | **EN D** | 49 |
| *Puccinia eriophorae* | *Trichophorum cespitosum* agg. | **EN D** | 49 |
| *Puccinia fergussonii* | *Viola palustris* | **EN D** | 41, 42, 44, 46, 48, 49, 52 |
| *Puccinia fergussonii* | *Viola palustris* ssp. *palustris* | **EN D** | 46, 49 |
| *Puccinia festucae* | *Festuca "glauca"* (H) | LC | 42, 44 |
| *Puccinia festucae* | *Festuca ovina* | LC | 42, 44, 46, 50 |
| *Puccinia festucae* | *Festuca rubra* | LC | 41, 44-46, 49, 51, 52 |
| *Puccinia festucae* | *Festuca rubra* ssp. *junceae* | LC | 45, 46 |
| *Puccinia festucae* | *Festuca rubra* ssp. *litoralis* | LC | 46 |
| *Puccinia festucae* | *Festuca rubra* ssp. *rubra* | LC | 35, 41, 42, 44-46, 48, 49 |
| *Puccinia festucae* | *Festuca vivipara* | LC | 42 |
| *Puccinia festucae* | *Lonicera cultivar* (H) | LC | 49, 52 |
| *Puccinia festucae* | *Lonicera periclymenum* | LC | 41, 43, 47, 48, 49, 52 |
| *Puccinia galii-cruciatae* | *Cruciata laevipes* | **EN B2,D** | 41, 42, 44 |
| *Puccinia galii-verni* | *Cruciata laevipes* | LC | 35, 42-44, 50 |
| *Puccinia galii-verni* | *Galium saxatile* | LC | 35, 41, 43, 45, 46, 49, 50 |
| *Puccinia galii-verni* | *Galium* sp. | LC | 52 |
| *Puccinia galii-verni* | *Galium uliginosum* | LC | 42, 46 |
| *Puccinia galii-verni* | *Galium verum* | LC | 48 |
| *Puccinia galii-verni* | Undetermined | LC | 47 |
| *Puccinia glechomatis* | *Glechoma hederacea* | LC | 35, 41-52 |
| *Puccinia glomerata* | *Senecio jacobaea* | **NT** | 44, 46, 47, 49-52 |
| *Puccinia glomerata* | *Senecio aquaticus* | **NT** | 47 |
| *Puccinia graminis* | Undetermined | DD | [48], 50 |
| *Puccinia graminis* | *Agrostis capillaris* | DD | 41, 44, 46, [48] |
| *Puccinia graminis* | *Agrostis gigantea* (A) | DD | 42 |
| *Puccinia graminis* | *Agrostis stolonifera* | DD | 41, 44 |
| *Puccinia graminis* | *Anthoxanthum odoratum* | DD | 35, 41, 43, 44 |
| *Puccinia graminis* | *Arrhenatherum elatius* | DD | 42 |
| *Puccinia graminis* | *Arrhenatherum elatius* var. *bulbosum* | DD | 41, 42, 44, 46 |
| *Puccinia graminis* | *Avena* sp. (H) | DD | [49] |
| *Puccinia graminis* | *Berberis thunbergii* f. *atropurpurea* (H) | DD | 46 |
| *Puccinia graminis* | *Berberis vulgaris* (H) | DD | 42-44, 46, 47 |
| *Puccinia graminis* | *Briza media* | DD | 44 |
| *Puccinia graminis* | *Dactylis glomerata* | DD | 41, 42, 44, 45 |
| *Puccinia graminis* | *Deschampsia cespitosa* | DD | 41, 44 |
| *Puccinia graminis* | *Elytrigia repens* | DD | [35], [49] |
| *Puccinia graminis* | *Hordeum marinum* | DD | 41 |
| *Puccinia graminis* | *Lolium perenne* | DD | 35-44, 46, 48, 49, 52 |
| *Puccinia graminis* | *Mahonia aquifolium* | DD | 41 |
| *Puccinia graminis* | *Milium effusum* | DD | 42 |
| *Puccinia graminis* | *Phleum pratense* | DD | 44, 46 |
| *Puccinia graminis* | *Poa annua* | DD | 35 |
| *Puccinia graminis* | *Schedonorus arundinaceus* | DD | 44, 45 |
| *Puccinia graminis* | *Schedonorus giganteus* | DD | 43, 44 |
| *Puccinia graminis* | *Triticum* sp. (H) | DD | 46 |
| *Puccinia heraclii* | *Heracleum sphondylium* | **RE** | [46], [48], [49] |
| *Puccinia heucherae* | *Heuchera* cultivars (H) | NE | 44, 49, 52 |
| *Puccinia hieracii* | *Centaurea nigra* | LC | 35, 41-44, 46, 47-49, 52 |
| *Puccinia hieracii* | *Centaurea scabiosa* | LC | 41, 44 |
| *Puccinia hieracii* | *Cichorium intybus* | LC | 46 |
| *Puccinia hieracii* | *Hieracium pulchrius* | LC | 42 |
| *Puccinia hieracii* | *Hieracium repandulare* | LC | 42 |
| *Puccinia hieracii* | *Hieracium sabaudum* | LC | 42, 44, 46, 49 |
| *Puccinia hieracii* | *Hieracium* sp. | LC | 35, 41-44, 46, 47, 49-52 |
| *Puccinia hieracii* | *Hieracium umbellatum* | LC | 43, 47, 49 |
| *Puccinia hieracii* | *Hieracium vagum* | LC | 44, 46, 49 |
| *Puccinia hieracii* | *Leontodon hispidus* | LC | 35, 41-47, 50, 52 |
| *Puccinia hieracii* | *Leontodon* sp. | LC | 43, 49 |
| *Puccinia hieracii* | *Leontodon saxatilis* | LC | 44 |
| *Puccinia hieracii* | *Picris hieracioides* | LC | 41 |
| *Puccinia hieracii* | *Scorzoneroideis autumnalis* | LC | 35, 42, 43, 46, 49 |
| *Puccinia hieracii* | *Serratula tinctoria* | LC | 35, 42, 46, 49, 52 |
| *Puccinia hieracii* | *Taraxacum alatum* | LC | 49 |
| *Puccinia hieracii* | *Taraxacum eckmanii* | LC | 49 |
| *Puccinia hieracii* | *Taraxacum officinale* agg. | LC | 35, 41-52 |
| *Puccinia hieracii* | *Taraxacum pannulatum* | LC | 49 |
| *Puccinia hieracii* | *Taraxacum sellandii* | LC | 49 |
| *Puccinia hieracii* | *Taraxacum* sp. | LC | 35, 46, 48 |
| *Puccinia hieracii* | Undetermined | LC | 47, 49 |
| *Puccinia hieracii* var. *hypochaeridis* | *Hypochaeris radicata* | LC | 41-49, 52 |
| *Puccinia hieracii* var. *hypochaeridis* | *Hypochaeris* sp. | LC | 52 |
| *Puccinia hieracii* var. *piloselloidarum* | *Pilosella aurantiaca* (N) | LC | 44, 49 |
| *Puccinia hieracii* var. *piloselloidarum* | *Pilosella officinarum* | LC | 35, 41-44, 46, 49, 52 |
| *Puccinia hordei* | *Holcus lanatus* | LC | 41, 44, 46-48, 52 |
| *Puccinia hordei* | *Holcus mollis* | LC | 41, 42, 44, 46-49 |
| *Puccinia hordei* | *Holcus* sp. | LC | 35, 42, 49-51 |
| *Puccinia hordei* | *Hordeum distichon* (H) | LC | 46 |
| *Puccinia hordei* | *Hordeum murinum* (A) | LC | 41, 44, 49 |
| *Puccinia hordei* | *Hordeum murinum* ssp. *murinum* (A) | LC | 41, 44 |
| Puccinia hordei | Hordeum secalinum | LC | 42, 44, 52 |
| Puccinia hordei | Hordeum vulgare | LC | 52 |
| Puccinia hordei | Lolium multiflorum | LC | 35, 46 |
| Puccinia hordei | Lolium perenne | LC | 41, 44 |
| Puccinia hordei | Trisetum flavescens | LC | 35, 41, 44 |
| Puccinia hordei | Vulpia bromoides | LC | 35, 42, 43, 44, 46, 49, 52 |
| Puccinia hordei | Vulpia myuros (A) | LC | 44, 46 |
| Puccinia horiana | Chrysanthemum cultivar (H) | NE | 42, 44 |
| Puccinia hydrocotyles | Hydrocotyle vulgaris | **EN D** | 44, 46, 52 |
| Puccinia hystrium | Tragopogon pratensis | **VU D1** | 41, 45, [47] |
| Puccinia iridis | Iris cv. (H) | LC | 44, 47, 49, 52 |
| Puccinia iridis | Iris foetidissima (Part N) | LC | 41-44, 46, 49, 50, 52 |
| Puccinia iridis | Iris foetidissima cv. (H) | LC | 35, 44 |
| Puccinia lagenophorae | Bellis perennis cv. (H) | LC | 44, 46, 47-50, 52 |
| Puccinia lagenophorae | Bellis perennis | LC | 35, 41-52 |
| Puccinia lagenophorae | Senecio jacobaea | LC | 44 |
| Puccinia lagenophorae | Senecio squalidus (N) | LC | 35, 41, 44, 49-52 |
| Puccinia lagenophorae | Senecio vulgaris | LC | 35, 41-52 |
| Puccinia lapsanae | Lapsana communis ssp. communis | LC | 35, 41-52 |
| Puccinia longicornis | Sasa palmata (H) | NE | 41, 44, 46 |
| Puccinia luzulae | Luzula sp. | **VU D** | 47 |
| Puccinia luzulae | Luzula pilosa | **VU D** | 44, [46], 48, 49 |
| Puccinia maculosa | Mycelis muralis | **EN D** | 42-44, [48], [49], 51 |
| Puccinia magnusiana | Phragmites australis | LC | 35, 41, 44, 46-48, 49, 52 |
| Puccinia magnusiana | Ranunculus flammula | LC | 48 |
| Puccinia magnusiana | Ranunculus repens | LC | 35, 44, 46, 49, 52 |
| Puccinia magnusiana | Ranunculus repens ssp. repens | LC | 44 |
| Puccinia major | Crepis paludosa | **RE** | [48] |
| Puccinia malvacearum | Alcea rosea (H) | LC | 35, 42-44, 46, 47, 49, 52 |
| Puccinia malvacearum | Althaea officinalis | LC | 41, 44 |
| Puccinia malvacearum | Malva arborea | LC | 41, 44 |
| Puccinia malvacearum | Malva × clementii (N/H) | LC | 44 |
| Puccinia malvacearum | Malva moschata | LC | 35, 41-44, 46, 49 |
| Puccinia malvacearum | Malva neglecta (A) | LC | 49 |
| Puccinia malvacearum | Malva pusilla (N) | LC | 41 |
| Puccinia malvacearum | Malva sp. | LC | 44, 46, 49 |
| Puccinia malvacearum | Malva sylvestris (A) | LC | 35, 41-52 |
| Puccinia malvacearum | Undetermined | LC | 50, 52 |
| Puccinia menthae | Clinopodium ascendens | LC | 42, 45, 46, 49 |
| Puccinia menthae | Clinopodium sp. | LC | 45, 49 |
| Puccinia menthae | Clinopodium vulgare | LC | 42, 43, 49 |
| Puccinia menthae | Mentha aquatica | LC | 35, 41-50, 52 |
| Puccinia menthae | Mentha arvensis | LC | 44, 49 |
| Puccinia menthae | Mentha cv. (H) | LC | 41, 42, 44, 47 |
| Puccinia menthae | Mentha × dumetorum var. citrata (H) | LC | 44 |
| Puccinia menthae | Mentha × gracilis (H) | LC | 44, 46, 49 |
| Puccinia menthae | Mentha longifolia (N) | LC | 44 |
| Puccinia menthae | Mentha × piperita (N?) | LC | 42, 44, 46, 49 |
| Puccinia menthae | Mentha × piperita var. citrata (H) | LC | 49, 52 |
| Puccinia menthae | Mentha pulegium | LC | 44 |
| Puccinia menthae | Mentha sp. | LC | 41, 42, 44, 49 |
| Puccinia menthae | Mentha × smithiana (N) | LC | 44, 46 |
| Puccinia menthae | Mentha spicata (A) | LC | 41, 42, 44, 46, 49, 52 |
| Puccinia menthae | Mentha suaveolens | LC | 44, 46, 49, 52 |
| Puccinia menthae | Mentha × verticellata | LC | 42, 46, 49, 52 |
| Puccinia menthae | Mentha × villosa (N) | LC | 46, 52 |
| Puccinia menthae | Mentha × villosonervata (N) | LC | 46, 49 |
| Puccinia menthae | Origanum vulgare | LC | 42, 44, 46, 48, 49, 52 |
| Puccinia menthae | Undetermined | LC | 41, 48, 49, 52 |
| Puccinia nemoralis | Melampyrum pratense | **VU D2** | 48, 49 |
| Puccinia nemoralis | Molinia caerulea | **VU D2** | 48, 49 |
| Puccinia nitida | Aethusa cynapium ssp. agrestis | **VU D2** | 35, 46, [49] |
| Puccinia nitida | Aethusa cynapium ssp. cynapium | **VU D2** | 43, 44, 47 |
| Puccinia obscura | Bellis perennis | LC | 48, 49, 52 |
| Puccinia obscura | Luzula campestris | LC | 35, 41-44, 46, 48-50, 52 |
| Puccinia obscura | Luzula forsteri | LC | 43 |
| Puccinia obscura | Luzula multiflora | LC | 42, 46, 49, 52 |
| Puccinia obscura | Luzula pilosa | LC | 43, 48 |
| Puccinia obscura | Luzula sylvatica | LC | 35, 41-44, 46-50, 52 |
| Puccinia opizii | Carex spicata | **NT** | 41 |
| Puccinia opizii | Carex muricata ssp. pairae | **NT** | 42, 43, 46 |
| Puccinia opizii | Carex paniculata | **NT** | 44, 46 |
| Puccinia oxalidis | Oxalis adenophylla (H) | LC | 35 |
| Puccinia oxalidis | Oxalis articulata (N/H) | LC | 35, 44-46 |
| Puccinia oxalidis | Oxalis cv. (H) | LC | 42, 44, 46, 49, 52 |
| Puccinia oxalidis | Oxalis debilis (N/H) | LC | 35, 44, 50 |
| Puccinia oxalidis | Oxalis debilis var. corymbosa (N/H) | LC | 46 |
| Puccinia oxalidis | Oxalis latifolia (N/H) | LC | 35 |
| Puccinia oxalidis | Oxalis rosea (N/H) | LC | 49 |
| Puccinia oxyriae | Oxyria digyna | **CR D1, 2** | 49 |
| Puccinia paludosa | Carex bigelowii | LC | 48 |
| Puccinia paludosa | Carex binervis | LC | 44 |
| Puccinia paludosa | Carex elata | LC | 41, 42, 46, 52 |
| Puccinia paludosa | Carex elata var. aurea | LC | 41, 44, 46 |
| Puccinia paludosa | Carex nigra | LC | 35, 41-46, 49, 52 |
| Puccinia paludosa | Carex panicea | LC | 44, 46, 49 |
| Puccinia pelargonii-zonalis | Pelargonium sp. (H) | NE | 41, 44-46, 49, 50, 52 |
| Puccinia phragmitis | Phragmites australis | LC | 41, 44, 45, 48, 49, 52 |
| Puccinia phragmitis | Phragmites australis cv. | LC | 50 |
| Puccinia phragmitis | Rumex acetosa | LC | 49, 52 |
| Puccinia phragmitis | Rumex acetosella | LC | 52 |
| Puccinia phragmitis | Rumex conglomeratus | LC | 49, 52 |
| Puccinia phragmitis | Rumex crispus | LC | 44, 49, 52 |
| Puccinia phragmitis | Rumex crispus ssp. crispus | LC | 44 |
| Puccinia phragmitis | Rumex hydrolapathum | LC | 46 |
| Puccinia phragmitis | Rumex obtusifolius | LC | 44, 48, 49, 51, 52 |
| Puccinia phragmitis | Rumex sanguineus | LC | 44, 49, 52 |
| Puccinia phragmitis | Rumex sp. | LC | 45, 52 |
| *Puccinia phragmitis* | Undetermined | LC | 47, 52 |
| *Puccinia pimpinellae* | *Pimpinella saxifraga* | **CR D** | [41], 42, 43, [46], 49 |
| *Puccinia poarum* | *Poa annua* | LC | 44, 46, 49, 52 |
| *Puccinia poarum* | *Poa chaixii* (N) | LC | 43 |
| *Puccinia poarum* | *Poa compressa* | LC | 44 |
| *Puccinia poarum* | *Poa humilis* | LC | 35, 41, 44, 46 |
| *Puccinia poarum* | *Poa nemoralis* | LC | 44 |
| *Puccinia poarum* | *Poa trivialis* | LC | 44 |
| *Puccinia poarum* | *Tussilago farfara* | LC | 35, 41-47, 49-52 |
| *Puccinia poarum* | Undetermined | LC | 41, 49, 51 |
| *Puccinia polygoni-amphibii* | *Persicaria amphibia* | LC | 35, 41, 42, 44, 46, 48, 49, 52 |
| *Puccinia polygoni-amphibii* | *Persicaria maculosa* | LC | 35, 44 |
| *Puccinia polygoni-amphibii* var. *convolvuli* | *Fallopia convolvulus* (A) | **VU D** | [41], 42, 44, 49 |
| *Puccinia porri* | *Allium porrum* (H) | NE | 35, 42-46, 49, 52 |
| *Puccinia primulae* | *Primula vulgaris* | **NT** | 44-46, [48], 49, 52 |
| *Puccinia primulae* | *Primula veris* | **NT** | 52 |
| *Puccinia pulverulenta* | *Epilobium adenocaulon* | LC | 44 |
| *Puccinia pulverulenta* | *Epilobium ciliatum* (N) | LC | 43, 44, 46 |
| *Puccinia pulverulenta* | *Epilobium hirsutum* | LC | 35, 41-46, 49, 51-52 |
| *Puccinia pulverulenta* | *Epilobium montanum* | LC | 35, 42-46, 49, 52 |
| *Puccinia pulverulenta* | *Epilobium obscurum* | LC | 44 |
| *Puccinia pulverulenta* | *Epilobium parviflorum* | LC | 44, 52 |
| *Puccinia pulverulenta* | *Epilobium* sp. | LC | 41, 44, 46, 47, 49, 52 |
| *Puccinia pulverulenta* | *Epilobium tetragonum* | LC | 48 |
| *Puccinia punctata* | *Cruciata laevipes* | LC | 41, 42, 43, 44, 47, 51 |
| *Puccinia punctata* | *Galium aparine* | LC | 35, 44, 46 |
| *Puccinia punctata* | *Galium album* | LC | 44, 46 |
| *Puccinia punctata* | *Galium odoratum* | LC | 44, 48 |
| *Puccinia punctata* | *Galium palustre* | LC | 35, 41-44, 46, 48, 49, 52 |
| *Puccinia punctata* | *Galium palustre* ssp. *elongatum* | LC | 42, 44, 46 |
| *Puccinia punctata* | *Galium palustre* ssp. *palustre* | LC | 41, 44, 47 |
| *Puccinia punctata* | *Galium saxatile* | LC | 35, 42, 46, 49, 50, 52 |
| *Puccinia punctata* | *Galium sternerii* | LC | 42 |
| *Puccinia punctata* | *Galium uliginosum* | LC | 42, 49 |
| *Puccinia punctata* | *Galium verum* | LC | 35, 41-44, 46, 48, 49, 51, 52 |
| *Puccinia punctata* | Undetermined | LC | 47 |
| *Puccinia punctiformis* | *Cirsium arvense* | LC | 35, 41-52 |
| *Puccinia pygmaea* var. *pygmaea* | *Calamagrostis epigejos* | **CR D** | 46, [52] |
| *Puccinia pygmaea* var. *ammophilina* | *Ammophila arenaria* | LC | 41, 44-46, 48, 49, 51, 52 |
| *Puccinia recondita* | *Agrostis capillaris* | LC | 42, 44, 46 |
| *Puccinia recondita* | *Agrostis stolonifera* | LC | 44 |
| *Puccinia recondita* | *Alopecurus geniculatus* | LC | 44 |
| *Puccinia recondita* | *Alopecurus myosuroides* | LC | 44 |
| *Puccinia recondita* | *Alopecurus pratensis* | LC | 44, 46 |
| *Puccinia recondita* | *Anisantha sterilis* (A) | LC | 41, 44 |
| *Puccinia recondita* | *Bromopsis erecta* | LC | 41 |
| Puccinia recondita | Bromopsis ramosa | LC | 42-44, 49, 51 |
| Puccinia recondita | Bromus commutatus | LC | 35 |
| Puccinia recondita | Bromus hordeaceus ssp. hordeaceus | LC | 35, 41, 44, 47, 49 |
| Puccinia recondita | Elymus caninus | LC | 44 |
| Puccinia recondita | Elytrigia atherica | LC | 41, 44, 49 |
| Puccinia recondita | Elytrigia atherica var. setigera | LC | 41, 44 |
| Puccinia recondita | Elytrigia juncea ssp. boreoatlantica | LC | 41, 44-46, 48, 49, 52 |
| Puccinia recondita | Elytrigia repens ssp. repens | LC | 35, 41-46, 48-50, 52 |
| Puccinia recondita | Elytrigia repens ssp. repens var. aristata | LC | 41 |
| Puccinia recondita | Elytrigia × obtusiuscula | LC | 44 |
| Puccinia recondita | Hordeum murinum (A) | LC | 44 |
| Puccinia recondita | Secale cereale (Cas) | LC | 44, 49 |
| Puccinia recondita | Thalictrum flavum | LC | 41, 46 |
| Puccinia saniculae | Sanicula europaea | LC | 42, 44-49, 51, 52 |
| Puccinia saxifragae | Saxifraga granulata | **EN D** | 43 |
| Puccinia saxifragae | Saxifraga stellaris | **EN D** | [46], 48, 49 |
| Puccinia schroeteriana | Carex hostiana | **CR D** | 52 |
| Puccinia schroeteriana | Serratula tinctoria | **CR D** | 52 |
| Puccinia scirpi | Nymphoides peltata/Schoenoplectus lacustris | **DD** | 42 |
| Puccinia scorzonerae | Scorzonera humilis | **CR D** | 41 |
| Puccinia sessilis | Allium ursinum | LC | 35, 42-52 |
| Puccinia sessilis | Arum creticum (H) | LC | 52 |
| Puccinia sessilis | Arum italicum ssp. neglectum (N/H) | LC | 44 |
| Puccinia sessilis | Arum maculatum | LC | 35, 41-47, 49-52 |
| Puccinia sessilis | Dactylorhiza fuchsii | LC | 42, 52 |
| Puccinia sessilis | Dactylorhiza incarnata | LC | 52 |
| Puccinia sessilis | Dactylorhiza maculata | LC | 52 |
| Puccinia sessilis | Dactylorhiza praetermissa | LC | 42 |
| Puccinia sessilis | Dactylorhiza purpurella | LC | 52 |
| Puccinia sessilis | Gymnadenia conopsea | LC | 52 |
| Puccinia sessilis | Phalaris arundinacea | LC | 35, 41-46, 48-49, 51, 52 |
| Puccinia smyrnii | Smyrnum olusatrum (A) | LC | 35, 41, 42, 44-46, 48-52 |
| Puccinia striiformis var. dactylidis | Dactylis glomerata | LC | 35, 41, 42, 44-49 |
| Puccinia striiformis | Brachypodium sylvaticum | LC | 49, 52 |
| Puccinia striiformis | Elymus caninus | LC | 44 |
| Puccinia striiformis | Elymus magellanicus (H) | LC | 44 |
| Puccinia striiformis | Elytrigia repens ssp. repens | LC | 41 |
| Puccinia striiformis | Holcus lanatus | LC | 44 |
| Puccinia striiformis | Hordeum murinum (A) | LC | 41, 44 |
| Puccinia striiformis | Hordeum murinum ssp. murinum (A) | LC | 44 |
| Puccinia striiformis | Hordeum secalinum | LC | 41, 44 |
| Puccinia striiformis | Hordeum sp. | LC | 46 |
| Puccinia striiformis | Triticum aestivum 'Red Marvel' (H) | LC | 41 |
| Puccinia striiformis | Triticum turgidum 'Rivet Wheat' (H) | LC | 41 |
| Puccinia striiformis | Triticum sp. (H) | LC | 46 |
| Puccinia tanacetii | Artemisia absinthium (A) | LC | 44, 45, 49, 50, 52 |
| *Puccinia tanacetii* | *Artemisia arborescens* 'Powis Castle' (H) | LC | 44 |
| *Puccinia tanacetii* | *Artemisia vulgaris* (A) | LC | 41, 44, 46 |
| *Puccinia tanacetii* | *Seriphidium maritimum* | LC | 41, 44 |
| *Puccinia thymi* | *Thymus polytrichus* | **EN D** | 42, 45 |
| *Puccinia tumida* | *Conopodium majus* | LC | 35, 41-49 |
| *Puccinia uliginosa* | *Carex nigra* | **RE** | 52 |
| *Puccinia umbilici* | *Umbilicus rupestris* | LC | 35, 41-46, 48-50, 52 |
| *Puccinia urticata* | *Urtica dioica* | LC | 35, 41-50, 52 |
| *Puccinia urticata* var. *biporula* | *Carex pallescens* | **EN D** | 41, 42, 46 |
| *Puccinia urticata* var. *urticae-acutae* | *Carex nigra* | **DD** | 42, 44, 52 |
| *Puccinia urticata* var. *urticae-acutiformis* | *Carex acutiformis* | LC | 35, 42, 43, 46, 48, 49, 51, 52 |
| *Puccinia urticata* var. *urticae-acutiformis* | *Carex pseudocyperus* | LC | 44, 49 |
| *Puccinia urticata* var. *urticae-flacca* | *Carex flacca* | LC | 42, 44-46, 49, 50, 52 |
| *Puccinia urticata* var. *urticae-hirtae* | *Carex hirta* | LC | 35, 41-44, 46, 49, 52 |
| *Puccinia urticata* var. *urticae-inflatae* | *Carex rostrata* | **NT** | 43-47, 52 |
| *Puccinia urticata* var. *urticae-ripariae* | *Carex riparia* | **VU D1, 2** | 44, 46 |
| *Puccinia urticata* var. *urticae-vesicariae* | *Carex vesicaria* | LC | 42-44, 46, 49 |
| *Puccinia variabilis* | *Taraxacum officinale* agg. | LC | 41-44, 46-52 |
| *Puccinia variabilis* | *Taraxacum* sp. | LC | 42, 49-51 |
| *Puccinia veronicae* | *Veronica montana* | LC | 35, 41-52 |
| *Puccinia veronicae-longifoliae* | *Veronica spicata* | **RE** | [50] |
| *Puccinia vincae* | *Vinca major* (N/H) | LC | 35, 41, 43-47, 49, 50, 52 |
| *Puccinia vincae* | *Vinca* sp. (N/H) | LC | 47 |
| *Puccinia violae* | *Viola canina* | LC | 41, 46, 48, 49, 52 |
| *Puccinia violae* | *Viola canina* ssp. *canina* | LC | 44, 46, 48 |
| *Puccinia violae* | *Viola* cv. (H) | LC | 43, 44 |
| *Puccinia violae* | *Viola hirta* | LC | 44, 49 |
| *Puccinia violae* | *Viola odorata* | LC | 43, 52 |
| *Puccinia violae* | *Viola reichenbachiana* | LC | 35, 44, 52 |
| *Puccinia violae* | *Viola riviniana* | LC | 35, 41-44, 46-52 |
| *Puccinia violae* | *Viola* sp. | LC | 35, 41-43, 47-51 |
| *Puccinia violae* | *Viola tricolor* | LC | 41, 52 |
| *Puccinia virgae-aureae* | *Solidago virgaurea* | **CR D** | 44, 49 |
| *Pucciniastrum agrimonae* | *Agrimonia eupatoria* | **NT** | 41, [47], 48, 49, [50], 51 |
| *Pucciniastrum agrimonae* | *Agrimonia* sp. | **NT** | 35, 47-50 |
| *Pucciniastrum circaeae* | *Circaea* × *intermedia* | LC | 35, 41-43, 46 |
| *Pucciniastrum circaeae* | *Circaea lutetiana* | LC | 35, 41-46, 48-52 |
| *Pucciniastrum circaeae* | *Circaea* sp. | LC | 41, [47], 49, 50 |
| *Pucciniastrum epilobii* | *Chamerion angustifolium* | LC | 35, 41-44, 46-52 |
| *Pucciniastrum epilobii* | *Clarkia amoena* (H) | LC | 41, 44, 49 |
| Pucciniastrum epilobii | Epilobium alsinifolium | LC | 49 |
| Pucciniastrum epilobii | Epilobium ciliatum (N) | LC | 35, 42-44, 46-49 |
| Pucciniastrum epilobii | Epilobium ciliatum × montanum | LC | 47 |
| Pucciniastrum epilobii | Epilobium ciliatum/obscurum | LC | 46 |
| Pucciniastrum epilobii | Epilobium hirsutum | LC | 35, 41, 42, 44, 46, 49, 52 |
| Pucciniastrum epilobii | Epilobium lanceolatum | LC | 44 |
| Pucciniastrum epilobii | Epilobium montanum | LC | 41, 44, 46-50, 52 |
| Pucciniastrum epilobi | Epilobium obscurum | LC | 46, 49, 52 |
| Pucciniastrum epilobii | Epilobium palustre | LC | 42, 44-46, 49 |
| Pucciniastrum epilobii | Epilobium parviflorum | LC | 42, 44, 52 |
| Pucciniastrum epilobii | Epilobium sp | LC | 44, 46, 49 |
| Pucciniastrum epilobii | Fuchsia cv. (H) | LC | 44, 47, 49, 52 |
| Pucciniastrum epilobii | Undetermined | LC | 42, 44, 46, 49, 51 |
| Pucciniastrum guttatum | Galium odoratum | LC | 44 |
| Pucciniastrum guttatum | Galium palustre | LC | 44, 46 |
| Pucciniastrum guttatum | Galium saxatile | LC | 35, 42, 44, 46, 49, 52 |
| Pucciniastrum guttatum | Galium uliginosum | LC | 42 |
| Pucciniastrum guttatum | Galium verum | LC | 44, 46, 48, 49, 52 |
| Pucciniastrum guttatum | Sherardia arvensis | LC | 49, 52 |
| Trachyspora intrusa | Alchemilla glabra | **NT** | 42, 44, 49 |
| Trachyspora intrusa | Alchemilla sp. | **NT** | 41-44, 47, 49 |
| Trachyspora intrusa | Alchemilla vulgaris | **NT** | 48, 49 |
| Trachyspora intrusa | Alchemilla xanthochlora | **NT** | 35, 41, 44, 49 |
| Tranzschelia anemones | Anemone nemorosa | LC | 35, 42, 44-47, [48], 49, 50, 52 |
| Tranzschelia anemones | Thalictrum minus | LC | 42 |
| Tranzschelia anemones | Undetermined | LC | 48 |
| Tranzschelia discolor | Prunus domestica (A/H) | LC | 35, 42-44, 46, 47, 49, 50 |
| Tranzschelia discolor | Prunus domestica ssp. domestica (A/H) | LC | 42, 46, 49 |
| Tranzschelia discolor | Prunus domestica ssp. insititia (A/H) | LC | 46 |
| Tranzschelia discolor | Prunus sp. | LC | 46 |
| Tranzschelia pruni-spinosae | Prunus domestica (A/H) | LC | 44 |
| Tranzschelia pruni-spinosae | Prunus spinosa | LC | 35, 41-46, 49, 52 |
| Tranzschelia pruni-spinosae | Prunus sp. | LC | [47] |
| Triphragmium filipendulae | Filipendula vulgaris | **VU B, D1** | 49 |
| Triphragmium ulmariae | Filipendula ulmaria | LC | 35, 41-52 |
| Uredo morvernensis | Thymus polytrichus | **CR D** | 44 |
| Uromyces acetosae | Rumex acetosa | LC | 41-49 |
| Uromyces acetosae | Rumex acetosa ssp. acetosa | LC | 42, 43, 46, 49 |
| Uromyces acetosae | Rumex acetosella | LC | 49, 52 |
| Uromyces airae-flexuosae | Deschampsia flexuosa | LC | 35, 41-44, 46-49, 52 |
| Uromyces ambiguus | Allium scorodoprasum (Cas) | NE | 46 |
| Uromyces ambiguus | Allium schoenoprasum (H) | NE | 46 |
| Uromyces anthyllidis | Anthyllis vulneraria | LC | 41, 44-46, 48-50, 52 |
| Uromyces anthyllidis | Anthyllis vulneraria ssp. vulneraria | LC | 41, 44 |
| Uromyces appendiculatus | Phaseolus coccineus (H) | NE | 35, 41-46 |
| Uromyces armeriae | Armeria maritima ssp. maritima | LC | 41, 44-46, 49, 51, 52 |
| Uromyces behenis | Silene uniflora | LC | 45, 46, 49, 52 |
| Uromyces behenis | Silene vulgaris | LC | 52 |
| Uromyces beticola | Beta vulgaris ssp. vulgaris (H) | LC | 45, 49 |
| Uromyces beticola | Beta vulgaris ssp. cicla (H) | LC | 45 |
| Uromyces beticola | Beta vulgaris ssp. maritima | LC | 35, 41, 44, 45, 49, 51, 52 |
| Uromyces beticola | Spinacia oleracea (H) | LC | 41 |
| Uromyces chenopodii | Suaeda maritima | LC | 41, 44-46, 48, 49, 51, 52 |
| Uromyces dactylidis | Dactylis glomerata | LC | 35, 42, 44, 45, 49, 52 |
| Uromyces dactylidis | Poa annua | LC | 44 |
| Uromyces dactylidis | Poa humilis | LC | 44 |
| Uromyces dactylidis | Poa nemoralis | LC | 44, 45 |
| Uromyces dactylidis | Poa pratensis | LC | 44 |
| Uromyces dactylidis | Poa sp. | LC | 35, 42, 48-50, 52 |
| Uromyces dactylidis | Poa trivialis | LC | 35, 42-45, 47 |
| Uromyces dactylidis | Ranunculus acris var. acris | LC | 44, 45 |
| Uromyces dactylidis | Ficaria verna | LC | 35, 41-52 |
| Uromyces dactylidis | Ficaria verna ssp. fertilis | LC | 35, 41-44, 46, 49, 51, 52 |
| Uromyces dactylidis | Ficaria verna ssp. verna | LC | 43-46 |
| Uromyces dactylidis | Ranunculus repens | LC | 35, 41-49, 52 |
| Uromyces dactylidis | Undetermined | LC | 35, 50 |
| Uromyces dianthi | Dianthus barbatus (H) | NE | 44 |
| Uromyces dianthi | Dianthus caryophyllus (H) | NE | 44 |
| Uromyces ervi | Vicia hirsuta | **NT** | 35, 42-44, 46-49, 52 |
| Uromyces ervi | Vicia tetrasperma | **NT** | 46 |
| Uromyces fallens | Trifolium medium | LC | 35, |
| Uromyces fallens | Trifolium sp. | LC | 51, 52 |
| Uromyces fallens | Trifolium pratense | LC | 35, 41-44, 46, 49, 52 |
| Uromyces ficariae | Ficaria verna | LC | 35, 42-50, 52 |
| Uromyces ficariae | Ficaria verna ssp. fertilis | LC | 43, 44 |
| Uromyces ficariae | Ficaria verna ssp. verna | LC | 35, 41-47, 49, 52 |
| Uromyces geranii | Geranium dissectum | LC | 52 |
| Uromyces geranii | Geranium molle | LC | 41, 49, 52 |
| Uromyces geranii | Geranium phaeum (N) | LC | 44 |
| Uromyces geranii | Geranium pratense | LC | 44, 46, 49, 50, 52 |
| Uromyces geranii | Geranium pyrenaicum (N) | LC | 35, 42, 44, 47-49, 51, 52 |
| Uromyces geranii | Geranium robertianum | LC | 35, 41, 42, 44, 46, 48, 49, 52 |
| Uromyces geranii | Geranium robertianum ssp. celticum | LC | 41, 44 |
| Uromyces inaequaltus | Silene nutans | **RE** | [51] |
| Uromyces junci | Juncus articulatus | LC | 41, 44, 46, 52 |
| Uromyces junci | Juncus bufonius | LC | 44 |
| Uromyces junci | Juncus subnodulosus | LC | 41, 44 |
| Uromyces junci | Pulicaria dysenterica | LC | 44, 45, 52 |
| Uromyces limonii | Limonium vulgare | LC | 41, 44, 48, 49, 52 |
| Uromyces lineolatus | Bolboschoenus maritimus | LC | 41, 44-46, 49, 52 |
| Uromyces lineolatus | Glaux maritima | LC | 44, 45, 52 |
| Uromyces lineolatus | Oenanthe crocata | LC | 41, 44, 45, 49, 52 |
| Uromyces lineolatus | Oenanthe lachenalii | LC | 44, 49, 52 |
| Uromyces minor | Trifolium campestre | LC | 44 |
| Uromyces minor | Trifolium dubium | LC | 35, 41-46, 49, 52 |
| Uromyces muscari | Hyacinthoides hispanica (N/H) | LC | 35, 44, 46, 49, 52 |
| Uromyces muscari | Hyacinthoides non-scripta | LC | 35, 41-52 |
| Uromyces muscari | Hyacinthoides non-scripta × hispanica (N) | LC | 41, 44, 46, 49, 50, 52 |
| Uromyces muscari | Scilla verna | LC | 35, 41, 45, 46, 49, 52 |
| Uromyces muscari | Undetermined | LC | 44, 49 |
| Uromyces pisi-sativi | Cytisus scoparius | LC | 44 |
| Uromyces pisi-sativi | Genista anglica | LC | 35 |
| Uromyces pisi-sativi | Genista pilosa | LC | 42 |
| Uromyces pisi-sativi | Lathyrus latifolius | LC | 44 |
| Uromyces pisi-sativi | Lathyrus odoratus cv. (H) | LC | 35 |
| Uromyces pisi-sativi | Lathyrus pratensis | LC | 41-44, 46, 48, 49, 52 |
| Uromyces pisi-sativi | Lathyrus sylvestris | LC | 44 |
| Uromyces pisi-sativi | Lotus corniculatus | LC | 35, 41-44, 46, 48, 49, 52 |
| Uromyces pisi-sativi | Lotus corniculatus var. sativus (N) | LC | 44 |
| Uromyces pisi-sativi | Lotus glaber | LC | 44 |
| Uromyces pisi-sativi | Lotus pedunculatus | LC | 41, 44 |
| Uromyces pisi-sativi | Medicago arabica | LC | 49 |
| Uromyces pisi-sativi | Medicago lupulina | LC | 35, 41, 44, 45, 49, 52 |
| Uromyces pisi-sativi | Onobrychis viciifolia (N) | LC | 44 |
| Uromyces pisi-sativi | Ulex europaeus | LC | 46 |
| Uromyces pisi-sativi | Ulex galli | LC | 46 |
| Uromyces polygoni-avicularis | Polygonum arenastrum (A) | LC | 41, 44, 46 |
| Uromyces polygoni-avicularis | Polygonum aviculare | LC | 35, 41-46, 49, 50, 52 |
| Uromyces polygoni-avicularis | Polygonum polychnemiforme | LC | 46 |
| Uromyces polygoni-avicularis | Polygonum rurivagum (A) | LC | 44 |
| Uromyces polygoni-avicularis | Undetermined | LC | 47 |
| Uromyces rumicis | Rumex acetosa ssp. acetosa | LC | 42, 44 |
| Uromyces rumicis | Rumex conglomeratus | LC | 44 |
| Uromyces rumicis | Rumex crispus | LC | 35, 41, 44-46, 52 |
| Uromyces rumicis | Rumex crispus ssp. crispus | LC | 44 |
| Uromyces rumicis | Rumex crispus ssp. littoreus | LC | 46 |
| Uromyces rumicis | Rumex hydrolapathum | LC | 41, 46 |
| Uromyces rumicis | Rumex obtusifolius | LC | 44, 45 |
| Uromyces rumicis | Rumex sanguineus | LC | 35 |
| Uromyces rumicis | Rumex sanguineus var. viridis | LC | 44 |
| Uromyces rumicis | Rumex sp. | LC | 47-49, 52 |
| Uromyces salicorniae | Salicornia dolichostachya | LC | 44, 46, 49, 52 |
| Uromyces salicorniae | Salicornia europaea | LC | 44, 49, 52 |
| Uromyces salicorniae | Salicornia pusilla | LC | 41, 44 |
| Uromyces salicorniae | Salicornia pusilla × ramosissima | LC | 41 |
| Uromyces salicorniae | Salicornia ramosissima | LC | 41, 44, 46, 49 |
| Uromyces salicorniae | Salicornia sp. | LC | 49, 52 |
| Uromyces scrophulariae | Scrophularia auriculata | **EN D** | 44, [47] |
| Uromyces silphii | Juncus tenuis (N) | LC | 41, 44, 46, 49 |
| *Uromyces sommerfeltii* | *Solidago virgaurea* | **EN D** | 46, 49 |
| *Uromyces sparsus* | *Spergularia marina* | **NT** | 41, 44, 49, 52 |
| *Uromyces sparsus* | *Spergularia media* | **NT** | 46, 52 |
| *Uromyces trifolii* | *Trifolium fragiferum* | LC | 44 |
| *Uromyces trifolii* | *Trifolium hybridum* (N) | LC | 44 |
| *Uromyces trifolii* | *Trifolium pratense* | LC | 49 |
| *Uromyces trifolii* | *Trifolium repens* | LC | 35, 41-44, 46, 49, 50 |
| *Uromyces trifolii* | *Trifolium* sp. | LC | 52 |
| *Uromyces trifolii-repentis* | *Trifolium hybridum* (N) | LC | 35, 44 |
| *Uromyces trifolii-repentis* | *Trifolium repens* | LC | 44 |
| *Uromyces valerianae* | *Valeriana officinalis* | LC | 35, 41-49, 52 |
| *Uromyces valerianae* | *Valeriana officinalis* ssp. *sambucifolia* | LC | 46 |
| *Uromyces viciae-fabae* | *Lathyrus pratensis* | LC | 49, 50, 52 |
| *Uromyces viciae-fabae* | *Vicia cracca* | LC | 35, 41-44, 46, 49, 50, 52 |
| *Uromyces viciae-fabae* | *Vicia hirsuta* | LC | 41, 44, 49, 52 |
| *Uromyces viciae-fabae* | *Vicia lathyroides* | LC | 48 |
| *Uromyces viciae-fabae* | *Vicia sativa* | LC | 41, 44, 46, 48-50, 52 |
| *Uromyces viciae-fabae* | *Vicia sativa* ssp. *segetalis* (A) | LC | 44, 46 |
| *Uromyces viciae-fabae* | *Vicia sepium* | LC | 35, 41-44, 46, 49, 51, 52 |
| *Uromyces viciae-fabae* | *Pisum sativum* (H) | LC | 49 |
| *Uromyces viciae-fabae* | Undetermined | LC | 47 |
| *Uromyces viciae-fabae* | *Vicia faba* (H) | LC | 41-44, 46, 48, 49 |
| *Uromyces viciae-fabae*<br>var. *orobi* | *Lathyrus linifolius* var. *montanus* | **VU D** | 43, 47, 48 |
| *Xenodochus carbonarius* | *Sanguisorba officinalis* | LC | 42, 43, 44, 47 |

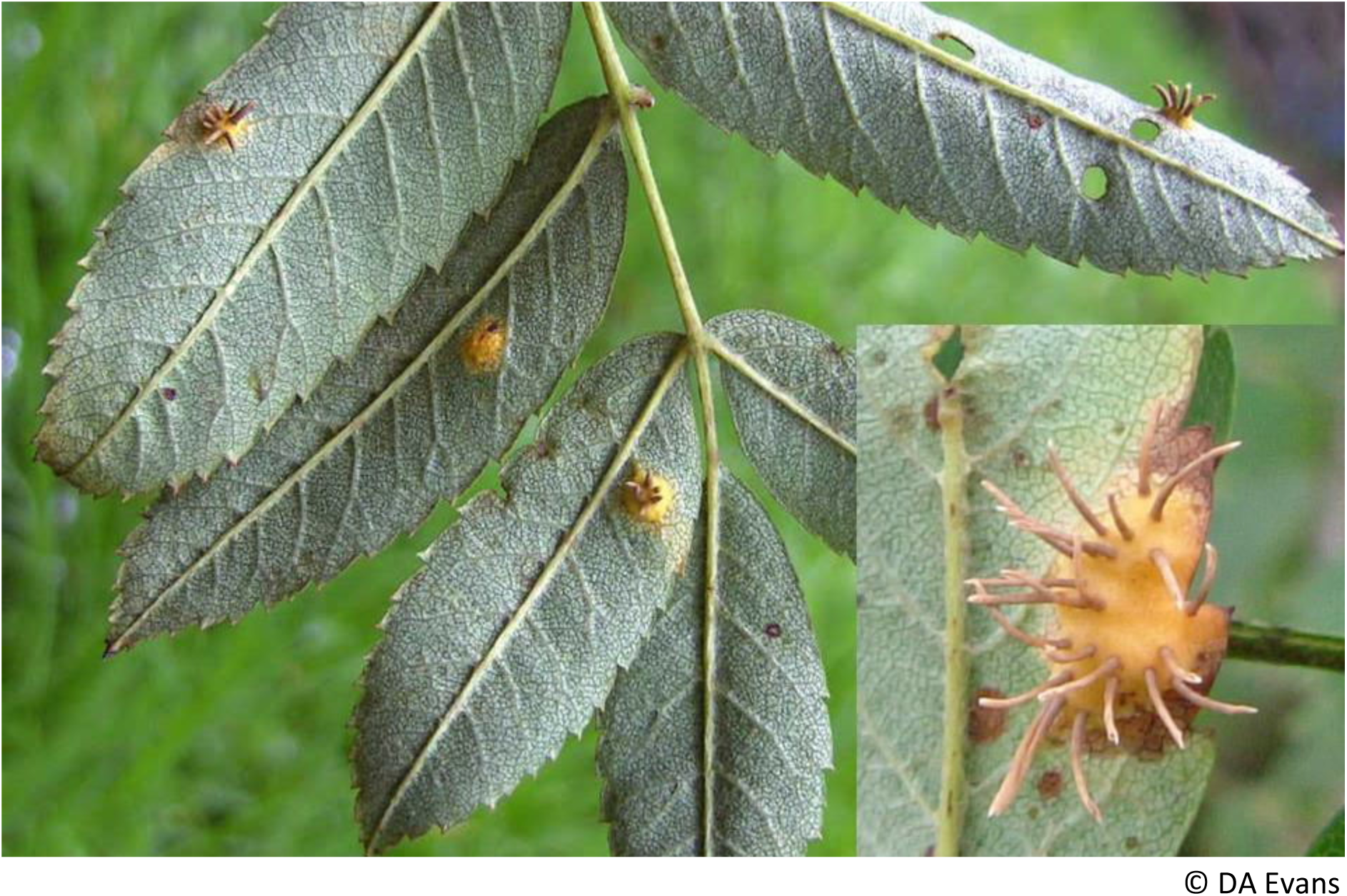
*Gymnosporangium cornutum* on *Sorbus aucuparia*

## Acknowledgements

The authors acknowledge with grateful thanks the generosity of numerous recorders who were willing to share their records and answer queries. The following referees helped with difficult taxa:- Herbert Boyle, Halvor Gjaerum, Sarah Hambleton, Jarkka Hantula, Douglas Henderson, Ming Pei, Tom Preece, Charlotte Swartz, Jan Zadoks and Peter Zwetko.

During the collection of these records numerous herbarium packets have been created. Some material has been deposited in the fungaria of the National Museum of Wales, Cardiff and in the Royal Botanic Gardens at Kew and Edinburgh, though the majority of the material remains in the hands of the recorders and authors. We are also indebted to Paul Smith for his help in preparing the manuscript for press, for permission to use some of his images and in his wise counsel in helping to draft this account and to Eilir Evans for the Welsh translations.

## Appendix 1

### Vice-County Distribution of Rusts on Cultivars of Plants

The following table lists the rust taxa recorded on cultivars of plants. The coverage across Wales is uneven and the list far from complete. Records were sufficiently numerous from Carmarthenshire, Cardiganshire, Caernarvonshire and Anglesey to consider this list worth publishing. It provides a baseline against which changes in distribution of these rusts might be measured.

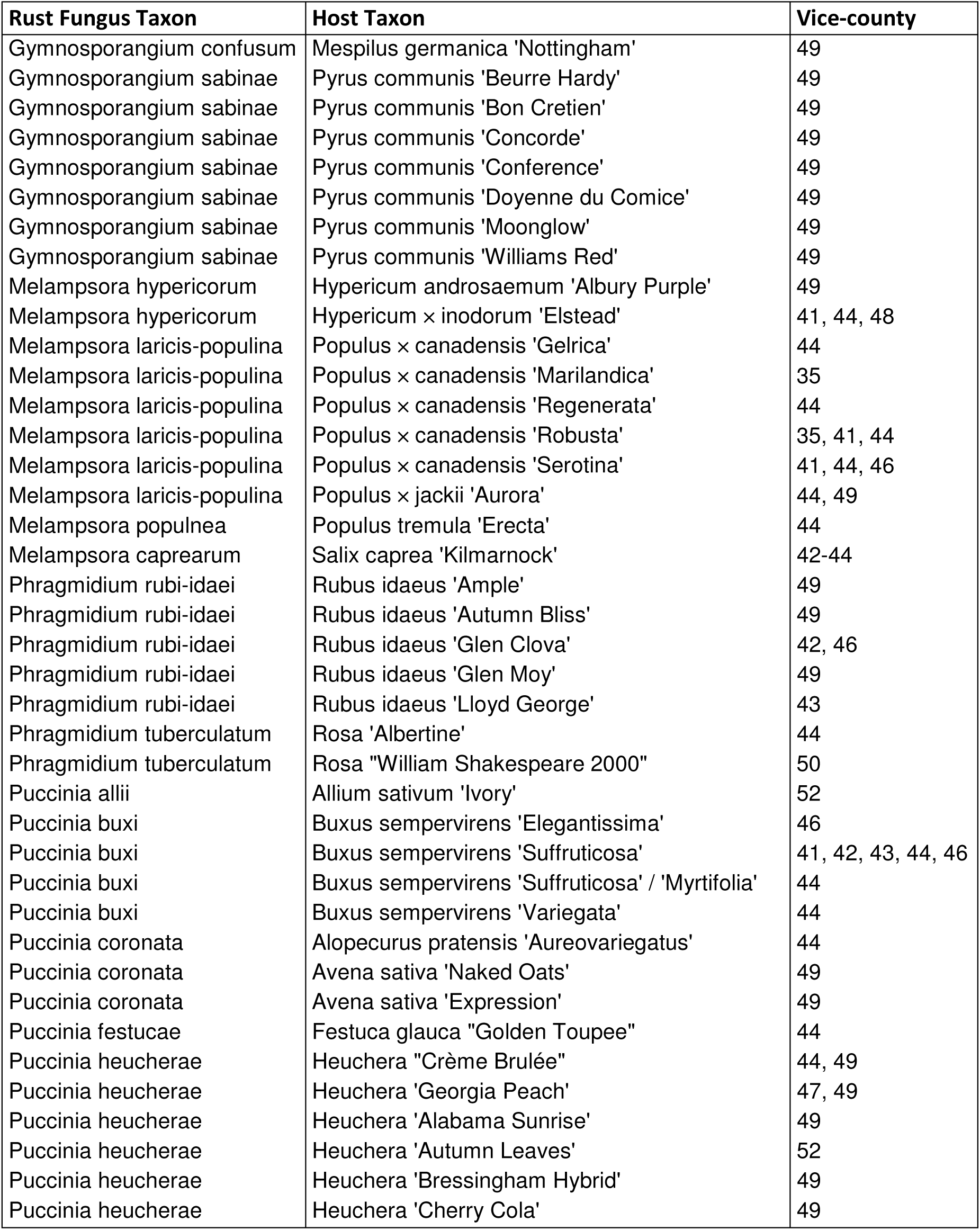

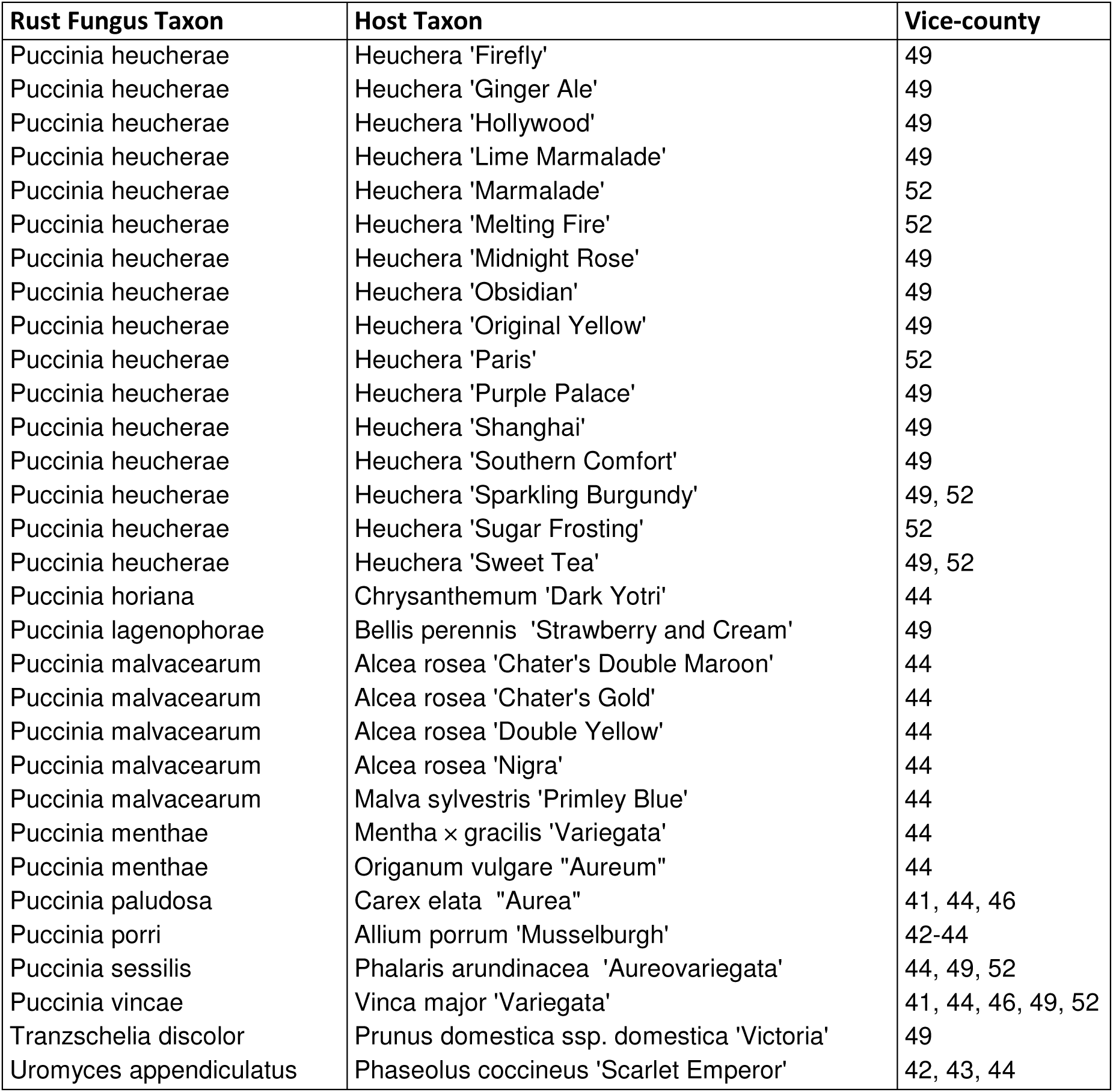

## Appendix 2

### Rust Taxa on Native Hosts Recorded from Elsewhere in Britain but not yet Confirmed from Wales

The following taxa may occur in Wales but have yet to be reported from the principality. They are listed by the habitat they may occur in.

#### Sub-montane, upland and sea cliff

*Hyalopsora adianti-capilli-veneris* On maidenhair fern *Adiantum capilli-veneris*, a rare fern confined to coastal districts in Wales. This rust has been rarely reported, though has been found in Scotland, Kent and County Galway, Ireland.

*Hyalopsora aspidiotus* Found on oak fern *Gymnocarpium dryopteris*, there are no recent British records of the rust fungus but the fern is widespread in Wales.

*Phragmidium acuminatum* The FRDBI lists a single record from Morayshire, Scotland in 1879 on stone bramble *Rubus saxatilis*. This species does, however, appear on a list of rusts from Forden Parish in Montgomeryshire (Vize 1882) but there are no records of the host from that part of Montgomeryshire and the record appears nowhere else in the literature. The rust is widespread in Europe with records from, amongst other countries, Norway, Denmark and Germany, and might just reach Wales.

*Puccinia pazschkei* var*. juliana* An uncommon rust in Scotland on purple saxifrage *Saxifraga oppositifolia* and yellow saxifrage *S. aizoides*. The former host is not uncommon on cliffs in Snowdonia and the Brecon Beacons. Other sub-montane rust species found in Wales have a markedly disjunct distribution with the nearest populations occurring in Highland Scotland, so this rust might occur in Wales.

*Puccinia septentrionalis* Occurs on alpine rue *Thalictrum alpinum* and viviparous bistort *Persicaria vivipara* in several locations in Scotland. The tiny populations of these hosts in Snowdonia should be searched for this rust.

*Pucciniastrum pyrolae* Reported from serrated wintergreen *Orthilia secunda* in the north of England and Scotland. The host in the Brecon Beacons should be searched for this rust.

*Uredinopsis filicina* This rust has been reported from Scotland and north and south-east England on beech fern *Phegopteris connectilis*, a widespread fern in the uplands of Wales.

#### Woodland

*Pucciniastrum areolatum* Alternating between Norway spruce *Picea abies* and bird cherry *Prunus padus*, there are a number of records of this rust from the north of England and Scotland. Given that both hosts are frequent in mid and north Wales this rust may yet be found in Wales.

#### Grassland and Wetland

*Puccinia clintonii* Rarely reported on marsh lousewort *Pedicularis palustris* and lousewort *P. sylvatica*. The FRDBI notes a record from Arklow, County Wicklow, Ireland and from the north of Scotland. It just may occur in Wales.

*Puccinia longissima* Occurring on crested hair-grass *Koeleria macrantha* as near to Wales as east Gloucestershire, this rust should be sought in South Wales.

*Puccinia moliniae* Alternating between purple moor-grass *Molinia caerulea* and selfheal *Prunella vulgaris,* there is a recent record on this latter host on the FRDBI from east Gloucestershire and old records from the former host from north Wiltshire and north Somerset. There are numerous records from Scotland and occasional records from the north of England. Given the frequency of the host plants in Wales there is a distinct probability of this rust occurring in Wales.

*Puccinia moliniae* on purple moor-grass

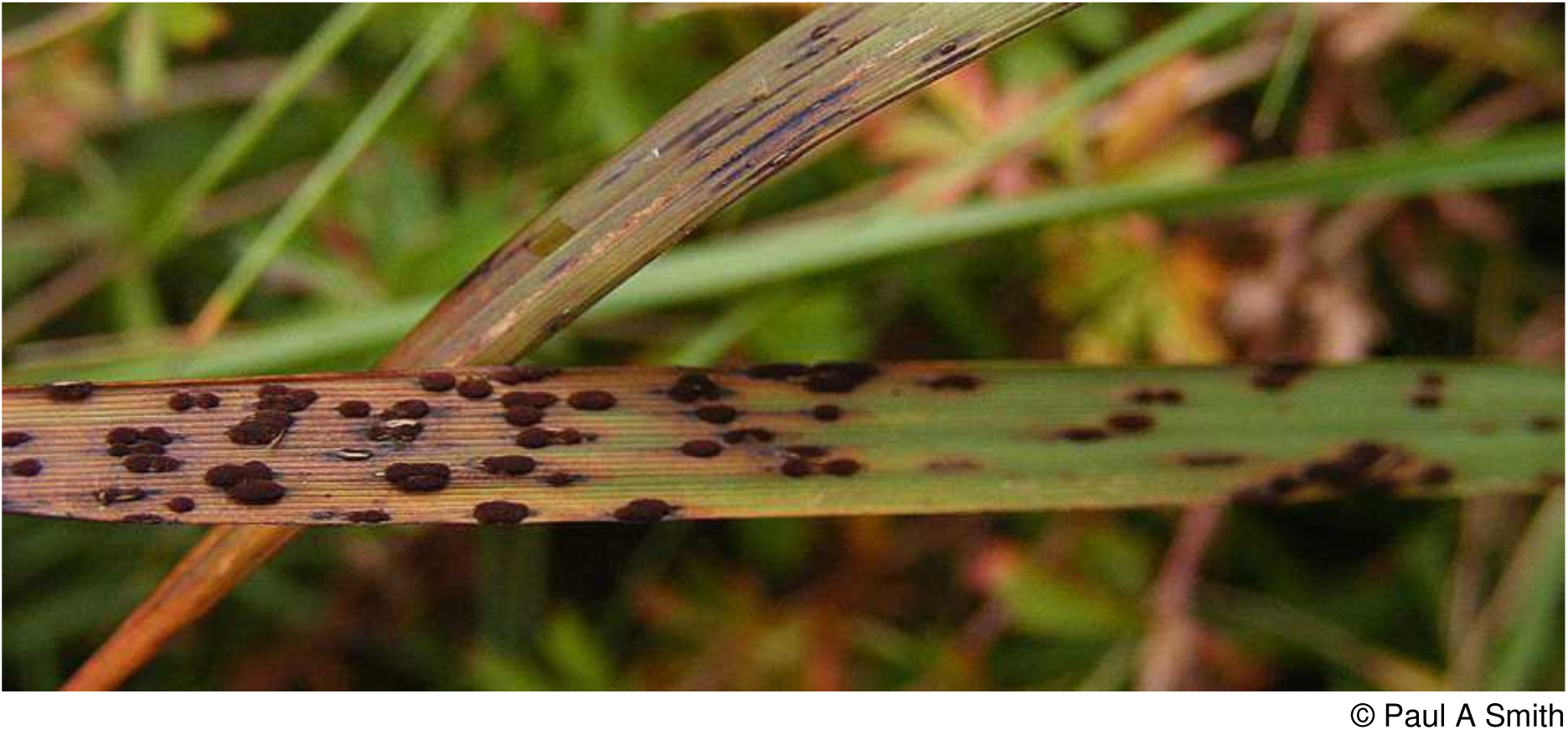

## Appendix 3

### Pursuing an Interest in Rusts

Should this Red Data List and Census Catalogue spark an interest in this fascinating group of fungi the following sources may prove to be helpful.

### Introducing the rusts

David Ingram and Noel Robertson provide one of the most readable introductions to this group of fungi, in chapter 9 of HarperCollins New Naturalist Volume 84 *Plant Disease*, published in 1999, entitled “A Treacherous, Mutable Tribe - The Rusts”.

In 1981 Iowa State University Press, Ames, Iowa published Larry J. Littlefield’s *Biology of the Plant Rusts - An Introduction*. This book is an excellent introduction to plant rusts for those people who are not familiar at all with these fungi. The book deals with the history of plant rusts, life cycles and taxonomy as well as host-pathogen relationships. The later chapters cover spore dispersal mechanisms and the control of rust diseases. The book was aimed at undergraduate level and although dated is not too technical and is suitable for the beginner.

### Identification Guides

There is no up to date, well-illustrated guide to British rust fungi in print.

Martin B. Ellis and J. Pamela Ellis in their magnificent *Microfungi on Land Plants. An Identification Handbook* published as a new enlarged edition by The Richmond Publishing Co. Ltd. in 1997, whilst expensive, is an invaluable guide to all fungi found on land (and many aquatic plants) in Britain. The fungi are described and many illustrated by each host genus. So if you know the host you have immediately eliminated most species from consideration.

The most comprehensive monograph on British rusts was written by Malcolm Wilson and Douglas M. Henderson. Entitled *British Rust Fungi*, its last edition was produced in 1966 and in consequence is somewhat out of date. It has, however, been returned to print by Cambridge University Press and is available in both hard and soft back. Each species is described and many spore forms are illustrated by black and white line drawings. There is a key to genera but it is entirely reliant on teliospores so in practice it is of limited use. Instead, provided the host has been identified, recourse must be had to the index and comparison made between the descriptions of the rust species if more than one rust occurs on a particular host. Despite this it is a book worth acquiring. The easiest work to acquire, and the most practical for the beginner to use for basic naming is Douglas M. Henderson’s *The Rust Fungi of the British Isles. A Guide to Identification by their Host Plants*. Published in 2004 by the British Mycological Society, Kew this lists the host plants by their families and the rusts they support. It can be downloaded as a PDF file from the web on www.aber.ac.uk/waxcap/links/. Couple this publication with Malcolm Storey’s many fine illustrations on www.bioimages.org.uk and good progress is possible.

The only truly modern book with clear descriptions and colour illustrations is A.J. Termorshuizen and C.A. Swertz’s *Roesten van Nederland* (*Dutch Rust Fungi*). Published in 2011 and sadly now out of print, the text is Dutch, followed by an English translation in a slightly smaller font. It covers 260 species, and takes in most if not all of the Welsh rust mycota.

### Featured Species

These two species have been selected to illustrate the value of producing a Rust Fungus Red Data List for Wales. The first has highlighted the need in some sites to manage them for a normally little-regarded host species whilst the second identifies better targeted survey approaches to elucidate the true distribution of a little-recorded rust fungus.

#### Marsh Pennywort Rust (*Puccinia hydrocotyles*) Endangered in Wales

Distinctive and easily spotted on marsh pennywort leaves by its cinnamon-brown pustulate uredosori scattered over the upper surface of the leaves, many thousands of host colonies have been searched, but only a handful have been found with this rust in Wales. All sites support other notable wetland species and unless alerted to the presence and importance of this rust, such a common species as marsh pennywort could be sacrificed to create new open water or allowed to be colonised by wet woodland.

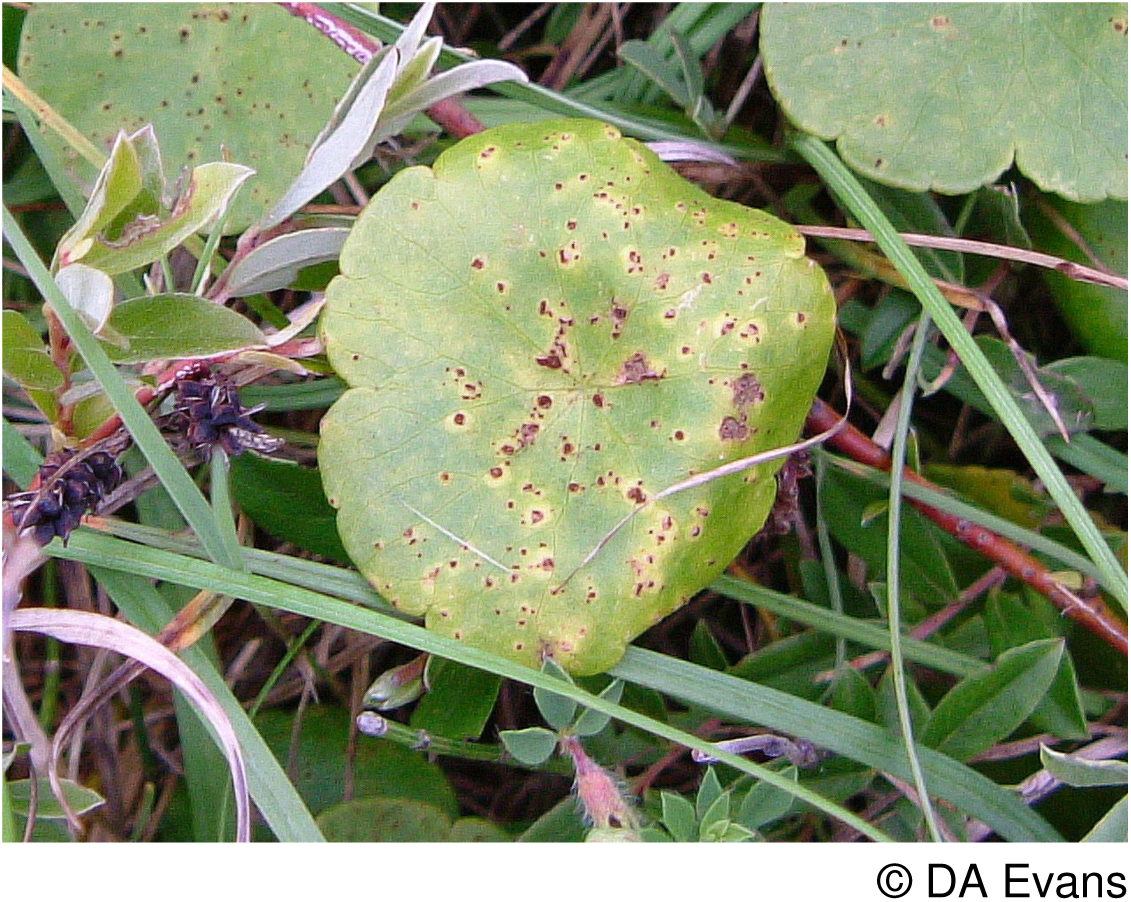

#### A rust fungus (*Puccinia dioicae*) on Small Sedges and Thistles Endangered in Wales

This rust alternates between the diminutive dioecious sedge or slightly larger common yellow sedge and meadow or marsh thistle. Finding the rust on the small leaves of sedges, whilst standing in usually a very wet peaty flush, is difficult. Targeting marsh thistles in close proximity to the sedges has located new records.

There is a single record on meadow thistle from the North of England. Mid and South Wales support internationally notable populations of meadow thistle and should clearly be searched for this rust.

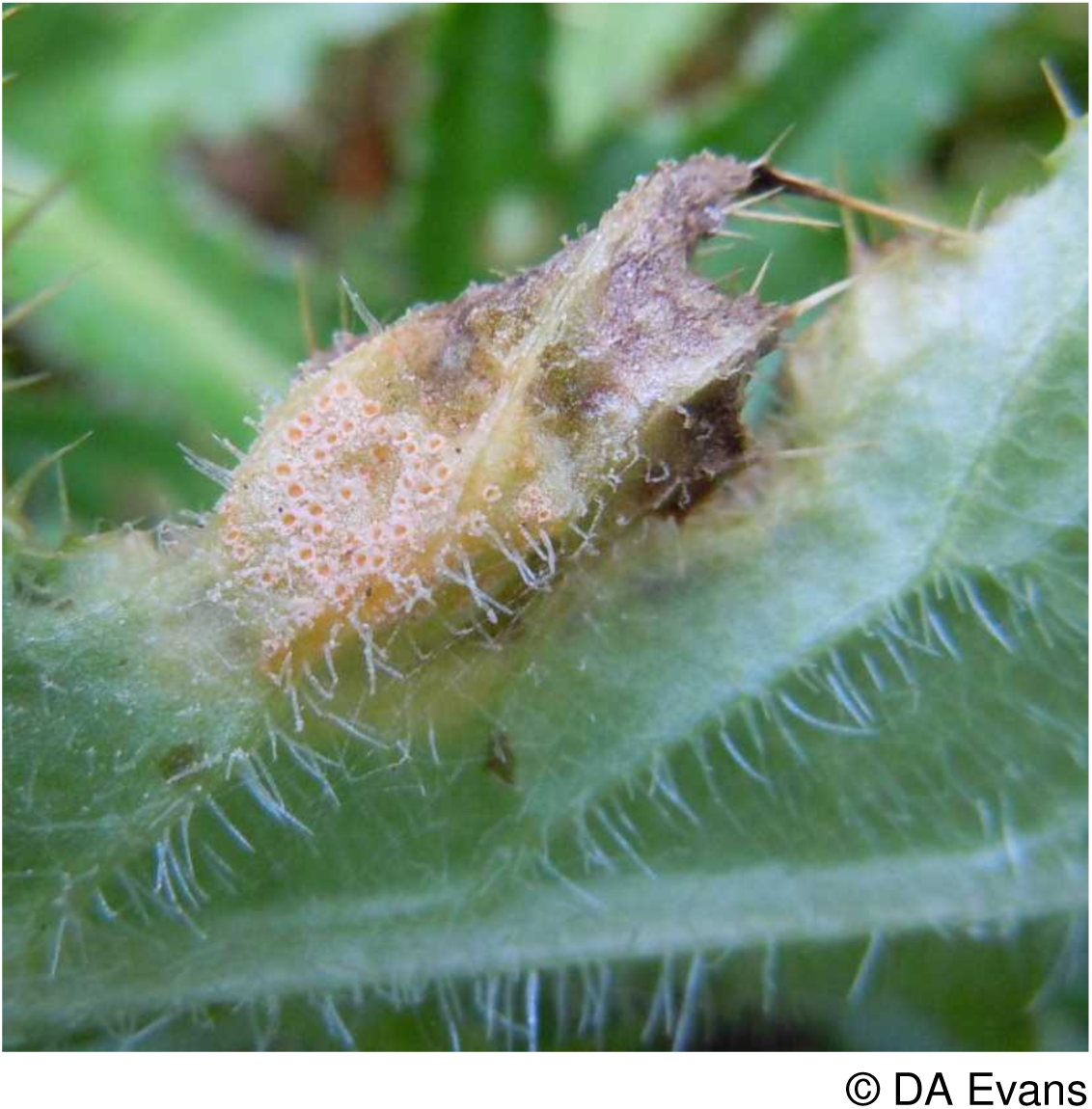

Rear cover image:

*Triphragmium filipendulae* on dropwort, Great Orme, Caernarvonshire © DA Evans

## References

Aron, C. (2005). Fungi of Northwest Wales. Privately published by the author.

Bosanquet, S.D.S. & Dines, T.D. (2011). A Bryophyte Red Data List for Wales. Plantlife, Salisbury.

Bowen, H. (2000). The Flora of Dorset. Pisces Publications, Newbury.

Braithwaite, M.E., Ellis, R.W. & Preston, C.D. (2006). Change in the British Flora 1987-2004. Botanical Society of the British Isles, London.

Burdon, J.J., Ericson, L. & Muller, W.J. (1995). Temporal and spatial changes in a metapopulation of the rust pathogen *Triphragmium ulmariae* and its host, *Filipendula ulmaria*. Journal of Ecology, 83, 979–989.

Cheffings, C.M. & Farrell, L. (eds.) (2005). The Vascular Plant Red Data List for Great Britain. Joint Nature Conservation Committee, Peterborough.

Clark, M.C. (ed.) (1980). A Fungus Flora of Warwickshire. British Mycological Society, London.

Dines, T.D. (2008). A Vascular Plant Red Data List for Wales. Plantlife International, Salisbury.

Ellis, R.G. (1983) Flowering plants of Wales. National Museum of Wales, Cardiff.

Evans, S., Henrici, A. & Ing, B (2006). Red Data List of Threatened Fungi. http://www.britmycolsoc.org.uk/mycology/conservation/red-data-list/

Hawksworth, D.L. (2001). The magnitude of fungal diversity: the 1.5 million species estimate revisited. Mycological Research 105 (12), 1422–1432.

Helfer, S. (1993). Rust Fungi - A Conservationist’s Dilemma. In Pegler, D.N., Boddy, L., Ing, B. & Kirk, P.M. (eds.). Fungi of Europe: Investigation, Recording and Conservation, pp.287–294, Royal Botanic Gardens, Kew.

Helfer, S., Berndt, R., Denchev, C. M., Moricca, S., Scheuer, C., Scholler, M. & St.Quinton, J. M. (2011). A call for a renewed and pan-European strategic effort on the taxonomy of rust fungi (*Uredinales*). Mycologia Balcanica, 8, 78–80.

Henderson, D.M. (2004). The Rust Fungi of the British Isles. A Guide to Identification by their Host Plants. British Mycological Society, Kew.

Ingram, D.S. (1999). Biodiversity, plant pathogens and conservation. Plant Pathology, 48, 433–442.

Ingram, D.S. & Robertson, N. (1999). *Plant Disease A Natural History*. HarperCollins, London

IUCN (2001). IUCN Red List Categories and Criteria. Version 3.1. IUCN, Gland, Switzerland.

IUCN (2003). Guidelines for the Application of IUCN Red Data List Criteria at Regional Levels: Version 3.0. IUCN Species Survival Commission. IUCN, Gland, Switzerland & Cambridge, UK.

Legon, N.W. & Henrici, A. with Roberts, P.M., Spooner, B.M. & Watling, R. (2005). Checklist of the British and Irish Basidiomycota. Royal Botanic Gardens, Kew.

Liu, M. & Hambleton, S. (2013). Laying the foundation for a taxonomic review of *Puccinia coronata* s.l. in a phylogenetic context. Mycological Progress, 12, 63–89.

Moricca, S., Ragazzi, A. & Assente, G. (2006). Biocontrol of rust fungi by *Cladosporium tenuissimum*. In Pei, M. H. & McCracken, A.R. (eds.). Rust Disease of Willow and Poplar pp. 213–229. CABI Publishing, Wallingford, Oxfordshire.

Pei, M.H. & McCracken, A.R. (eds.) (2006). Rust Diseases of Willow and Poplar. CABI Publishing, Wallingford, Oxfordshire.

Preece, T.F. (1995). Shropshire Fungus Group A List of Uredinales for the County. Manuscript.

Preece, T.F. (2004). James Edward Vize, Curate and Vicar of Forden, Montgomeryshire, 1869-1910, Mycologist. Montgomeryshire Collections, 92, 117–125.

Preston, C.D., Pearman, D.A. & Dines, T.D. (2002). New Atlas of the British and Irish Flora. Oxford University Press, Oxford.

Scottish Montane Willow Research Group (2005). Biodiversity: taxonomy, genetics and ecology of sub-arctic willow scrub. Royal Botanic Gardens, Edinburgh.

Smith, J.A., Blanchette, R.A. and Newcombe, G. (2004). Molecular and morphological characterization of willow rust fungus, *Melampsora epitea* from Arctic and Temperate hosts in North America. Mycologia, 96(6), 1330–1338.

Stace, C. (2010). New Flora of the British Isles (3rd ed.). Cambridge University Press, Cambridge.

Stevens, D.P., Smith, S.L.N., Blackstock, T.H., Bosanquet, S.D.S. & Stevens, J.P. (2010). Grasslands of Wales. University of Wales Press, Cardiff.

Stringer, R.N., Smith, P.A. & Davies, R.H. (2011). Infection of *Arum maculatum* by the rust *Puccinia sessilis* at Carmel NNR, South Wales. Field Mycology, 12 (1), 5–8.

Trueman, I., Morton, A. & Wainwright, M. (1995). The Flora of Montgomeryshire. The Montgomeryshire Field Society and the Montgomeryshire Wildlife Trust, Welshpool.

Vize, J.E. (1882). The Parish of Forden. II.-Natural History. II.-Botany. Montgomeryshire Collections, 15, 165–178.

Watson, H.C. (1883). Topographical Botany. 2nd ed. B. Quaritch, London.

Wilson, M. & Henderson, D.M. (1966). British Rust Fungi. Cambridge University Press, Cambridge.

Woods, R.G. (1993). Flora of Radnorshire. Bentham-Moxon Trust, Kew.

Woods, R.G. (2010). A Lichen Red Data List for Wales. Plantlife, Salisbury.

Zadoks, J.C. (2005). Sea Lavender, Rust and Mildew. A perennial Pathosystem in the Netherlands. Wageningen Academic Publishers. The Netherlands.

